# Aging disrupts spatiotemporal coordination in the cycling murine ovary

**DOI:** 10.1101/2024.12.15.628550

**Authors:** Tammy C.T. Lan, Alison Kochersberger, Ruth Raichur, Sophia Szady, Hien Tran, Radiana Simeonova, Ashley Helmuth, Andrew Minagar, Maria José Orozco Fuentes, Vipin Kumar, Giovanni Marrero, Irving Barrera, Sarah Mangiameli, Alex K. Shalek, Pardis C. Sabeti, Fei Chen, David S. Fischer, Jennifer L. Garrison, Hattie Chung

## Abstract

Throughout the female reproductive lifespan, the ovary completes hundreds of cycles of follicle development, ovulation, and tissue regeneration^1–3^. These processes rely on the precisely coordinated intricate multicellular interactions across time and space^4^. How aging disrupts these interactions, leading to an overall decline in reproductive and endocrine functions, remains understudied. To understand the multicellular dynamics that underlie ovarian function and their changes with age, here we use Slide-seq, a near-cellular spatial transcriptomics method, to profile 22 mouse ovaries across the reproductive cycle and chronological age, representing 610,620 near-cellular spots across 69 spatial transcriptomic profiles^5,6^. We develop a segmentation analysis to identify spatial niches that capture different states of folliculogenesis from static snapshots *in situ*, allowing us to examine the multicellular dynamics of 358 oocytes, 668 follicles, and 236 corpora lutea. We find that aging disrupts both the spatial organization and temporal coordination of folliculogenesis before the cessation of cycling, which may contribute to the dysregulation of hormone production and signaling. These disruptions are marked by altered immune cell dynamics, inflammatory signaling, and global tissue disorganization that impair the cyclic remodeling required for ovarian function. Our findings reveal how multicellular niches orchestrate ovarian function and demonstrate how age-related breakdown of tissue organization across time and space precedes reproductive decline.

## Main Text

The ovary is a dynamic organ comprising diverse cell types that undergo continuous remodeling to regulate reproductive and endocrine function throughout the female lifespan. During the female reproductive period, ovarian cells undergo a repetitive cycle of phenotypic switching to collectively orchestrate oocyte maturation, ovulation, and hormone production throughout the estrous or menstrual cycle^4,7^. This process of "homeostatic cycling" occurs approximately 150–650 times in mice and 300–400 times in humans over the course of the female reproductive lifespan^1,2^. During chronological aging, overall reproductive function declines and culminates in the cessation of cycling. Here, we investigate the molecular interactions underlying this process, including the progressive depletion of the ovarian reserve, the decline in the precision of cellular interactions, and the increase in tissue-wide inflammation and fibrosis^8–13^.

Central to ovarian function are spatially organized dynamic multicellular structures – primarily follicles, which develop in distinct stages before transforming into corpora lutea (CL) after ovulation. Folliculogenesis begins with a primordial follicle consisting of a dormant oocyte encased by flattened granulosa cells (GCs). Activation, the awakening of the follicle from its dormant phase, triggers GCs to adopt a cuboidal shape and begin proliferation, forming the preantral follicle where they coordinate morphological and functional changes to support oocyte growth. As the follicle transitions to the antral stage, a fluid-filled cavity (antrum) forms, and the follicle becomes gonadotropin-dependent for growth and begins producing steroid hormones^14,15^. Spatial complexity further increases as theca cells create an outer layer and respond to gonadotropin hormones by producing steroids^16,17^. Ultimately, less than 1% of follicles recruited to grow successfully progress to ovulation^18^, while the remainder undergo atresia, a form of apoptosis^18^. Though atresia claims most follicles during their maturation process – which spans approximately two weeks in mice and more than a year in humans – the dominant surviving follicle releases a mature oocyte at ovulation. After oocyte ejection, the remaining follicular cells transform into the CL, a transient hormone-producing structure essential for pregnancy and critical in regulating the complex cycle of ovulation^15,19^. This creates an extraordinarily complex tissue environment where, at any given time, dozens (mice) or thousands (humans) of follicles at different developmental stages share the same physical space with multiple CLs^20,21^. The molecular mechanisms coordinating this intricate asynchronous spatiotemporal organization remain unclear, highlighting the need for spatially resolved profiling to better understand the dynamics of these cellular ecosystems.

Previous studies of ovarian aging focused primarily on follicle depletion and decline in oocyte-intrinsic properties such as aneuploidy^22,23^. However, mounting evidence suggests that dysfunction across the entire ovarian tissue environment driven by the repeated cyclic remodeling of folliculogenesis and ovulation contributes significantly to reproductive aging through dysregulation of these multicellular ecosystems^10,12,23–25^, particularly through disruptions in granulosa cell signaling^26–28^, extracellular matrix remodeling^29–32^, and immune infiltration^12,33–35^, underscoring the need to investigate how multicellular dysregulation drives ovarian aging. Single-cell RNA sequencing is a powerful tool to profile cellular heterogeneity and uncover novel regulatory mechanisms, but it faces limitations in studying ovarian tissue. Droplet-based single-cell methods are limited in their ability to profile large cells like oocytes, which exceed 100 µm in diameter resulting in capture of early stage oocytes in low abundance making analysis challenging^36–39^ and do not resolve interacting cell types in the follicular niche, while coarse spatial transcriptomics approaches lack the resolution to characterize follicular architecture^25,40^ and often rely on predetermined gene panels that constrain discovery of novel regulatory patterns within and across niches^41^.

Here, we present a spatiotemporal atlas of the naturally cycling C57BL/6J murine ovary across chronological aging at near-single-cell resolution using Slide-seq^5,6^. Our study comprises *n*=610,620 spots at 10 µm resolution spanning 69 spatial transcriptomic profiles from 22 biological samples collected during the estrus and metestrus stages of the estrous cycle, across three age groups. To characterize the cooperative nature of critical multicellular structures in the ovary, we developed a segmentation pipeline that identified 358 primordial and growing oocytes, 668 follicles of which 301 enclosed oocytes, and 236 CLs, enabling detailed analysis of *in situ* molecular signatures. Comparing profiles between young and aged mice revealed that aging disrupts the temporal dynamics of folliculogenesis and corpus luteum function during the estrous cycle. This disruption coincides with increased inflammatory signatures and a decline in tissue organization, which may compound ovarian dysfunction. Our work highlights how multicellular niches cooperate within the ovary and proposes that temporal dysregulation of these niches contributes to the decline in reproductive and endocrine function with age.

### Spatial transcriptomics reveals multicellular organization of the cycling and aging ovary

We used Slide-seq to generate 69 spatial transcriptomics profiles from 22 mice harvested during ovulatory (estrus) and post-ovulatory (metestrus) stages across young (10-12 weeks), reproductively middle aged (36-40 weeks), and reproductively old (52-54 weeks) mice^42^ (**Fig. 1a**; **Extended Data Fig. 1a, Extended Tables 1, 2**; **Methods**). This design enabled us to examine the post-ovulatory tissue-wide remodeling across aging. Unsupervised clustering and *post hoc* annotation of 601,831 high-quality 10 µm spot transcriptomes identified all major ovarian cell types, including oocytes (*Gdf9*), multiple GC subtypes, two luteal cell populations (defined by *Parm1* and *Akr1c18*), theca cells (*Col1a2*), epithelial cells (*Epcam*), fibroblasts (*Pdgfra*), smooth muscle cells (*Tagln*), immune (*Lyz2*), endothelial (*Flt1*), stromal cells (*Aldh1a1*), and others (**Fig. 1b,c**; **Extended Data Fig. 1b**; **Extended Table 3**; **Supplementary File 1**). Cell types were found across all ages, with an age-associated decrease in oocytes and an increase in immune cells (**Fig. 1c**). To validate our Slide-seq annotations with truly single-nucleus transcriptomics, we also performed snRNA-seq on whole ovaries, which revealed a consistent set of cell populations, but with a notable loss of oocytes and cumulus cells, highlighting the advantage of our combined approach for capturing a more complete ovarian cellular atlas (**Extended Data Fig. 1c,d**).

**Figure 1.**
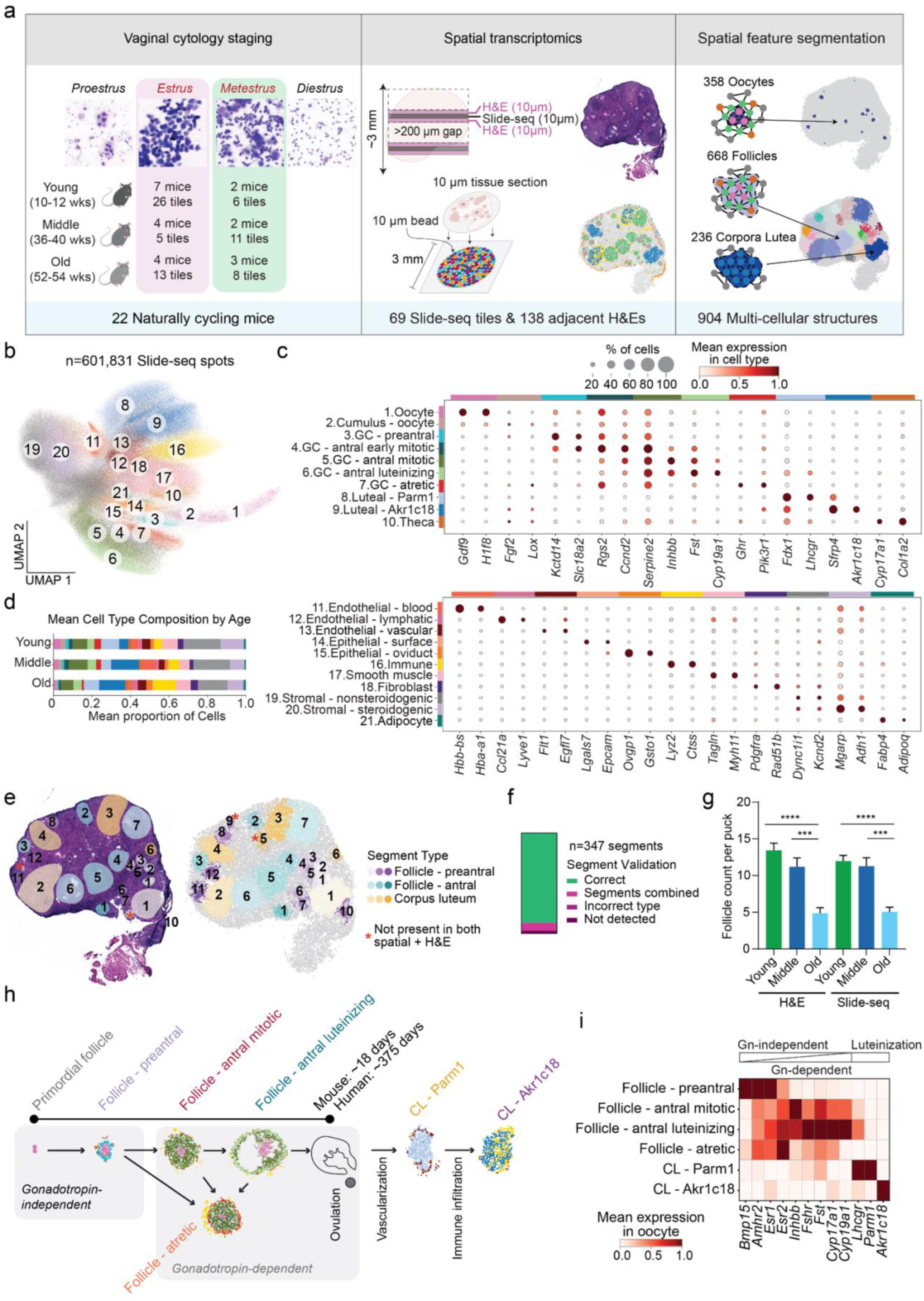
A spatiotemporal map of the cycling and aging ovary identifies spatial niches driving folliculogenesis. **a.** Overview of experimental design. Mice were staged via vaginal cytology and harvested from estrus and metestrus stages (*n*=22; young 10-12 weeks, middle 36-40 weeks, old 52-54 weeks). Multiple non-adjacent sections (10 µm thickness) were profiled using Slide-seq from each ovary with a minimum gap of 200 µm to avoid resampling the same follicle or CL structure. Adjacent sections were taken for histological analysis. After clustering and cell type annotation, multicellular segments were defined with an automated segmentation pipeline. **b.** UMAP embedding of RNA profiles (dots) captured by Slide-seq spots (n=610,620; each at 10 µm resolution) across 69 Slide-seq pucks (**Methods**). Spots colored by post hoc annotation (color legend). “GC”: granulosa cell. **c.** Mean expression (log normalized counts, dot color) and proportion of expressing cells (dot size) of marker genes (columns) used for annotating cell type clusters (rows). **d.** Mean cell type composition across young (*n*=32), middle (*n*=16), and old (*n*=21) pucks. **e.** Representative H&E sections and corresponding spatial segment maps showing annotated preantral follicles (purple), antral follicles (green-blue), and corpora lutea (yellow). Asterisks denote segment structures only present in either H&E or Slide-seq sections due to sampling, rather than an error of the segmentation. **f.** Quantification of segmentation accuracy across *n*=347 segments where an adjacent H&E section was present to validate with, the majority (*n*=311) were correct. Error types included segments that were grouped with another object in the pipeline, segments where the incorrect type was assigned, and segments that were not detected by the pipeline. Cohen’s kappa=0.85. **g.** Mean follicle counts of adjacent H&E sections and Slide-seq segmentation across young (*n*=32), middle (*n*=16), and old (*n*=21) pucks. Error bars: SEM (\*\*\**P*<0.001, \*\*\*\**P*<0.0001 two-tailed t-test; all other bar groups, not significant). **h.** Schematic of folliculogenesis and CL formation. GCs form a single layer around a dormant oocyte in primordial follicles. Activation triggers GCs to proliferate, forming the preantral follicle. The follicle then transitions to the gonadotropin-dependent antral stage during which a fluid-filled cavity forms the antrum. Theca cells are recruited to create an outer layer, and GCs differentiate into multiple subtypes. Follicle atresia can occur during any stage of folliculogenesis. After ovulation, GCs luteinize into luteal cells that form the CL, which is subsequently cleared from the tissue. **i.** Mean expression (log normalized counts and variance-scaled) of key hormone receptors and synthesis genes (columns) across segment types (rows), grouped by gonadotropin-independent, gonadotropin-dependent, and luteinization stages. Segment counts: preantral (*n*=238), antral mitotic (*n*=191), antral luteinizing (*n*=43), atretic (*n*=181), CL - Parm1 (*n*=87), CL - Akr1c18 (*n*=147); *n*=22 mice across all ages.

To analyze anatomically related groups of spots representing meaningful tissue units, we developed a segmentation pipeline that identified oocytes, follicles, and corpora lutea. As large oocytes often exceed 70 µm in diameter^43^ thereby spanning multiple Slide-seq spots, we clustered spatially adjacent putative oocyte spots into segments to create composite oocyte profiles for further analysis (**Methods**). As oocytes in primordial follicles typically comprise roughly 2 spots, we identified these through a separate pipeline that segmented at least 2 neighboring oocyte spots (**Methods**). The identity of primordial oocytes was further validated by expression of additional marker genes such as *Sohlh1* and *Nobox*^44^ (**Extended Data Fig. 1e**).

Larger structures, follicles and CL, were identified via unsupervised clustering of the spatial neighborhood graph and classified based on their GC and luteal composition. These segments revealed distinct stages of follicular development, including preantral (n=238), antral mitotic (n=191), antral luteinizing (n=43), and atretic (n=181) follicles, as well as two types of CLs classified as *Parm*1 (n=87) and *Akr1c18* (n=147) (**Extended Data Fig. 1g,h**; **Methods**). We validated the segmentation pipeline by manually annotating adjacent H&E-stained sections (**Fig. 1e-g**; **Extended Data Fig. 1i**; **Supplementary Files 2,3**; **Methods**). Follicle classifications captured the progression of GC function that drives folliculogenesis (**Fig. 1h**), supported by stage-specific expression of key hormone receptors, including the anti-Müllerian hormone receptor *Amhr2* in gonadotropin-independent preantral follicles^45^ and *Fshr* and *Lhcgr* in gonadotropin-dependent antral follicles^46,47^ (**Fig. 1i**). Additionally, to provide orthogonal support for identifying atretic follicles beyond transcriptional signatures, we examined the spatial distribution of immune cells within follicles^37^. We observed that atretic follicles show increased immune infiltration toward the follicle center, whereas healthy follicles exhibit minimal immune presence (**Extended Data Fig. j**; *P*<0.001 antral *vs* atretic follicles, Tukey’s test), which aligns with prior observations that immune cells infiltrate dying follicles for tissue clearance^12^. Altogether, these results demonstrate that our segmentation pipeline captures the multicellular architecture and functional organization of the ovary.

### Oocytes follow coordinated transcriptional programs during follicular development

Despite the traditional characterization of oocytes as transcriptionally silent maternal RNA storage units^48^, recent studies have demonstrated active transcriptional regulation during their maturation^43,49,50^. From follicle activation, primordial oocytes grow approximately 7-fold in size, progressing from prophase I to arrested metaphase II over the course of 18 days in mice^51^. We examined distinct transcriptional patterns of oocytes in their native tissue context, defined by the enclosing follicle typ^36,52–54^. *Sohlh1* was elevated in primordial oocytes^55^, *Figla* and *Kit* in preantral follicles, and *Gdf9* and *H1f8* in antral follicles (**Fig. 2a-c**). We next identified distinct gene programs that differ across oocyte maturation via nonnegative matrix factorization, revealing five programs that significantly differed in their utilization across oocytes at different follicle stages (NMF; **Fig. 2d**; **Methods**; **Extended Data Fig. 2a-c**). Oocyte program 2, characterized by zona pellucida glycoproteins *Zp2* and *Zp3*, was upregulated in primordial and preantral oocytes, suggesting upregulation of extracellular matrix components necessary for follicular assembly immediately after primordial follicle activation^56,57^. Oocyte program 4, enriched in F-box proteins (*Fbxw19, Fbxw21, Fbxw24*) that are critical components of the SCF ubiquitin E3 ligase complex controlling cell cycle progression^58,59^ and *Bmp15*, emerged in mature antral oocytes, reflecting the rapid shifts necessary for meiotic progression. Oocyte program 0 which contains the regulator of meiosis resumption, *Bcl2l10*, and early embryonic development genes *Btg4* and *Tcl1* was elevated in mature follicles preparing to respond to the gonadotropin surge (**Fig. 2e**; **Extended Data Fig. 1b**; *P*<0.001 preantral vs antral; t-test)^60–62^. Oocytes within atretic follicles showed divergent expression of regulators of maturation processes, with lowered expression of *Padi6* and upregulation of oocyte program 0 (**Fig. 2a,e**; **Extended Data Fig. 2b**; *P*<0.05 antral vs atretic; t-test). Altogether, our findings reveal that oocytes undergo dynamic transcriptional regulation throughout folliculogenesis, switching from processes required to maintain oocyte dormancy to proliferative programs upon follicle activation.

**Figure 2.**
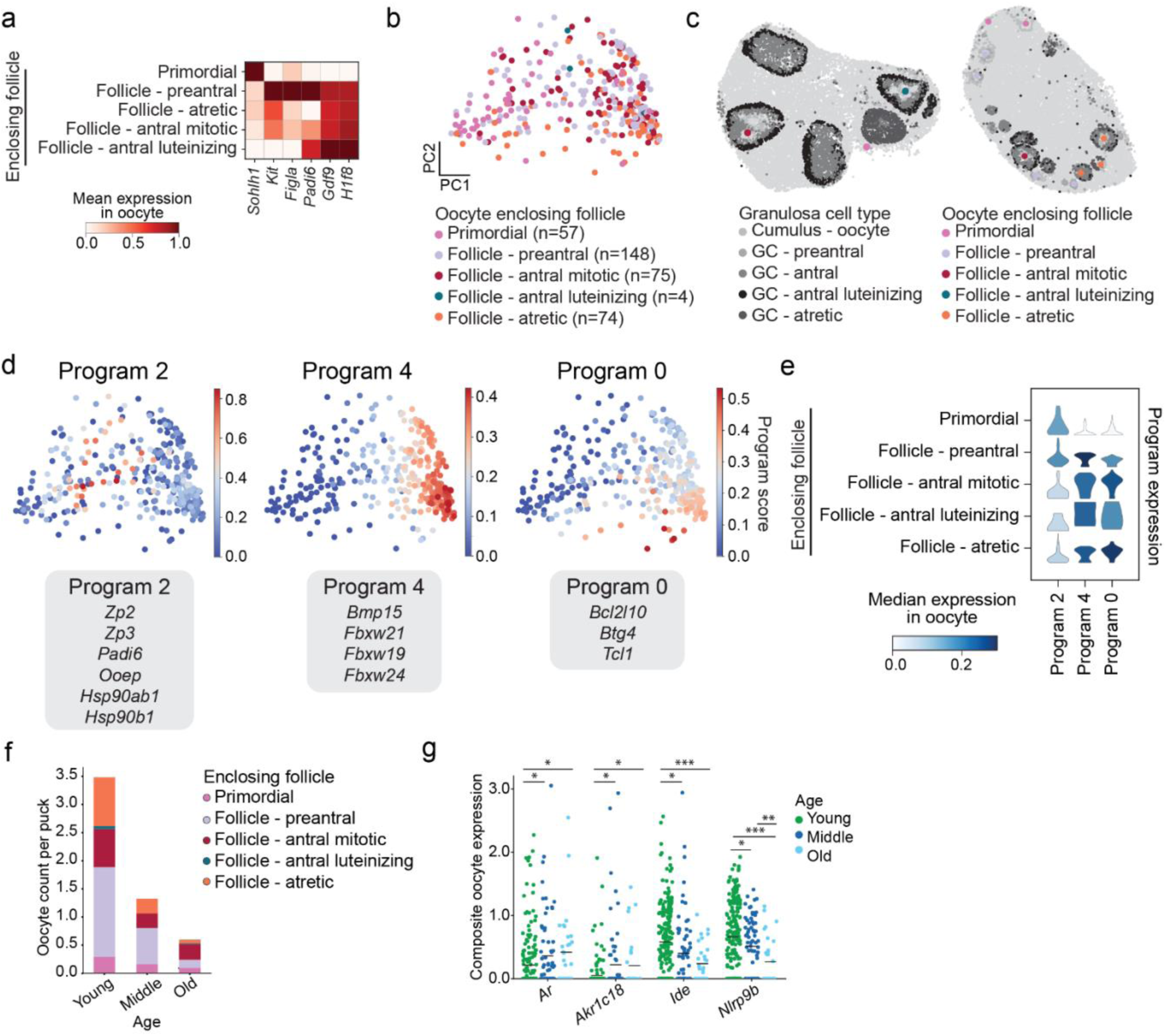
Oocyte transcriptional dynamics across folliculogenesis and age. **a.** Mean expression (log normalized counts and variance-scaled) of oocyte marker genes (columns) grouped by the enclosing follicle at different stages of folliculogenesis (rows); *n*=358 oocytes across 22 mice, all age groups. Oocyte counts per follicle type in **Extended Data Table 4**. **b.** Principal component (PC) embedding of RNA profiles (dot) indicated by its enclosing follicle stage (color); *n=*358 oocytes across 22 mice, all age groups. **c.** Example spatial plots of two pucks showing oocytes (colored dots) in their tissue context depicting the stage of its enclosing follicle (color), with surrounding GC spots (gray). **d.** Expression (color) of gene programs identified in oocytes (*n*=358) via non-negative matrix factorization. Top genes contributing to each oocyte gene program, gray text box. **e.** Violin plot of median expression (color) and distribution (kernel density) of oocyte gene programs (columns) across stages of folliculogenesis (rows); *n*=358 oocytes across 22 mice. Oocyte counts per follicle type and age in **Extended Data Table 4**. **f.** Average composite oocyte count per puck (y axis) by enclosing follicle type (color) across age groups (x axis). Oocyte frequencies per age group: young (*n*=229), middle (*n*=89), and old (*n*=40). **g.** Differentially expressed genes of oocytes across age. Select genes (*y* axis) showing significant age-associated changes in oocytes (dots), with colors indicating age; \**P*<0.05, \*\**P*<0.005, \*\*\**P*<0.0005, two-tailed t-test, Benjamini-Hochberg correction. Differentially expressed genes were first identified between young and old non-primordial oocytes, then tested pairwise across young, middle, and old to identify monotonic trends. Non-primordial oocyte counts by age in **Extended Data Table 4**.

### Aging alters regulators of oocyte maturation

Aging leads to both the depletion and functional decline of maturing oocytes, with reduced oocyte quality observed across follicle stages^63,64^. Consistent with this, we observed a sharp decrease in the average oocyte numbers in each puck across all follicle types with age (**Fig. 2f**). Differential expression analysis between maturing oocytes of young and old animals revealed multiple genes that change with age (**Extended Data Fig. 2d**; **Methods**). In particular, expression of the nuclear androgen receptor *Ar* and hydroxysteroid dehydrogenase *Akr1c18,* known to metabolize progesterone, increased in aging oocytes (**Fig. 2g**; **Extended Data Fig. 2d**). These genes are known to be involved in oocyte maturation and meiosis^65–67^, and their upregulation may reflect an adaptive shift by oocytes to changing hormone levels in the aging ovarian environment^68,69^. Furthermore, we observed decreased expression of the insulin-degrading enzyme *Ide*, which attenuates insulin signaling in healthy folliculogenesis^70,71^. Insulin signaling regulates the final stages of oocyte maturation, although high insulin levels can reduce oocyte quality, underscoring the need for balanced signaling^71,72^. Interestingly, aging oocytes show downregulation of the *Nlrp9b* family genes (*Nlrp9a*/*b/c*), which normally increases during late maturation in young oocytes^73^ (**Extended Data Fig. 2e**). These NOD-like receptor homologs localize to the subcortical maternal complex and are thought to regulate the oocyte-to-embryo transition^74,75^, though their role is under debate^75^. Furthermore, these changes are consistent across follicle types enclosing the oocytes, suggesting that these trends are not driven by changes in the follicular composition with age (**Extended Data Fig. 2f**). Together, our results indicate that aging may disrupt the integration of external signals that guide oocyte maturation, suggesting a potential mechanistic basis for age-related decline in oocyte quality.

### Molecular programs define spatially organized follicle lifecycles in young ovaries

Oocyte maturation occurs in concert with dynamic remodeling of surrounding granulosa cells. To characterize this organization, we systematically compared GC states across hundreds of follicles in young ovaries by quantifying cell states as a function of scaled radial distance from the follicle center, revealing extensive intra-follicle heterogeneity (**Fig. 3a, b**; **Extended Data Fig. 3a**; **Methods**). Cell cycle analysis and diffusion pseudotime (DPT) analysis of GC trajectories toward their two terminal fates, luteinization or atresia, revealed that proliferating GCs are enriched near the follicle center, whereas differentiated GCs with higher pseudotime localize to the periphery, consistent with literature that proximity to the theca-granulosa boundary influences specialization (**Fig. 3c,d; Extended Data Fig. 3b,c**; **Methods**). Immunostaining confirmed these patterns, with *Ki67* marking proliferative GCs near the center and *Cyp19a1* labeling mature GCs at the periphery (**Fig. 3e,f**). To further identify spatially organized transcriptional patterns, we applied consensus non-negative matrix factorization (cNMF)^76^ to identify co-expressed GC gene expression programs (GEPs) and mapped their spatial distribution (**Extended Data Fig. 3d,e,f**). In non-atretic follicles, a GC maturation program (GEP2; *Inhbb*, *Fst*) localized centrally, while a luteinization program (GEP1; *Hsd3b1*, *Lhcgr*, *Mro*) was enriched at the periphery (**Fig. 3g**). This compartmentalization was disrupted in atretic follicles, which showed globally reduced expression of both programs. Altogether, our findings indicate that granulosa cells maintain a spatial division of labor balancing proliferation and differentiation during follicle development, which is lost with atresia.

**Figure 3.**
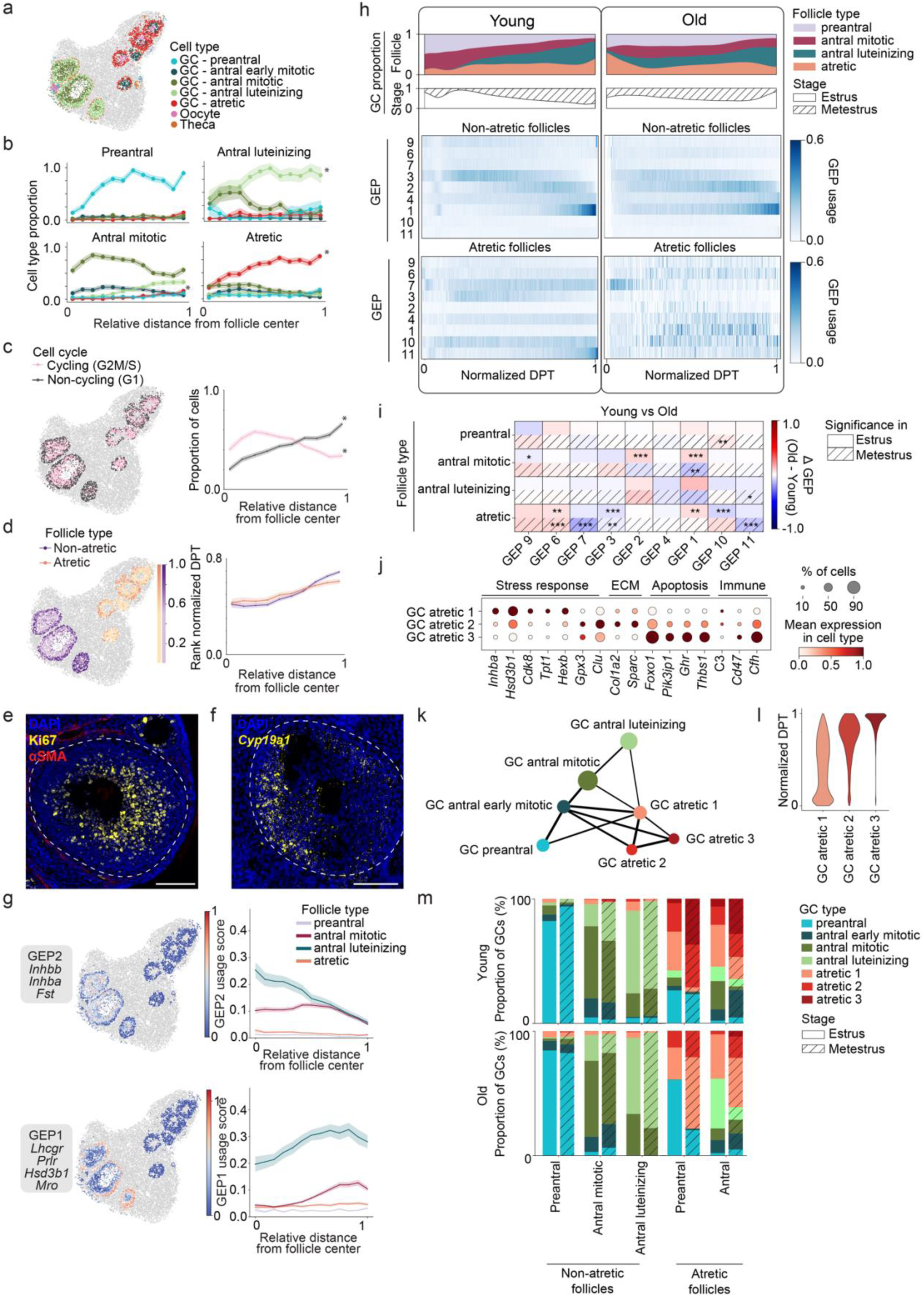
Spatiotemporal coordination of granulosa cells within the follicular niche. **a.** Spatial plot of an example puck showing follicular cell types (colors), including preantral, antral early mitotic, antral mitotic, antral luteinizing, and atretic granulosa cell (GC) subtypes, and oocyte and theca. **b.** Radial distribution of GC subtypes (colored lines, y axis) across follicle type (panels). Lines show the mean proportion of each subtype across 10 normalized radial bins as a function of the relative radial distance from the center to the edge (x axis; 0 = center, 1 = edge); shading indicates standard error of the mean (SEM) across follicles. Monotonically significant radial trends are indicated (\**P*<0.05, Spearman correlation coefficient); *n*=653 total follicles across 22 mice, all age groups. Follicle counts by type in **Extended Data Table 4**. **c.** GC cell cycle states. Left: spatial example showing cycling (G2M/S, pink) and non-cycling (G1, gray) GCs. Right: Mean proportion of cycling vs non-cycling GCs (y axis) across radial bins; shading indicates SEM across follicles. Monotonically significant radial trends are indicated (\**P*<0.05, Spearman correlation coefficient; *n*=668 follicles). **d.** Diffusion pseudotime (DPT) of GCs. Left: Spatial example showing DPT values (colorbar) across two follicle fates, non-atretic (purple) and atretic (orange). Right: Mean DPT of GCs (y axis) across radial bins; shading indicates SEM across follicles. Monotonically significant radial trends are indicated (\**P*<0.05, Spearman correlation coefficient); *n*=653 total follicles across 22 mice, all age groups. Follicle counts by type in **Extended Data Table 4**. **e.** Ki67 (yellow) and ɑSMA (red) immunofluorescence in an antral follicle of a young ovary. Scale bar, 100 µm. **f.** *Cyp19a1* (yellow) mRNA detected by fluorescence in situ hybridization in a young ovary. Scale bar, 100 µm. **g.** GC gene programs (GEPs) identified by cNMF. Left: Spatial expression of two distinct GEPs (colorbar), FSH regulation (left) and luteinization (right), with top contributing genes in gray. Right: Mean program usage score (y axis) across radial bins (x axis) for preantral (purple), antral mitotic (maroon), antral luteinizing (teal), and atretic (orange) follicles (shading indicates SEM); *n*=653 total follicles across 22 mice, all age groups. Follicle counts by type in **Extended Data Table 4**. **h.** Dynamics of GC GEPs across pseudotime (DPT), follicle type, and estrous stage in young (*n*=9) and old (*n*=7) mice. Top: kernel density distribution of follicle types (preantral, antral mitotic, antral luteinizing, and atretic) across normalized pseudotime (gaussian_kde); estrous stage composition below. Heatmaps show mean normalized GEP usage (color) for GC spots in non-atretic (middle panels) and atretic (bottom panels) follicles across pseudotime (*n*=500 bins). **i.** Age-associated differences in GC GEP usage. Each cell shows the mean difference in usage (ΔGEP = [Age= Old]) per GEP (columns) between Old (*n*=7) and Young (*n*=9) mice within follicular types (rows), in estrus (upper half) and metestrus (lower half). Red = higher usage in Old; blue = lower. Statistical significance from linear models (GEP ∼ Age; **Methods)**; Stars indicate Benjamini-Hochberg FDR-adjusted significance (*q<0.05, **q<0.01, ***q<0.001). Follicle counts by age, stage, and type in **Extended Data Table 4**. **j.** Mean expression (log normalized counts, dot color) and proportion of expressing cells (dot size) of marker genes (columns) used for annotating atretic GC subtypes (rows); *n*=17,175 spots across 22 mice, all age groups. Spot counts by GC type in **Extended Data Table 4**. **k.** Partition-based Graph Abstraction (PAGA) of GC types. **l.** Distribution of normalized pseudotime (DPT; x axis) for spots within three distinct atretic granulosa cell clusters. **k.** Mean GC type composition of follicles by age, stage, and follicle type; *n*=105,390 spots across 22 mice, all age groups. Follicle counts by age, stage, and type in **Extended Data Table 4**.

### Aging disrupts the temporal trajectory of GC differentiation

We next examined how GCs utilize GEPs across their differentiation trajectory, spatial context, estrous stage, and reproductive age by ordering usage profiles by diffusion pseudotime to better visualize usage patterns (**Fig. 3h**; **Extended Data Fig. 3e,g,h**). In young, non-atretic follicles, GEP usage followed the expected sequence of folliculogenesis defined by canonical markers. For instance, GCs in preantral follicles during estrus exhibited high usage of an early development program GEP9 (*Wt1, Wnt4*) but low activity of proliferative GEP3 (*Top2a*, *Mki67*) and follicular dominance GEPs 2 and 4 (*Inhbb*, *Fst*; *Esr2*, *Greb1l*). By contrast, GCs in luteinizing follicles showed increased activity of the follicular dominance GEPs as well as the luteinization GEP1 (*Lhcgr*, *Hsd3b1*), particularly during metestrus, while atretic follicles expressed apoptosis-associated GEPs 10, 11 (*Thbs1*, *Pik3ip1*; *Ghr, Ptprd*) (**Fig. 3h, Extended Data Fig. 3f,g**). In aged ovaries, this orderly transcriptional trajectory was significantly altered. In non-atretic follicles, expression of early development (GEP9) decreased, follicular dominance and luteinization (GEP2, 1) lost stage specificity, and an apoptosis-associated program (GEP10) was elevated (**Fig. 3h**,**i**; **Extended Data Fig. 3g,h**). Notably, GCs in atretic follicles were the most affected by aging, with downregulation of canonical apoptosis-associated (GEP10, 11) and proliferation (GEP3) modules but aberrant upregulation of the extracellular matrix (ECM) remodeling program (GEP6; *Col1a2*, *Lama2*) typically confined to preantral follicles in young animals. These patterns suggest a shift from regulated cell to dysregulated remodeling, blurring developmental and degenerative boundaries in aged follicles. Together, these results indicate that aging disrupts the defined temporal sequence of GC differentiation and impairs the regulation of follicle atresia.

### Aging alters the progression of follicular atresia

Given the pronounced transcriptional alterations in atretic follicles, we next conducted a deeper investigation of atretic GC heterogeneity and progression. Subclustering atretic GCs revealed three states (**Fig. 3j**): state 1, marked by metabolic reprogramming (*Cdk8*, *Hexb*, *Hsd3b1*, *Tpt1*)^77–81;^ state 2, enriched for oxidative stress response and matrix remodeling (*Clu*, *Gpx3, Sparc*, *Col1a2*)^82–84;^ and state 3, defined by apoptotic and immune-modulatory signatures (*Foxo1*, *Ghr*, *Pik3ip1*, *Cfh*, *Cd47*)^85–89^. While analysis from transcriptomic snapshots cannot confirm developmental origins and trajectories, we performed partition-based graph abstraction (PAGA) to infer how different GC states relate to each other and found that transitions to atretic GC may arise from multiple non-atretic GC states, most strongly from antral early GCs, consistent with the high rates of follicular atresia before dominant follicle selection^90,91^ (**Fig. 3k**). Furthermore, pseudotime analysis suggested a trajectory from state 1 to state 3 (**Fig. 3l**), supported by the observation that states 1 and 2 were enriched in immature follicles, whereas state 3 predominated in mature and fully atretic follicles. These distributions suggest a progression from early metabolic reprogramming through stress and ECM remodeling to terminal apoptosis, transitioning from early atretic subtypes in estrus to late apoptotic populations in metestrus (**Fig. 3m**).

Aging disrupted atretic progression. Across all follicle types, we observed a decrease in terminal apoptotic and immuno-modulatory GC atretic 3 in aged ovaries (**Fig. 3m**). This failure to transition to the terminal atretic state was accompanied by upregulation of genes associated with the complement system, reduced levels of the phagocytosis inhibitor “don’t-eat-me-signal” *Cd47* (**Extended Data Fig. 3i**). Together, these findings suggest that effective atresia requires a coordinated transition to an apoptotic, immune-active state that promotes follicular clearance, and that aging impairs this process, potentially leading to unresolved atresia, chronic inflammation, and accumulation of follicular debris in the aged ovary.

### Multicellular dynamics and interactions shaping folliculogenesis progression

Folliculogenesis progresses through coordinated interactions among multiple cell types, extending beyond cell-autonomous granulosa programs. To resolve this higher-order organization (**Fig. 4a**), we arranged follicle segments from reproductively young mice along a computationally inferred folliculogenesis trajectory (**Fig. 4b,c**). Pairwise distributional distances between segments, calculated using the Wasserstein distance^92^ (**Fig. 4b,c**), enabled clustering of follicles and corpora lutea by multicellular similarity, visualized in a UMAP embedding (**Fig. 4d**; **Extended Data Fig. 4a**; **Methods**). Diffusion pseudotime analysis recovered the expected transitions from preantral to antral or atretic follicles (**Fig. 4e**), accompanied by gradual shifts in cell type composition (**Extended Data Fig. 4b**). To examine how compositional changes were reflected in cell-cell communication, we applied MultiNicheNet^93^ to infer receptor-ligand interactions among oocytes, granulosa, theca, and immune cells across preantral, antral, and atretic follicles, treating each follicle as a sample (**Methods**; **Extended Data Fig. 4c**). Follicle stage-specific interactions included *Gdf9*-*Fxyd6* between oocytes and GCs in preantral follicles, *Fst*-*Bmpr1b* between GCs and theca cells in antral follicles, and *Apoa4*-*Lrp1* within GCs in atretic follicles. Receptor-ligand interaction scores varied smoothly across the folliculogenesis trajectory (**Fig. 4f**; **Methods**). The candidate atretic interaction, *Apoa4-Lrp1*, suggests follicle metabolic reprogramming, as *Lrp1* (low-density lipoprotein receptor-related protein 1) regulates lipid transport and glucose metabolism^94,95^, while *Apoa4* promotes lipid metabolism, and their interaction facilitates glucose uptake^96^. Notably, *LRP1* has been genetically linked to polycystic ovarian syndrome (PCOS), a disorder characterized by metabolic and hormonal dysfunction^97^.

**Figure 4.**
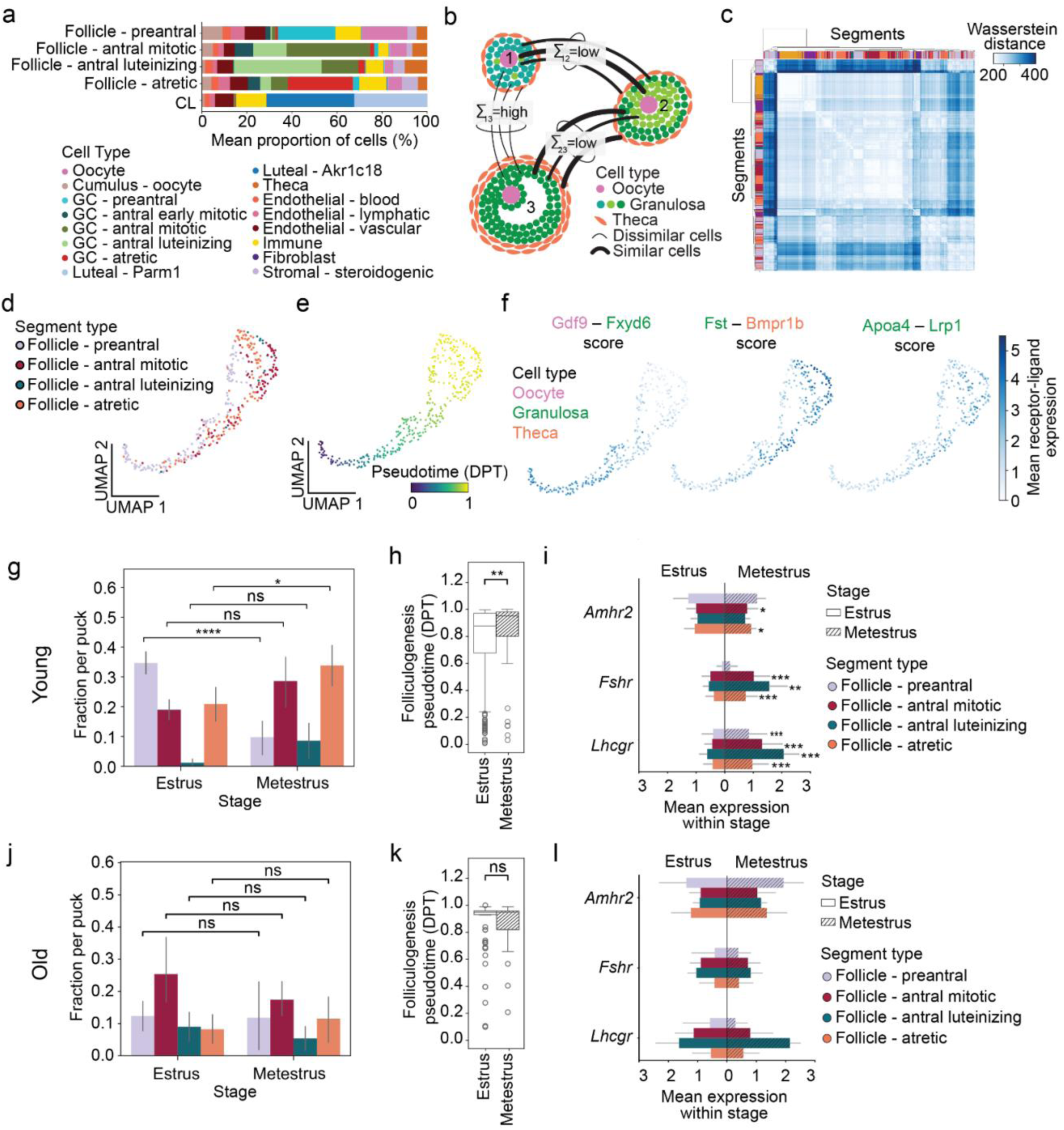
Aging disrupts the temporal variation in folliculogenesis. **a.** Mean cell type composition of preantral (*n*=238), antral mitotic (*n*=191), atretic (*n*=181), and antral luteinizing (*n*=43) follicles, and CL (*n*=234). **b.** Schematic of optimal transport inference of multicellular transitions, where pairs of follicles are aligned by minimizing their distributional distance. Transcriptionally similar pairings (e.g. GC to GC) incurs low cost, while whereas dissimilar pairings (e.g. theca to GC) incur high cost. **c.** Pairwise transition matrix of segments from young mice (left; rows, columns) showing transport costs (blue) organized by unsupervised hierarchical clustering (dendrogram, axes) and annotated by segment type (color). **d,e.** Interpolated folliculogenesis trajectory in young mice (*m*=382 follicles across 8 mice). UMAP of follicles (dots) based on transport distances, colored by segment type (**d**) and diffusion pseudotime (DPT, colorscale) (**e**). **f.** UMAPs of follicles (dots) showing mean ligand-receptor expression scores (color) for significant interactions identified by MultiNicheNet (**Methods**). Ligand (panel title, left) and receptor (panel title, right) components are colored by the expressing cell type; follicle counts: preantral *n*=144, antral *n*=120, atretic *n*=160. **g-l.** Follicle characteristics in young (*n*=9) (**g-i**) and old (*n*=7) mice (**j-l**). **g,j.** Fraction of follicle types per puck (y axis) across estrus and metestrus (x axis; \*\**P*<0.005, ns, not significant; two-tailed t test). Puck counts: 23 in young estrus, 6 in young metestrus, 12 in old estrus, 6 in old metestrus. **h,k.** Boxplots of follicle DPT (y axis) by stage (x axis, bar shading) (\*\**P*<0.005, ns, not significant; two-tailed t-test). Center line indicates median, box bounds represent first and third quartiles, whiskers span from each quartile to the minimum or the maximum (1.5 interquartile range below 25% or above 75% quartiles). Follicle counts: young estrus *n*=281, young metestrus *n*=66, old estrus *n*=58, old metestrus *n*=23. **i,l.** Mean expression of hormone receptor genes (x axis) across follicles grouped by type (color) and stage (bar shading). Error bars: SEM (\**P*<0.05, \*\*\**P*<0.01, two-tailed t-test; all others ns). Follicle counts by age, stage, and type in **Extended Data Table 4**.

### Temporal coupling in follicle activation, maturation, and hormone sensing

Although all follicles in the ovary experience the same systemic hormones, they progress asynchronously through distinct developmental states across the estrous cycle. We therefore analyzed how follicle populations of different stages coexist across the cycle in reproductively young mice. In metestrus, the proportion of preantral follicles was significantly lower compared to estrus (**Fig. 4g**; *P*=0.001, two-tailed t-test). This shift likely reflects an increase in antral and atretic follicles at metestrus, as preantral follicle numbers remain largely stable across the cycle, while larger follicles expand^98–101^. Furthermore, this change is likely not attributed to differential preantral atresia, since death rates of small follicles remain constant^99,101^. To assess how follicle maturation progresses across the cycle, we compared antral follicle features between estrus and metestrus. The hormonal dynamics preceding estrus are critical: surges in follicle-stimulating hormone (FSH) and luteinizing hormone (LH) trigger competition among antral follicles for dominance, driving them toward either ovulation or atresia such that mature antral follicles are typically depleted by estrus^100–104^. Consistent with this, antral mitotic follicles were significantly smaller during estrus than metestrus (**Extended Data Fig. 4d;** *P*=0.001, two-tailed t-test) and exhibited lower pseudotime values, indicating a less mature state (**Fig. 4h**; **Extended Data Fig. 4e;** *P*=0.002, two-tailed t-test). Functionally, antral follicles at estrus showed reduced expression of the gonadotropin receptors *Lhcgr* and *Fshr* (**Fig. 4i**; \**P***<**0.01, \*\**P*<0.001, two-tailed t-test), suggesting that they are in an immature, non-hormone-responsive state, possibly due to their recent transition from the gonadotropin-independent preantral state^14^.

### Aging disrupts homeostatic temporal coupling

The coordinated timing of follicle maturation and hormone responsiveness in young mice prompted us to ask how this coupling changes with age. A hallmark of ovarian aging is the onset of irregular cycling, as observed during perimenopause in humans, indicating disrupted temporal regulation of endocrine signaling and ovarian function^105,106^. Although the reproductively aged mice in our study were cycling, their molecular profiles revealed a loss of temporal coordination observed in young mice. Unlike young mice, old mice showed no stage-dependent shifts in preantral follicle composition and antral follicle size (**Fig. 4j**; *P*=0.942, two-tailed t-test; **Extended Data Fig. 4f**). Projecting follicles from old mice onto the reference trajectory of young mice revealed a distortion in pseudotime distribution, showing no significant distinction between estrus and metestrus (**Fig. 4k**; *P*=0.677, two-tailed t-test; **Extended Data Fig. 4e,g**; **Methods**). Consistently, *Fshr* and *Lhcgr* expression within antral follicles did not vary across stages in old mice (**Fig. 4l**). In young ovaries, gonadotropins synchronize follicle cohorts to ensure coordinated progression toward ovulation or atresia, yielding immature antral follicles at estrus. By contrast, old ovaries had larger, more mature follicles at estrus, consistent with disrupted follicle development. While reduced follicle numbers in old ovaries limit statistical power, the consistent absence of stage-dependent differences in follicle composition, pseudotime, and hormone receptor expression suggests that temporal regulation of folliculogenesis becomes uncoupled from the estrous cycle with age, preceding acyclicity and reproductive senescence.

### Temporal dysregulation of corpus luteum clearance in aged ovaries

We next examined whether the corpus luteum dynamics are also altered with age. Of note, we observed two spatially segregated luteal cell populations with opposing progesterone regulatory programs (**Fig. 5a, Extended Data Fig. 5a**)^107^. Marked by *Parm1* and *Akr1c18*, these populations correspond to known luteal subtypes previously attributed to granulosa and theca origins^108,109^ (**Extended Data Fig. 5b)**. However, our data suggest that these populations reflect a temporal phenotypic switch during the CL lifecycle (**Extended Data Fig. 5c**)^37^. *Parm1*-enriched CLs (‘CL – Parm1’) express progesterone synthesis genes, contain vascular endothelial cells, are enriched at estrus, and consist of smaller histologically^110,111^, features consistent with newly formed, hormonally active structures (**Extended Data Fig. 5a-e**)^19,37,107,112,113^. In contrast, *Akr1c18*-enriched CLs (‘CL – Akr1c18’) express the regression marker prostaglandin F receptor *Ptgfr*, infiltrated by immune cells, more frequent at metestrus, and contain histologically larger cells^110,111^, features indicating a regression state (**Extended Data Fig. 5a-f**). In aged ovaries, while we find the transcriptional identities of the luteal cells are maintained (**Extended Data Fig. 5g**), *Akr1c18* CLs abnormally accumulated at estrus (**Fig. 5b**, *P*=0.006, two-tailed t-test). Because timely CL clearance and accompanying drop in progesterone are required to initiate the next cycle^19,104,113^, persistence of these degenerating CLs likely disrupts progesterone homeostasis, potentially impairing normal cycle progression^4,14^.

**Figure 5.**
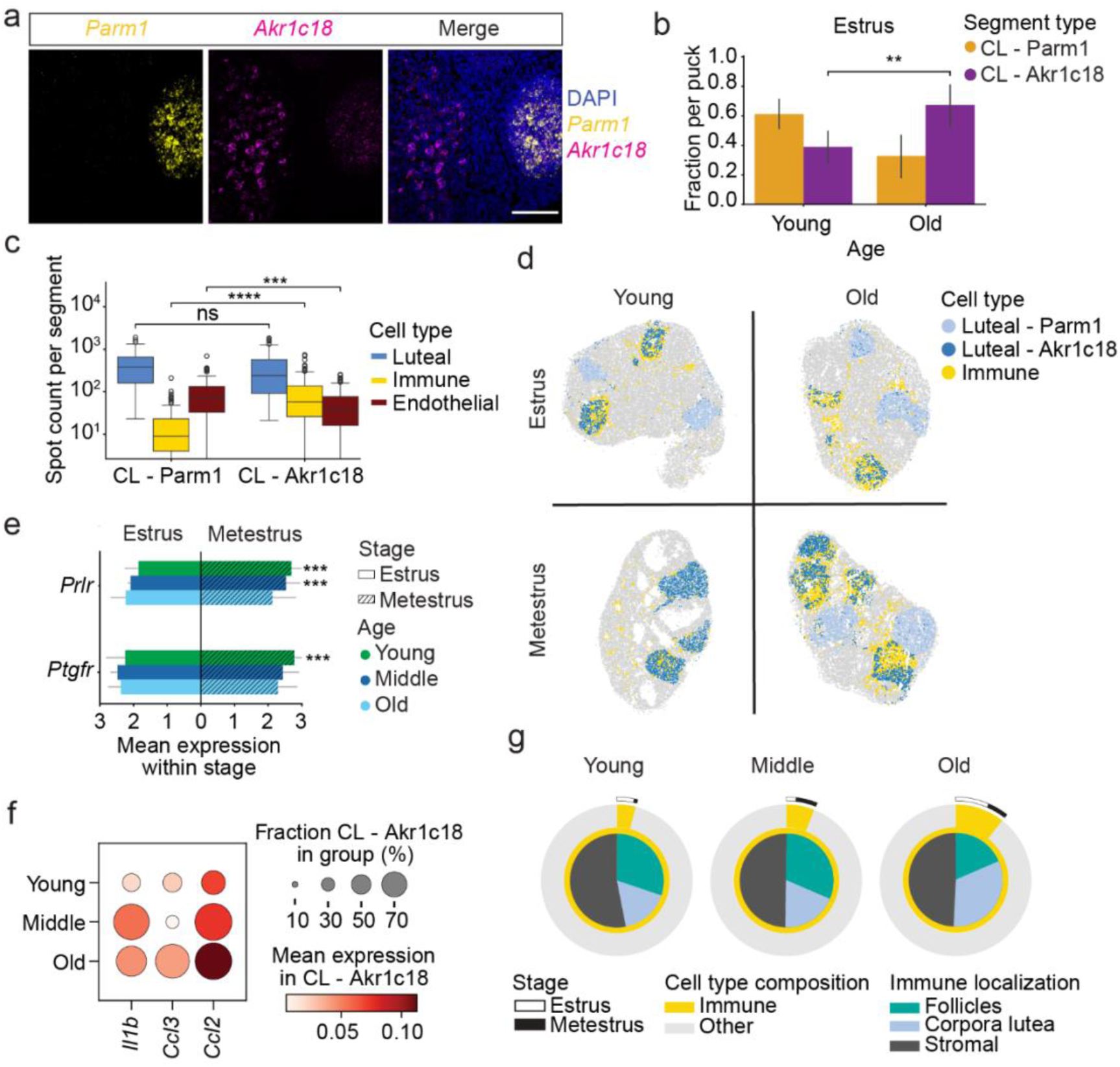
Aging disrupts corpus luteum clearance. **a.** *Parm1* (yellow) and *Akr1c18* (teal) mRNA labeled by fluorescence in situ hybridization in a young ovary. Scale bar, 100 µm. **b.** Fraction of each CL type per puck (y axis) during estrus, grouped by age (x axis). Error bars: standard deviation across pucks (young *n*=23, old *n*=12). **c.** Cell type composition of CL subtypes. Boxplots show spot counts (y axis, log_10_ scale) for luteal (early and late combined), immune, and vascular cells (color) across CL - Parm1 and CL - Akr1c18 (\*\*\**P*<0.001, \*\*\*\**P*<0.0001; two-tailed t-test). Center line indicates median, box bounds represent first and third quartiles, whiskers span from each quartile to the minimum or the maximum (1.5 interquartile range below 25% or above 75% quartiles); *n*=234 corpora lutea across 22 mice, all age groups. CL type counts in **Extended Data Table 4**. **d.** Spatial maps from representative pucks across stage (estrus, metestrus) and age (young, old) showing luteal (blues) and immune (yellow) cells; all other cell types, gray. **e.** Mean expression of CL regression receptor genes (x axis) in Akr1c18 CLs across young (*n*=9), middle (*n*=6), and old (*n=7*) mice (color) by stage (shading). Error bars: SEM (\**P*<0.05, \*\*\**P*<0.01, two-tailed t-test; all others ns). Akr1c18 CL counts by age and stage in **Extended Data Table 4**. **f.** Mean expression and proportion of CL - *Akr1c18* segments (dot size) expressing selected genes (x-axis) involved in CL clearance and immune cell recruitment, across young (*n*=9), middle (*n*=6) and old (*n=7*) mice (rows). CL - Akr1c18 counts by age in **Extended Data Table 4**. **g.** Immune population changes with age. Outer ring: fraction of immune (yellow) vs. non-immune (gray) spots across estrus (white) and metestrus (black). Inner ring: distribution of immune spots across spatial compartments for young (*n*=31), middle (*n*=16), and old (*n*=21) pucks.

Investigating the molecular aberrations in persistent CLs revealed dysregulated inflammatory and luteolysis signaling with age. Under homeostatic conditions, immune cells accumulate in *Akr1c18* CLs at metestrus to facilitate tissue clearance during luteolysis^12^ (**Fig. 5c**). In aged ovaries, immune infiltration was significantly elevated in *Akr1c18* CLs at metestrus (**Fig. 5d**; **Extended Data Fig. 5h**; *P*<0.005, two-tailed t-test). Furthermore, aged *Akr1c18* CLs failed to upregulate the prolactin receptor *Prlr* and the prostaglandin receptor *Ptgfr* (**Fig. 5e**), which normally trigger luteolysis, but instead showed elevated expression of the immune-recruiting chemokine *Ccl2*, along with pro-inflammatory cytokines *Il1b* and *Ccl3*^33,38,39^ (**Fig. 5f**; **Extended Data Fig.** 5i). Across ovarian compartments, immune populations increased most prominently within CLs, implicating defective CL clearance as a major driver of inflammaging (**Fig. 5g**). Altogether, our findings indicate that aged CLs exhibit dysregulation of prolactin- and prostaglandin- mediated regression, leading to persistent CLs that sustain inflammation and contribute to endocrine decline.

### Global immune compositional changes define age-associated inflammation

As impaired tissue clearance and chronic inflammation are major hallmarks of ovarian aging^10,33^, we sought to closely examine the functional and spatial dynamics of immune populations. We identified four macrophage subsets, along with dendritic cells, mast cells, plasma cells, and T cells with distinct spatial localization within tissues (**Fig. 6a,b**; **Methods**). The macrophage subsets included canonical phagocytic (Macrophage 1: *Cd36*, *Mmp12*), antigen-presenting (Macrophage 2: *Cd74*, *H2-Aa*, *S100a4/6*), efferocytic/repair (Macrophage 3: *Mrc1*, *Stab1*, *Mertk*), and metabolically active (Macrophage 4: *Acsbg1*, *Adh1*) populations. Phagocytic and antigen-presenting macrophages (Macrophages 1 and 2) were the most abundant and expanded significantly with age (**Fig. 6b**). Spatial mapping revealed that the most significant accumulation in immune cells occurred in preantral and atretic follicles and CLs (**Fig. 6c**). Across all ages, phagocytic macrophages (Macrophage 1) localized to the periphery, while antigen-presenting macrophages (Macrophage 2) were enriched at the core, suggesting their key role in inflammaging (**Fig. 6d,e**). This pattern was accentuated in aged ovaries, which showed a general trend towards greater immune infiltration towards the segment center (**Fig. 6d**). Immunofluorescence of a regressing CL revealed CD11b+ macrophages and CD3+ T cells at the CL center, supporting the likely role of antigen presentation in CL clearance (**Fig. 6f**). Although low in abundance, T cells expand in CLs of aged ovaries, which may be the functional site of prior reports of cytotoxic and double negative T cell accumulation in the aging ovary^39^. Altogether, spatial profiling highlights age-related immune infiltration within ovarian structures, with phagocytic and antigen-presenting macrophages showing particularly strong enrichment in atretic and luteinizing follicles.

**Figure 6.**
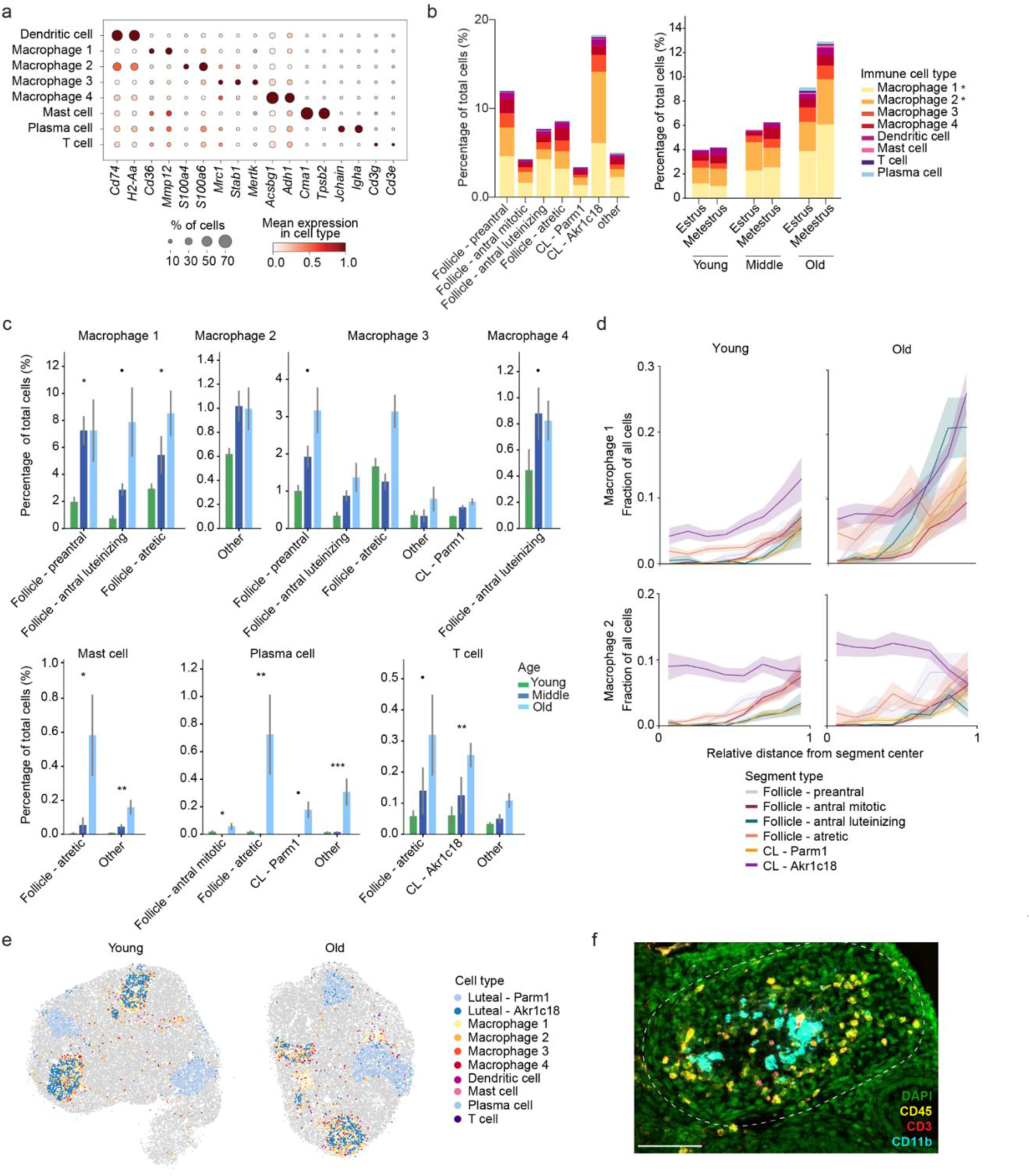
Age-associated changes in immune composition. **a.** Mean expression (log normalized counts, color) and proportion of expressing spots (size) of marker genes (columns) of immune cell types (rows); spot counts in **Extended Data Table 4**. **b.** Spatial and temporal distribution of immune cell types. Left: Percentage of immune cell types across spatial structures. Right: Percentage across stage and age. (\**P*<0.05, age-associated changes identified by scCODA). Follicle type and puck counts by age and stage in **Extended Data Table 4**. **c.** Immune populations significantly altered with age within spatial structures. Error bars: SEM across young (*n*=9), middle (*n*=6), and old (*n=7*) mice. (\**P*<0.05, \*\*\**P*<0.01, two-tailed t-test; all others, ns). Follicle type by age in **Extended Data Table 4**. **d.** Distribution of macrophage subtypes (panels) by mean proportion in segment type (colored lines, y axis) in young (*n*=9) and old (*n=7*) mice across radial bins (x axis; 0 = center, 1 = edge). Shading indicates standard error of the mean (SEM); frequencies of segment types by age in **Extended Data Table 4**. **e.** Representative spatial maps of immune cell types in young and old animals. **f.** Immunofluorescence labeling of immune cells in a regressing CL (dotted line) from a young ovary. Scale bar, 100 µm.

A hallmark of ovarian aging is the accumulation of multinucleated giant cells (MNGCs), which reflect chronic inflammation and are associated with fibrosis^11,33,114,115^. MNGCs, visible as clusters of fused cells on H&E stains, increased markedly with age (**Fig. 7a,b**; **Extended Data Fig. 6a**). To define their molecular features, we performed image registration between adjacent pairs of H&E images and Slide-seq pucks using anatomical landmarks to identify immune spots located within *vs* outside of MNGCs (**Fig. 7c**; **Extended Data Fig. 6b; Methods**). Macrophages (four distinct subsets), particularly the phagocytic subtype (Macrophage 1), were the predominant immune population enriched within MNGCs (**Fig. 7d**). Differential expression analysis revealed distinct transcriptional shifts in MNGC-associated macrophages (**Extended Data Fig. 6c**). Phagocytic macrophages (Macrophage 1) upregulated ribosomal and proliferative genes (*Rps24*, *Cdk11c*) while downregulating chemokines (*Ccl6*, *Ccl9*) and *Spp1*, suggesting reduced immune signaling (**Fig. 7e**). Antigen-presenting macrophages (Macrophage 2) upregulated complement (*C3*), fusion-related genes (*Tagln*, *Acta2*), and *Itgae* (CD103), consistent with a tissue-resident state. Efferocytic/repair macrophages (Macrophage 3) upregulated clearance and lysosomal genes (*Gpnmb*, *Mfge8*, *Ctsd*) but reduced *Adgre1* (F4/80). Metabolically active macrophages (Macrophage 4) expressed higher levels of inflammatory and lipid regulators (*Pla2g7*, *Lyz2*, *Lgals3*). Together, these findings show that macrophages within MNGCs adopt specialized transcriptional programs distinct from those in the surrounding stroma, reflecting coordinated roles in cell fusion, complement activation, efferocytosis, and metabolic remodeling.

**Figure 7.**
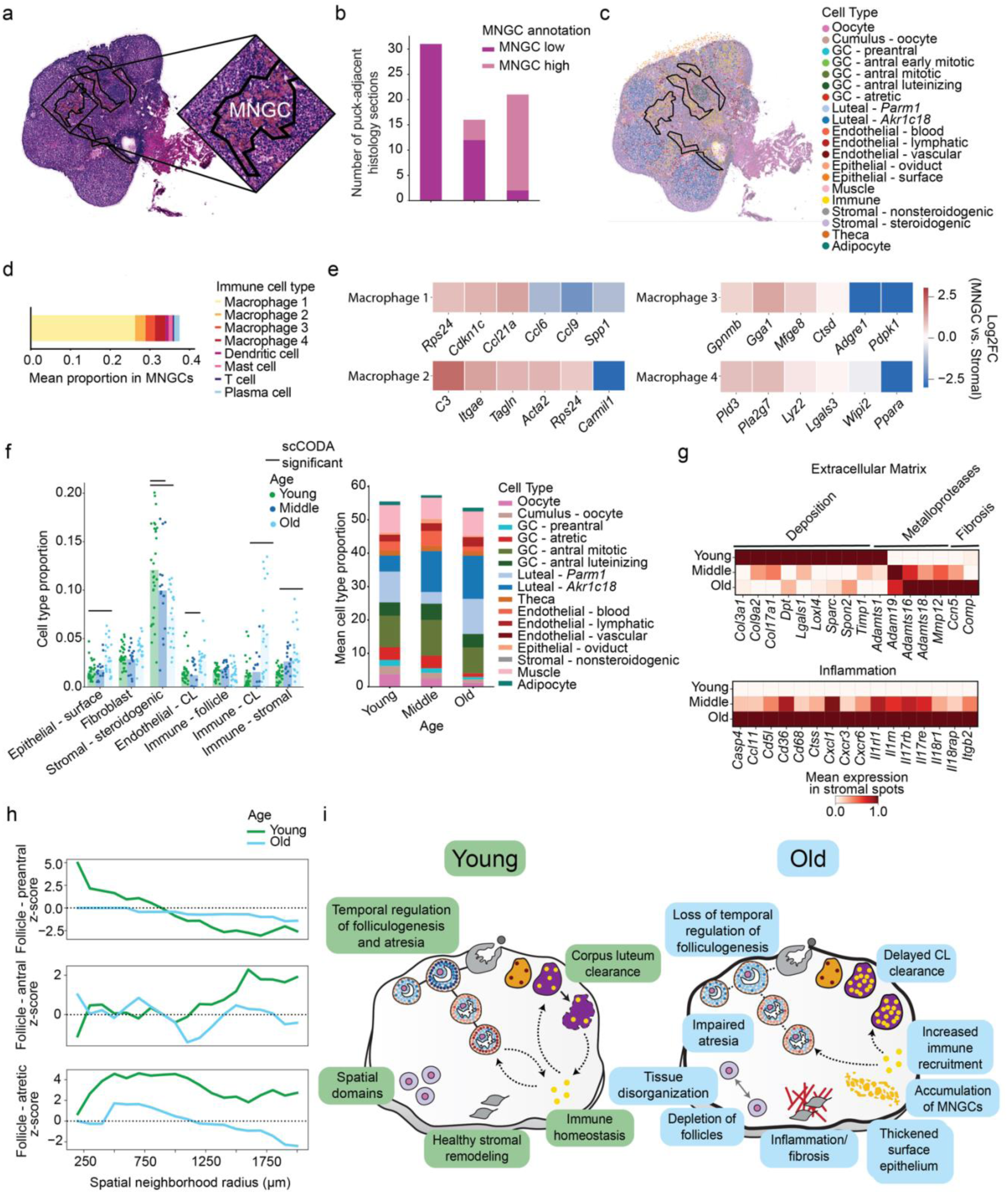
Loss of tissue-wide homeostatic remodeling with aging. **a.** Identification of multi-nucleated giant cells (MNGCs) in aged ovaries. Example H&E section with manually drawn MNGC masks (**Methods**). **b.** Distribution of histology sections (y axis) annotated as MNGC-high or MNGC-low across young (*n*=31), middle (*n*=16), and old (*n*=21) pucks (x axis). **c.** Example registration of MNGC masks from adjacent H&E to Slide-seq using Napari (**Methods**). Slide-seq spots are colored by annotated cell type (colors). **d.** Mean proportions of immune subtypes within MNGC regions in old animals. **e.** Log2 fold changes in representative genes across macrophage subsets (rows), comparing stromal macrophages inside vs. outside MNGCs in old animals. Spot counts in **Extended Data Table 4**. **f.** Age-associated compositional changes in stromal populations. Left: Proportion (y axis) of select cell types (x axis) for individual pucks (dots) and mean across pucks (bars); significant changes identified by scCODA using all cell types with “Epithelial – oviduct” as the reference (**Extended Data** Fig. 6d; **Methods**; black bars, FDR=0.05). Right: Mean proportion of all remaining cell types (y bars) across ages (x axis); puck counts: young *n*= 32, middle *n*=16 pucks, old *n*=21 pucks. **g.** Mean expression (log normalized, variance-scaled) of ECM- and inflammation-related genes (columns) in stromal subtypes (“stromal – non-steroidogenic”, “stromal – steroidogenic”, “fibroblast”) across young (*n*=9), middle (*n*=6) and old (*n=7*) mice (rows). Genes identified by differential expression analysis between young and old stromal spots (**Methods**); spot counts in **Extended Data Table 4**. **h.** Spatial neighborhood analysis of homotypic follicle co-localization. Z-scores of co-localization (y axis) as a function of increasing spatial radius (x axis; 200 µm steps) compared to a null model (z-score 0, black dotted line) for preantral (top), antral (middle), atretic (bottom) follicles in young (*n*=9) and old (*n=7*) mice (colors); follicle counts by age in **Extended Data Table 4**. **i.** Schematic hallmarks of ovarian aging. Aging impacts key processes underlying tissue-wide homeostasis, impacting both the reproductive and endocrine functions of the ovary. Young ovaries maintain temporal regulation of folliculogenesis, spatial domain integrity, immune homeostasis, and coordinated remodeling. Aged ovaries show disrupted timing, impaired atresia and CL clearance, tissue disorganization, follicle loss, MNGC accumulation, inflammation, fibrosis, and surface epithelial thickening.

Ovarian aging is marked by global shifts in tissue composition and function, including increased inflammation that may broadly impair tissue homeostasis^11,116^. We examined alterations in the stroma, including all cell types located predominately outside of follicles and corpora lutea. Analysis of stromal cell populations revealed an age-associated increase in surface epithelial and immune cells, coupled with a decline in steroidogenic stromal cells, progressing monotonically from young to middle-aged and old mice^10,116–119^ (**Fig. 7f**; compositional test with scCODA^120^; **Extended Data Fig. 6d**; **Methods**). The thickening of the surface epithelium was independently confirmed by morphometric measurements on adjacent H&E sections (**Extended Data Fig. 6e**; **Methods**). These compositional shifts coincided with transcriptional changes characterized by reduced expression of extracellular matrix genes, increased metalloproteases involved in tissue remodeling, and elevated markers of fibrosis and inflammation^116^ (**Fig. 7g; Extended Data Fig. 6f**). Together, these changes suggest that aging disrupts immune-mediated maintenance of stromal homeostasis, promoting chronic inflammation and fibrosis within the ovarian microenvironment.

### A model for global tissue disorganization in ovarian aging

We hypothesized that age-related remodeling of the ovarian stroma during disrupt coordination between follicles. To test this, we quantified the spatial co-localization of homotypic follicles, i.e. those of the same follicle developmental stage. At short length scales (<500 µm), both preantral and atretic follicles were highly co-localized in young but not aged mice (**Fig. 7h**; **Methods**), consistent with prior reports that follicle crowding facilitates early activation and preantral development^121,122^. The loss of preantral clustering with age likely reflects, in part, increased extracellular matrix rigidity^9,11,116^. Co-localization of atretic follicles in young ovaries suggests a role for spatially regulated paracrine signaling during follicle selection, potentially influenced by the dominant follicle. In contrast, antral follicles did not co-localize at short length scales in either young or old mice, consistent with their function as isolated “islands” of dominant follicles during selection. Modest co-localization at larger scales (> 2,000 µm) in both young and old mice may simply reflect the overall larger size of the ovary in young mice (∼3,000 µm in total diameter). Together, these data reveal that ovarian aging reduces follicular crowding, disrupting the spatial coordination of follicle development and interfollicular communication.

The loss of global spatial organization in aged ovaries suggests that inflammation disrupts both individual follicle function and their coordinated development, marking molecular hallmarks of ovarian aging across cells, niches and the entire tissue (**Fig. 7i**). At the cellular level, oocytes function declines as the number of maturing oocytes decreases and their transcriptional states shift with age. These changes are compounded by alterations in surrounding niches: inflammatory gene expression increases within follicles, and their responsiveness to the estrous cycle diminishes. In old mice, this is reflected by preantral follicles that no longer vary by cycle stage and by the accumulation of late corpora lutea due to defective clearance. At the organ level, the surface epithelium thickens, and the stroma becomes dominated by chronic inflammation and multinucleated giant cells. Together, these findings show that age-related inflammation progressively erodes ovarian organization and remodeling across multiple spatial scales, from cells and niches to the whole organ.

## Discussion

Here, we present a 10µm-resolution, >600,000 spot spatial transcriptomic atlas of the naturally cycling murine ovary across reproductive ages, revealing that aging disrupts the spatial and temporal coordination of the ovarian tissue environment across multiple scales. Aging attenuates oocyte transcriptional programs, erodes the spatial polarity of granulosa cells, and alters the timing of follicle maturation and CL clearance. These changes coincide with increased immune infiltration, MNGC formation, and loss of global tissue coordination, collectively defining ovarian aging as a progressive tissue-wide breakdown of spatiotemporal regulation rather than a singular decline in oocyte number or quality. The extended presence of CLs expressing progesterone-metabolizing and immune recruitment genes coupled with an expansion of infiltrating immune cells may trigger chronic inflammation, while the emergence of MNGCs containing immune cells with altered signaling marks a shift to sustained inflammatory and lipid-associated activity. In parallel, loss of spatial coupling between neighboring follicles suggests that extracellular matrix remodeling and paracrine signaling required for coordinated development deteriorate with age. Together, these changes precede the cessation of cycling, propagating local follicular and luteal spatiotemporal regulation into global tissue disorganization, representing early molecular and structural hallmarks of reproductive aging.

During each estrous cycle, numerous follicles compete for dominance and ovulation, with the majority undergoing atresia. The spatial organization of GCs within follicles may represent an elegant mechanism to efficiently compete for metabolic resources while sustaining coordinated growth. By segregating proliferative activity to inner GCs and hormone-sensing, steroidogenic functions to outer GCs positioned near the vasculature and stroma, follicles optimize resource allocation under the energetic constraints of this competitive environment. We hypothesize that the success of spatial compartmentalization of GC function during folliculogenesis could contribute to follicular fate, with dominant follicles effectively maintaining functional polarity while less efficient competitors undergo atresia^123,124^. In addition, our results support and refine current understanding of follicular atresia as a heterogeneous process that can arise at multiple follicular stages. We identified engagement of metabolic reprogramming, stress response, extracellular matrix remodeling, and immune-modulatory pathways as well as apoptotic pathways that are concentrated at the follicle edge and vary with follicle state and estrous cycle context. While our results outline a plausible sequence of atresia commitment arising from multiple GC lineages, the specific molecular drivers require experimental validation^17,125–127^.

Our findings highlight the critical role of temporal coordination in ovarian function. While the ovulation cycle is often viewed primarily as a mechanism for oocyte release, our results highlight the tissue-wide remodeling required for successful cycling. In young mice, these processes are tightly coupled to the estrous cycle, reflecting finely tuned synchronization between local follicular processes and systemic endocrine signals consistent with previous reports^98–101,103,104^. With age, however, this temporal pattern deteriorates: follicles lose stage-and follicle-type restricted expression of folliculogenesis and atretic pathways and CLs persist across stages with altered expression in progesterone metabolism. This mirrors luteal-phase changes described in perimenopausal women and suggests that reproductive aging manifests as a breakdown in cyclical synchronization between local tissue dynamics and systemic hormonal rhythms. We note that these multicellular features of reproductive function in young mice and their progressive dysfunction in old mice are exceptionally dynamic with an estrous cycle length of only four days, allowing spatial transcriptomics to capture snapshots of meaningful niche transitions that describe changes across multiple scales, from radial heterogeneity within follicles, to variation across follicles, and spatial patterns of follicle co-localization that underscore the importance of interfollicular coordination.

These results have potential implications for understanding and mitigating human reproductive aging, which remains challenging to profile at scale and high temporal resolution. The observation that spatial and temporal coordination break down before cycle cessation suggests new approaches for detecting and potentially intervening in age-related fertility decline. Additionally, the role of tissue-wide organization in maintaining function suggests that therapeutic strategies may need to target the entire ovarian environment rather than focusing solely on oocytes or follicles.

Several key questions requiring targeted perturbation and ability to track follicular fate emerge from our findings. How do GCs maintain distinct functional zones despite their high proliferation rate? Would improved compartmentalization alter atresia? What signals coordinate states between neighboring follicles? How are atretic GCs, regressing luteal cells and infiltrating immune cells cleared from the tissue? Would interventions targeting cellular turnover or inflammatory pathways preserve ovarian function with age? Understanding these mechanisms could reveal new principles of tissue remodeling relevant not only for ovarian function but also regeneration and aging in other contexts.

We acknowledge several limitations in our study. While spatial transcriptomics provides unprecedented insights into tissue organization, we lack direct observation of fate decisions and cannot pinpoint the exact step at which follicles commit to atresia, which follicles contain successfully ovulated oocytes, or the age of a corpus luteum. Future studies integrating lineage tracing and hormone measurements across the full estrous cycle would provide a more complete picture of ovarian temporal dynamics and their changes with aging. Additionally, validation in human tissue will be important for translating these findings into clinical applications.

Here we demonstrate that ovarian function requires precisely coordinated spatial and temporal organization across multiple scales, from cell-type specific patterning within follicles, stage specific coordination of follicles and CLs, to tissue-wide cellular remodeling. Spatial transcriptomics technologies can capture meaningful multicellular transitions if carefully analyzed, positioning this methodology as a key instrument to derive mechanistic insights into reproductive function and decline, opening new avenues for preserving fertility and ovarian endocrine function.

## METHODS

### Mice

*Buck mice.* Female C57BL/6J (Jax 000664) mice were bred at the University of San Francisco (UCSF). All mice were maintained under SPF conditions on a 12-hour light-dark cycle, at ambient temperature 21.5 + 1°C and relative humidity between 30% and 70% and provided food and water ad libitum. All mouse experiments were approved by and performed in accordance with the Institutional Animal Care and Use Committee guidelines at UCSF on a mouse protocol under Zachary Knight.

*Broad mice*. Female C57BL/6J (Jax 000664) mice were purchased from The Jackson Laboratory at ages 10, 36, and 50 weeks. Mice were acclimated to the vivarium for 2 days (“Broad cohort 1”) or 2 weeks (“Broad cohort 2”) upon arrival. All mice were maintained under SPF conditions on a 12-hour light-dark cycle, at ambient temperature 21.5 ± 1°C and relative humidity between 30% and 70% and provided food and water ad libitum. All mouse experiments were approved by and performed in accordance with the Institutional Animal Care and Use Committee guidelines at the Broad Institute on a mouse protocol under H.C. and F.C.

*Yale mice.* Female C57BL/6J (Jax 000664) mice were purchased from The Jackson Laboratory at ages 8, 36, and 42 weeks. Mice were acclimated to the vivarium for 14 days upon arrival and maintained under SPF conditions on a 12-hour light-dark cycle, at ambient temperature 21.5 ± 1°C and relative humidity between 30% and 70%, with food and water provided ad libitum. All mouse experiments were approved by and performed in accordance with the Institutional Animal Care and Use Committee guidelines at Yale University on a mouse protocol under H.C.

### Estrous cycle staging mice by vaginal cytology

*Buck mice.* Mice collected at UCSF were staged near-daily for 8 consecutive days using vaginal cytology, as previously described^128,129^. A total volume of 100 µL sterile saline (in 20-50 µL increments) was expelled 5-10 times with a P200 pipette into the vaginal canal, taking care to avoid touching the vaginal opening with the pipette tip then expelled onto a glass slide and imaged immediately using Nikon Eclipse Ts2 (10X objective) and a Nikon DS Qi2 camera.

*Broad and Yale mice.* Female mice were staged near-daily for 8 consecutive days using vaginal cytology, as previously described^128,129^. Briefly, mini-tip flocked nasopharyngeal swabs (VWR SLIN501CS01) were moistened with sterile PBS, and ∼1/4^th^ of the flocked tip was inserted into the vaginal canal of manually restrained mice. Swabs were rotated 5 times clockwise, 5 times counterclockwise, and then smeared onto a non-treated glass slide (VWR #16004-422). Slides were air-dried and stained with a 0.1% crystal violet solution (Jorgenson #J0322) for 1–2 minutes. Excess stain was rinsed with PBS and sterile water, and the slides were counterstained with 1% eosin (Jorgenson #J0322A) for 10–20 seconds. Following the counterstaining step, slides were rinsed again, air-dried completely, examined under a light microscope, and imaged with a Leica Aperio VERSA Brightfield, Fluorescence & FISH Digital Pathology Scanner, using the 20x objective.

Cytology evaluation identified the predominant cell types (epithelial cells, cornified cells, and leukocytes), enabling the classification of the estrous cycle stages (proestrus, estrus, metestrus, and diestrus) as previously described^128,129^. In brief, proestrus, estrus, and diestrus are characterized by a predominance of nucleated epithelial cells, cornified epithelial cells, or leukocytes, respectively, while metestrus is identified by a mixture of cornified epithelial cells and leukocytes^130^. The time course of the estrous cycle is in **Extended Data Table 1**.

### Tissue collection

Once identified to be in the appropriate estrous stage for collection, mice were placed under anesthesia with 3% isoflurane for 1 minute and maintained via a nose cone flowing 1% isoflurane. Anesthesia was confirmed by the absence of a pedal reflex. Transcardial perfusion was performed with PBS through the left ventricle to remove blood from tissues before entire ovaries were removed and immediately frozen on dry ice in a cryomold containing Optimal Cutting Temperature (OCT) compound. Tissues were then either processed immediately or stored at -80°C for storage.

### Hematoxylin and eosin stain of adjacent histology sections

Using a Leica ST5010 Autostainer XL (Leica Biosystems), 10 µm sections from fresh frozen tissues were immersed in xylene, sequentially processed through a graded ethanol series (100% and 95%), and then stained with hematoxylin (Leica Biosystems 3801571). The nuclei were differentiated with mild acid treatment (Leica Biosystems 3803598), followed by "bluing" through exposure to a weakly alkaline solution (Leica Biosystems 3802918) and subsequent water washing. Sections were then stained with Eosin (Leica Biosystems 3801616), processed through a graded ethanol series (100% and 95%), xylene, dehydrated, and finally coverslipped using the Leica CV5030 Fully Automated Glass Coverslipper. Brightfield images were captured with the Leica Aperio VERSA Brightfield, Fluorescence & FISH Digital Pathology Scanner, using the 20x objective.

### RNA-FISH via Hybridization Chain Reaction (HCR)

RNA in situ hybridization was performed on 10 µm-thick cryosections using the HCR v3.0 (Molecular Instruments) for fresh frozen tissue sections with the HCR-Gold RNA-FISH kit according to the manufacturer’s instructions. Probes targeting mouse genes were obtained from Molecular Instruments without custom design. Sections were fixed with ice-cold 4% paraformaldehyde (PFA) in PBS for 15 minutes at 4°C, then sequentially immersed in 50%, 70%, 100% EtOH at room temperature, and washed with PBS. Samples were pre-hybridized in 50 µl of pre-warmed HCR hybridization buffer for 10 minutes at 37°C in a humidified chamber. Probe solution (2 µL of each probe in 100 µL HCR Hybridization Buffer) was applied to tissue sections, covered, and incubated overnight at 37°C in a humidified chamber. Sections were washed four times with pre-warmed HCR probe wash buffer for 15 min each at 37°C. For amplification, samples were pre-amplified with 50 µL of HCR Gold Amplifier Buffer for 30 min at RT in a humidified chamber. Hairpins h1 and h2 (2 µL each) were heated at 95°C for 90 sec, cooled to RT in the dark for 30 min, diluted in 100 µL HCR Gold Amplifier Buffer, and 50 µL of this amplification solution was added to the sample. Slides were covered and incubated overnight at RT in the dark. Samples were washed four times with HCR Gold Amplifier Wash Buffer for 15 minutes each at room temperature in the dark, then dried by blotting with a Kimwipe. Nuclei were counterstained with DAPI (1 µg/mL in PBS, Miltenyi #130-111-570) and washed once with PBS. Autofluorescence was quenched by incubating samples with TrueBlack (1x in 70% ethanol, Biotium #23007) for 1 minute at room temperature, followed by three PBS washes. Samples were mounted with antifade mounting media (Thermo Fisher #P10144) and imaged on an Andor Dragonfly 505 spinning disk confocal system. Images were analyzed using ImageJ software.

### Multiplexed immunofluorescence of immune markers

Fresh-frozen tissue samples were embedded in optimal cutting temperature (OCT) compound and cryosectioned into 10 μm serial sections. Sections were mounted onto SuperFrost microscope slides (VWR, Cat# 48311-703) and stored at −80 °C until staining. Slides were fixed in 4% paraformaldehyde (PFA) for 10 minutes, then transferred to the Molecular Cytology Core Facility at Memorial Sloan Kettering Cancer Center (MSKCC). Slides were blocked for 20 minutes in phosphate-buffered saline (PBS) containing 0.2% Triton X-100, 5% normal goat serum (NGS), and 5% bovine serum albumin (BSA). Iterative immunofluorescence was performed on the Lunaphore COMET staining platform over 11 cycles using Alexa Fluor 555- and 647-conjugated secondary antibodies. Primary antibodies were applied as follows: CD11b (Abcam, Cat# ab182981, 0.125 µg/mL, 4 min exposure, cycle 3), Ki-67 (Invitrogen, Cat# 14-5698-82, 1:2000 dilution, 8 min exposure, cycle 5), and CD3 (Roche, Cat# N03789, 1:2 dilution, 8 min exposure, cycle 9). Following COMET imaging, slides were transferred to a Leica automated staining platform for antigen retrieval using pH 9 retrieval buffer for 15 minutes, followed by staining with CD45 (Proteintech, Cat# 60287-1-Ig) using an Alexa Fluor 594-conjugated secondary antibody. Images from both platforms were aligned for analysis.

### Single nucleus RNA sequencing

Nuclei were isolated from three frozen ovarian tissue, one from each age group at estrus, using EZ-lysis buffer supplemented with recombinant RNase inhibitor (1 μL/mL). Pre-chilled douncers were prepared by washing with RNase Zap, 70% ethanol, and ultrapure water. Tissue was transferred to a douncer containing 2 mL of EZ-lysis buffer and homogenized with 20 strokes each of pestle A and pestle B. The homogenate was transferred to a 15 mL conical tube, brought to 5 mL total volume with EZ-lysis buffer, incubated on ice for 5 minutes, and centrifuged at 500*g* for 5 minutes at 4°C. The pellet was washed once more with 5 mL EZ-lysis buffer using the same incubation and centrifugation conditions, then resuspended in 200 μL PBS containing RNase inhibitor and filtered through a 40 μm strainer. Three V4 (GEM-X) 3’ 10x channels were loaded, each channel with 20,000 nuclei from one sample. Reverse transcription, cDNA amplification, and library construction were conducted according to the standard 10X Genomics single cell 3’ V4 protocol.

### *In situ* transcriptome processing via Slide-seq

Tissue sections were processed and sequenced using Slide-seq v2 as previously described^5^. Slide-seq arrays were purchased from Curio Biosciences (Curio Seeker) or prepared in-house using custom-synthesized barcoded beads functionalized with the sequence^30^ (5’TTT_PC_GCCGGTAATACGACTCACTATAGGGCTACACGACGCTCTTCCGATCTJJJJJJJJTCTTCAGC GTTCCCGAGAJJJJJJNNNNNNNVVT30-3’) with a photocleavable linker (PC), a bead barcode sequence (J, 14 bp), a UMI sequence (NNNNNNNVV, 9 bp), and a poly dT tail. The sample table of all Slide-seq data, denoting the puck origin and other metadata, is in **Extended Data Table 2**.

OCT-embedded frozen tissue samples were warmed to -20°C, sectioned to 10 μm on a cryostat (Leica CM3050S), placed on a pre-cooled Slide-seq array and melted onto the array by warming it briefly. For each sample, 2-3 sections for Slide-seq were collected >200 μm to maximize sampling diversity of tissue sub-structures and consecutive sections were collected for hematoxylin and eosin staining. The affixed tissue section and array was moved into a 1.5 mL Eppendorf tube for tissue digestion and transcriptomic library preparation as previously described^5,6^. The last library amplification was performed in a final volume of 200 μL of PCR mix, divided into four 0.2 ml tubes and cycled with the following program: 1 cycle of 98°C for 2 min, 4 cycles of 98°C for 20 s, 65°C for 45 s, 72°C for 3 min, 9 cycles of 98°C for 20 s, 67°C for 20 s, 72°C for 3 min, and 1 cycle of 72°C for 5 min. Libraries were sequenced using the following read structure on a NextSeq 1000/2000 (P2; Illumina) or NovaSeq (S4 or X; Illumina): Read1: 42 bp; Read2: 50bp; Index1: 8 bp, and sequences were processed as previously described^5^ using the pipeline available at (https://github.com/MacoskoLab/slideseq-tools).

### Hematoxylin and eosin stain of adjacent histology sections

Serial sections of 10 μm thickness were obtained from the frozen tissue samples and mounted onto glass microscope slides (VWR SuperForst 48311-703). Using a Leica ST5010 Autostainer XL (Leica Biosystems), sections were immersed in xylene, sequentially processed through a graded ethanol series (100% and 95%), and then stained with hematoxylin (Leica Biosystems 3801571). The nuclei were differentiated with mild acid treatment (Leica Biosystems 3803598), followed by "bluing" through exposure to a weakly alkaline solution (Leica Biosystems 3802918) and subsequent water washing. Sections were then stained with Eosin (Leica Biosystems 3801616), processed through a graded ethanol series (100% and 95%), xylene, dehydrated, and finally coverslipped using the Leica CV5030 Fully Automated Glass Coverslipper. Brightfield images were captured with the Leica Aperio VERSA Brightfield, Fluorescence & FISH Digital Pathology Scanner, using the 20x objective.

### Pre-processing of single nucleus and spatial transcriptomics data

Raw sequencing reads were demultiplexed using Cellranger (v.6.1.2) mkfastq (10x Genomics)^131^ and aligned to the mouse mm10 reference genome using STAR v2.7.8a with default parameters^132^. For spatial data, spatial barcode assignment and bead-based feature extraction were performed using the Slide-seq pipeline (v2022-01-01) and represented as a gene-by-spot expression matrix directly within the Slide-seq pipeline for downstream analyses^5,6^.

### Initial data processing

Gene expression data of individual samples or pucks were concatenated using an outer join. For spatial data, spatial coordinates were assigned by matching barcodes from puck-specific barcode matching files. Metadata, including puck and sample IDs, estrous stage, and animal age category, were appended to the AnnData object. The combined dataset was filtered to retain spots with at least 50 detected genes and 150 UMI counts. For spot clustering and embeddings, all gene expression values were scaled and centered. The top 5,000 highly variable genes were identified from log-normalized counts across all batches using Scanpy’s highly_variable_genes function (batch key=puck)^133^. Dimensionality reduction was conducted by first regressing out total UMI counts on log-normalized counts, then conducting principal component analysis (PCA) on highly variable genes, followed by correcting for puck-specific batch effects using a PyTorch implementation of Harmony^134^, then constructing a nearest-neighbor graph with *k*=10 neighbors using the top 40 principal components. Unsupervised clusters of spots were identified with the Leiden algorithm^135^ (resolution=1.5) and visualized using UMAP^136^, all implemented in Scanpy^133^ (v1.10.1).

#### Smear removal

Smearing artifacts were manually identified and removed using an interactive selection tool. Spatial scatter plots were generated using Squidpy to visualize cell spots color-coded by cluster identity^137^. Using an interactive LassoSelector widget, regions of spots suspected of containing technical artifacts were removed and excluded from further analyses. This process was repeated for each puck, resulting in refined AnnData objects free from smear artifacts. Following smear removal, the dataset was re-processed to refine dimensionality reduction and clustering as described above, resulting in a total of *n*=601,831 spots from 68 spatial transcriptomes.

### Projection of additional pucks

To incorporate additional Slide-seq data into the existing reference atlas, raw gene-by-spot matrices from newly generated pucks (young, metestrus stage) were first pre-processed as described above. Briefly, spatial coordinates were assigned, metadata were appended, and smear artifacts were removed using the same interactive lasso-based filtering procedure.

To ensure compatibility between the query puck and the reference atlas, we harmonized the gene sets prior to integration. Specifically, genes present in the reference but absent in the query, as well as query-only genes, were identified. The query AnnData object was then subset and reordered to the intersection of reference and query genes, yielding a matched feature space for projection. The cleaned query AnnData was projected onto the existing atlas using scvi-tools (v1.1.6) to learn a shared latent representation. The pretrained SCVI model was reloaded and the query puck was registered as a new dataset. Latent embeddings for the query were obtained and concatenated with reference embeddings for downstream analysis.

For label transfer, scArches^138^ (v0.6.1) was used to adapt the pretrained SCVI model via the unsupervised surgery pipeline as described in the scArches documentation. This enabled alignment of the query data with the harmonized latent space and probabilistic annotation transfer. Spot-level labels were assigned for both fine-grained and major cell type categories. To increase annotation robustness, low-confidence calls were masked using an uncertainty threshold of 0.2. Spatial segmentation was performed as described below.

### Unsupervised clustering and cell type annotation

Cell types were annotated using top differentially expressed genes (DEG) identified for each Leiden cluster and cross-referenced with established marker genes from literature^37,40,125,139–145^. DEGs were identified using Scanpy’s rank_genes_groups function with the Wilcoxon rank-sum test to detect cluster-specific marker genes. Spatial scatter plots and UMAP embeddings were used to visualize marker gene expression and confirm cluster-specific identities. Granulosa cells (GCs) were further subclustered by re-running Leiden clustering on GC subsets, followed by DEG analysis and marker-based assignment. The top 50 marker genes identified for each cluster are in **Extended Data Table 3**.

### Immune and granulosa cell subclustering

To refine annotation of specific populations, we performed iterative subclustering of immune cells and granulosa cells (GCs). First, AnnData objects were subset to include only the cell types of interest (e.g., Immune, atretic GCs). Raw counts were normalized to a total of 10,000 transcripts per cell with highly expressed genes down-weighted (max_fraction=0.05), followed by log-transformation. Highly variable genes were identified using Scanpy’s highly_variable_genes function (flavor=“seurat”, batch_key=“segment_label”), and the data were scaled to unit variance with a maximum value of 10. Principal component analysis (PCA) was then performed on the highly variable gene set, and neighborhood graphs were constructed using the top 50 PCs as input.

UMAP embeddings were generated for visualization, and Leiden clustering was used to define subclusters within each subset. Differentially expressed genes (DEGs) were identified for each subcluster using the Wilcoxon rank-sum test (min log fold-change ≥ 1) on normalized counts. For annotation, we required that selected DEGs be (i) specific to the cluster, (ii) highly expressed across the majority of cells within that cluster, and (iii) supported by prior literature as having cell type–defining functional roles. Representative markers were visualized using dot plots to confirm specificity and coverage.

For immune cells, the first round of subclustering resolved broad classes including macrophages, T cells, plasma cells, dendritic cells, and mast cells. Because macrophages formed a large and heterogeneous population, we performed a second round of subclustering restricted to this compartment, which enabled identification of functionally distinct macrophage subtypes supported by known marker genes. A similar iterative approach was applied to GCs (**Extended Data Fig. 1b)**. Broad subclustering resolved major GC states, after which the atretic GC compartment was isolated and re-clustered separately. This refinement revealed transcriptionally distinct atretic GC subpopulations, consistent with prior descriptions of heterogeneity in follicular atresia. For a detailed description of the iterative subclustering strategy, DEG-based marker selection, and literature-based justification for GC and immune subtype annotation, see **Supplementary File 1**. The top statistically ranked DEGs for each GC subcluster are provided in **Extended Data Table 3**.

### Comparison of luteal populations with published datasets

To assess the concordance of our luteal populations with previously published corpora lutea (CL) classifications, we compared our data to the well-annotated single-cell RNA-seq dataset from Morris et al. (2022)^37^. Raw clustered AnnData objects were obtained and reprocessed. For clarity, we excluded the “Regressing CL” cluster reported in Morris et al. as this population originates from post-partum ovaries, which were not part of our study design. Cell type composition was quantified using contingency tables of cluster identity against estrous stage. To directly compare gene expression profiles, we visualized differentially expressed genes (DEGs) in luteal subtypes identified in both our dataset and Morris et al. (2022) as dotplots. Gene-level concordance was assessed by overlapping marker sets and expression distributions across annotated luteal subtypes.

### Spatial segmentation of oocytes, follicles, and CL

Code underlying these analyses can be found in II.2_segmentation.ipynb and II.3_segment_extension.ipynb.

#### Oocyte segmentation

We defined spots in the ‘Oocyte’ or the ‘Oocyte - cumulus’ spot cluster as putative oocyte spots. ‘Oocyte - cumulus’ spots were included in initial segmentation to ensure robustness against variations in cluster boundaries between ‘Oocyte’ and ‘Oocyte - cumulus’ spots but were subsequently removed from the final oocyte segments as described below. For each puck, a spatial neighborhood graph of putative oocyte spots (radius=50) was constructed and clustered with the Leiden algorithm^135^ (resolution=0.1) to define preliminary oocyte segments. ‘Oocyte - cumulus’ spots were then removed from these segments. Segments containing fewer than three spots were also discarded. Final oocyte segments were further refined through subclustering, merging, and manual curation based on tissue context. Details of these curation choices are documented in the supplementary code.

#### Additional decontamination of cumulus spots from oocyte spots

Non-negative matrix factorization (NMF)^146^ was used to identify gene programs corresponding to cumulus cell contamination in spots labeled as oocyte from Leiden clustering. scikit-learn^147^ was used to perform NMF with k=8 programs on log1p normalized counts. Programs 1 (*Kctd14*, *Slc18a2*, *Wt1*), 2 (*Lars2*, *Hexb*, *Cdk8*), 4 (*Inhba*, *Inhbb*, *Nap1l5*), and 6 (*Col1a2*, *Col3a1*, *Cfh*) were identified as contaminating signals based on their constituent genes (**Extended Data Fig. 7a**). Spots were annotated as contaminants if they either had these programs as their top utilized program or if the program scores reached the following thresholds: Program 1: > 0.4, Program 2: > 0.3, Program 4: > 0.3, Program 6: > 0.4 (**Extended Data Fig. 7b**). Spots annotated as contaminants were removed from the object before a second round of decontamination was performed on remaining spots using NMF with k=8 programs. Program 1 (*Serpine2, Ccnd2, Kctd14*) was identified as a contaminating signal based on its constituent genes (**Extended Data Fig. 7c**). Spots that had this program as the top utilized program, or had a program score >0.3 were annotated as contaminants (**Extended Data Fig. 7d**) and removed from the oocyte segments for the computation of these averages.

#### Identifying primordial oocytes

To identify any remaining small, putatively primordial, oocytes, unassigned spots labeled as ‘Oocyte’ or expressing primordial oocyte marker gene *Sohlh1*^36,55^ that were at least 200 from an oocyte segment center were selected. A spatial neighborhood graph of these putative primordial oocyte spots for pucks containing at least two such spots using a radius of 20 µm to account for the expected small size of these primordial oocytes. Oocyte segments were identified after Leiden^124^ clustering (resolution=0.1). Spots labelled as ‘Oocyte - cumulus’ were removed from these segments. Segments containing less than 2 spots or lacking *Sohlh1* expression were excluded.

#### Segmentation of follicles and corpora lutea

Spots in the ‘Cumulus – oocyte’, ‘GC – preantral’, ‘GC - antral early mitotic’, ‘GC - antral luteinizing’, ‘GC - antral mitotic’, ‘GC – atretic’ clusters were defined as putative follicle spots, while spots in the ‘Luteal - early’, ‘Luteal - late’ clusters were defined as putative CL spots. A spatial neighborhood graph of the putative follicle and CL spots (radius 50 µm) was constructed and clustered with Leiden^135^ (resolution=0.1 for follicles and a smaller resolution=0.001 for CLs because they tended to be larger) for each puck, yielding segments. Segments with less than 20 spots were removed, the remaining were then subclustered and merged as appropriate based on manual inspection. These curation choices are documented in the supplementary code. Segments were then extended to include additional spots from the selected GC or luteal cell types, respectively, within 50 µm of a spot already assigned to a segment or the next closest segment if multiple were within 50 µm.

In a few cases, oocyte segments had been identified by the above oocyte segmentation but the corresponding follicle was either too small or irregular, resulting in partial or complete omission by this follicle segmentation pipeline. To address these cases, unassigned spots from the above selection of follicle cell types within 50 µm of an oocyte segment were assigned as follicle segments and grouped based on the closest oocyte segment. If these new follicle segments were closer than 100 µm (minimal spot-spot distance across a segment pair) to existing follicle segments, the segments were merged.

To capture remaining small follicles, unassigned spots in the ‘Cumulus – oocyte’, ‘GC – preantral’, ‘GC - antral early mitotic’, and ‘GC – atretic’ spot clusters were defined as mapping to putative small follicles. Within each puck, one spatial neighborhood graph was each constructed for these putative follicle and CL spots (radius 20 µm) and clustered with Leiden (resolution=0.1), resulting in additional segments. Segments with less than 5 spots assigned were removed. Remaining segments were merged with existing follicle segments if they were within 50 µm of existing segments (minimal spot-spot distance across segment pair). The final segments were further subclustered, merged, and reassigned to proximal oocytes as appropriate. These curation choices are documented in the supplementary code.

#### Assigning proximal spots to segments

Immune (‘Immune’), endothelial (all ‘Endothelial’ types), theca (‘Theca’ and ‘Stromal - steroidogenic’), and mesenchyme (‘Adipocyte’, ‘Stromal - nonsteroidogenic’) spots were assigned to follicle or CL segments that were less than 100 µm away.

#### Assigning oocytes to follicles

Oocyte segments were assigned to the closest follicle segment if it was less than 100 µm away. These assignments were manually curated where appropriate based on manual inspection of the segmentation results and the surrounding tissue; these curation choices are documented in the supplementary code.

### Follicles and corpora lutea segments labeling

First, segments were assigned as follicle or CL: CL segments were classified as those in which the most frequent spot type was a luteal type, or where the sum of the fraction of luteal types out of above granulosa (GC) and luteal types was larger than 50%; all remaining segments were classified as follicles. (**Extended Data Fig. 1g, h**). Note that this classification was performed on all segments from both the follicle and CL segmentation above to recover exceptional cases in this initial segmentation. Second, more specific state labels were assigned to the resulting follicle and CL segments. Follicle segments were labelled as ‘Follicle – atretic’ if more than 20% of their assigned spots were classified as ‘GC – atretic’. All other follicle segments were classified based on their most abundant GC type (‘GC – preantral’ as ‘Follicle – preantral’, ‘GC – antral luteinizing’ as ‘Follicle – antral luteinizing’, ‘GC – antral early mitotic’ and ‘GC – antral mitotic’ as ‘Follicle – antral mitotic’). CL segments were classified as ‘CL - early’ and ‘CL – late’ based on their majority luteal cell type.

### Validation of the segmentation pipeline with manually segmented adjacent H&Es

To validate the segmentation pipeline, adjacent H&E-stained sections were annotated for preantral follicles, antral follicles, and corpora lutea. Annotated H&E sections were then compared to spatial plots of segmented objects, with ‘follicle - antral luteinizing’,’follicle - antral mitotic’, and ‘follicle - atretic’ combined into a single ‘follicle - antral’ category, as by histology of fresh frozen sections these categories can not be distinguished. For all young sections, follicles were annotated as being consistent between the spatial plot and H&E (89.6%, **Fig. 1e,f, Supplementary File 2,3**) or annotated if there was a mismatch between the two sections. Differences in annotations between the adjacent H&E section and Slide-seq section are mostly due to structures that are not present in both sections–for example, the edge of a follicle was captured in the sequenced section, and the adjacent section that was H&E stained does not contain any cells from that follicle (**Supplementary File 2,3**). The remaining discrepancies are the result of separate follicles being segmented as a single object (7.9%), an incorrect segment identity being assigned (0.6%), and the segmentation pipeline not detecting an object (1.9%) (**Fig. 1f**). Cohen’s kappa was calculated with scikit-learn^147^. Follicle counts and error types were plotted using GraphPad Prism version 10.0.0.

### Calculating mean expression profiles of oocyte, follicle and CL segments

Raw (unfiltered and un-normalized) counts of spots assigned to a segment (either oocyte, follicle, or CL were normalized per spot to 10,000, then averaged within each segment. Where appropriate, the resulting segment expression profiles were log1p transformed for downstream analyses.

### Trajectory inference of GCs

Trajectory inference of GCs was performed using diffusion pseudotime (DPT), a graph-based method that models cellular transitions along a continuum, as described in previous single-cell RNA-seq studies^148^. DPT analysis was conducted on annotated GCs after dimensionality reduction using principal component analysis (PCA). A k-nearest neighbor (k-NN) graph was constructed using the top 40 principal components, with k set to 10, to define local cellular neighborhoods. Diffusion components (DCs) were computed from the neighborhood graph to capture the structure of cellular transitions in high-dimensional space. The root of the trajectory was manually selected based on inspection of DC embedding. DPT values were calculated in two complementary ways, depending on the analysis. For Fig 3d., pseudotime was computed separately for GC spots in non-atretic and atretic follicles to analyze spatial patterns of trajectories in follicles under developmental versus degenerative conditions. For Fig 3h., DPT was computed across all GC spots combined to provide a single ordering of granulosa cell states across the follicular lifecycle and facilitate comparison across follicle types, estrous cycle stage, and age (**Extended Data Fig. 3g**).

### Calculation of cell cycle scores

To assess cell cycle dynamics in annotated granulosa cell spots in mice, canonical markers of S-phase and G2/M-phase, as previously described^149^, were used to calculate cell cycle phase scores. Based on these scores, spots were classified into three phases: G1, S, or G2/M. Spots classified as G1 were further categorized as "Non-Cycling," while spots in S or G2/M phases were categorized as "Cycling." This classification was integrated with spatial transcriptomic data to map the cycling status of annotated granulosa spots. Dimensionality reduction methods were applied to visualize cell cycle phases in relation to spatial organization in tissues, enabling an analysis of proliferative activity within granulosa cell-associated regions of the ovary.

### Radial distance analysis within segments

To examine spatial trends in our analysis, radial distances for spots within follicular segments were computed. The radial distance for each spot was calculated as the Euclidean distance from the centroid of the segment, scaled by the maximum segment radius. This normalized distance, referred to as the "scaled radial distance," enabled a standardized comparison of spatial patterns across segments of varying sizes. Spots were binned along the scaled radial distance into equal intervals spanning the range [0, 1] (**Extended Data Fig. 3a**).

For each radial bin, the mean and standard error of the mean (SEM) of various features such as cell type composition, DPT and cell cycle scores, cNMF-derived gene program usage were first calculated across individual bins. Each mean and SEM were then aggregated for each distance bin within the same segment type. Segment-specific trends were then visualized by plotting the mean proportions or cNMF program scores for each distance bin, with shaded regions representing the SEMs.

To evaluate spatial trends in various features across scaled radial distance bins, the monotonic relationship between the radial bin index (independent variable) and the mean values of the feature of interest (e.g., cell type composition, cell cycle state, DPT, or cNMF program usage) for each segment type was assessed using Spearman rank correlation. For each segment type, the mean feature values within each radial distance bin were calculated as described above and correlated with the corresponding radial distance bins to determine whether the feature of interest varied systematically with distance from the segment centroid. Spearman correlation was chosen for its robustness in detecting monotonic relationships, regardless of the underlying distribution or linearity. The Spearman correlation coefficient (ρ) and associated p-value were calculated using the SciPy library. Bins with fewer than two data points were excluded from the analysis.

### Identifying gene programs with non-negative factorization

#### Oocytes

Gene programs in composite oocytes were identified using scikit-learn^147^ to perform non-negative matrix factorization (NMF)^146^. Log-normalized counts were used for NMF (*k*=5 programs) to reduce the noise of highly expressed genes in composite oocytes that were composed of few oocyte spots. To interpret the biological relevance of the gene programs, genes were ranked for each program by their contributions to the gene weight matrix. Program usage across oocytes were visualized using PCA embeddings.

#### Granulosa cells (GCs)

Gene programs in GCs were identified using consensus non-negative matrix factorization (cNMF), following the workflow described by Kotliar et al^76^. Using raw gene expression counts as input, 5,000 highly variable genes (HVGs) were identified by calculating gene variance across spots during preprocessing with the top 5,000 HVGs retained for downstream analyses. Batch correction was incorporated directly into the cNMF pipeline using an adapted version of Harmony^134^ and the resulting corrected matrix, with negative values set to zero, resulting in the batch-corrected matrix used for factorization. cNMF was run on the corrected matrix to factorize the expression data into two components: a gene weight matrix, in which each gene is associated with specific programs, and a usage matrix, in which each spot is assigned a score reflecting its degree of program activity. The number of programs (k) was set to 16 to balance program stability and error, calculated as the Pearson correlation between factorization results across 20 replicates for each *k*. For the selected *k*, cNMF was re-run with 200 replicates to generate final consensus matrices and extract robust and biologically meaningful programs.

To interpret the biological relevance of the gene programs, genes were ranked for each program by their contributions to the gene weight matrix (**Extended Data Fig. 3f**). Program activity scores were normalized per spot by dividing the raw usage matrix values by the total program activity per spot. Spots were grouped by their dominant GC program, defined as the program with the highest normalized usage score, to assess program separation and overlap (**Extended Data Fig. 3d**). Gene programs were annotated by comparing the top contributing genes against known GC markers and enriched functional pathways from the literature^14,17,25,150,151^.

### Analysis of gene expression programs (GEP) with pseudotime

To investigate transcriptional dynamics during follicle development and atresia, we analyzed cNMF-derived GEP usage scores ordered along diffusion pseudotime (DPT). As described above, DPT was calculated as a unified trajectory across all GC spots. Stage stream plots depict the distribution of follicle type occupancy along pseudotime. For each type, Gaussian kernel density estimates (KDEs) of pseudotime values were computed across a grid of 400 points, normalized at each position, and displayed as stacked area plots. Estrous phase lanes (second row, when available) were generated analogously for estrus and metestrus and plotted as a two-band stackplot to visualize stage occupancy along pseudotime. GEP heatmaps were generated separately for GC spots within non-atretic and atretic follicles. Pseudotime was divided into 500 equal-width bins, and average GEP usage scores were computed per bin.

For each GEP, and within each follicle type and estrous stage, we tested for differences between young and old ovaries using ordinary least squares (OLS) regression with cluster-robust standard errors. Specifically, for each stratum we fit the model Usage_k ∼ Age (with young as the reference) and extracted the p-value for the coefficient corresponding to old. Robust variance estimates were clustered by puck to account for within-puck correlation of spots. Strata lacking both age groups were returned as not estimable. Across all tested GEP × follicle type × stage combinations, p-values were adjusted for multiple testing using the Benjamini–Hochberg procedure, and significance was defined at FDR < 0.05. Significance panels adjacent to the heatmaps summarize the results: white = not significant, light gray = significant in Estrus only, dark gray = significant in Metestrus only, black = significant in both stages. Stars indicate the adjusted significance level (q < 0.05, q < 0.01, q < 0.001). Analyses were performed using pandas for data handling and statsmodels for regression and multiple testing correction (statsmodels.formula.api.ols, statsmodels.stats.multitest.multipletests).

For alternative visualization of this analysis (**Extended Data Fig. 3g**), we also summarized program activity for each cNMF program (Usage_1–Usage_16) using a three-level averaging scheme to respect biological replication. First, GEP usage scores were averaged within each follicle segment (segment_id) to obtain follicle-level profiles. Second, segment means were averaged within each puck (puck) for each follicle type, estrous stage, and age. Third, puck-level means were averaged across pucks to yield the final condition means for Age × Stage × Follicle type. We then constructed an interleaved categorical axis ordered as Age → Follicle type → Stage to enable side-by-side comparisons across ages within each follicle type and stage.

### Compositional analysis of cell and segment types

#### Global cell-type compositional analysis

The spot composition for each cell type was calculated for each puck. Mean composition was then calculated for young, middle, and old ages for each cell type. To determine significance of cell composition changes with age, the scCODA package was used^120^, with young used as the control group, and epithelial – oviduct as the reference cell type, as its composition has low variation across conditions. Subsets of immune spots were created based on their segment assignments to follicles or CL, or stroma if unassigned. Endothelial spots localized to CL were subset from other endothelial spots.

#### Cell-type composition within segments analysis

For each relevant predicted cell type, the fractional composition was computed as the proportion of spots assigned to that cell type within each segment. The mean and standard error of these fractions were calculated across segments and grouped by distinct segment types. These results were used to analyze patterns in cell type composition and visualize the relative contributions of different cell types and segments.

To statistically compare across follicle type and age, we applied a compositional regression framework in which cell type proportions were transformed using a centered log-ratio (CLR) to account for the closed-sum nature of compositional data. For each cell type within each follicle type, we fit ordinary least squares models of the form CLR ∼ Age, using cluster-robust standard errors (clusters = puck) to account for within-sample correlation. Age effects (Young as reference) were tested for each cell type and follicle type, and p-values were adjusted using the Benjamini–Hochberg procedure. Significance was defined at FDR < 0.10. For visualization, mean ± s.e.m. immune subtype proportions were plotted for Young, Middle, and Old ovaries, with colors denoting age groups and asterisks indicating significance levels (q < 0.05, *q < 0.01, *q < 0.001, •q < 0.10).

#### Segment type composition analysis

The relative abundances of different segment types, including follicular and corpus luteum segments, was analyzed across estrous stage and animal age. The proportions of each segment type (e.g., preantral, antral mitotic, antral luteinizing, and atretic segments) were calculated as the fraction of individual segments of the same segment type within each sample (puck). Mean fractions and standard errors were computed across pucks for each experimental condition, grouped by estrous stage and animal age. Statistical differences between the fraction of a particular segment type across different groups of pucks were assessed with t-tests, and significant differences were highlighted in visualizations.

### Follicle embedding and trajectory inference using optimal transport

Code underlying these analyses can be found in III_segment_OT_1_pca_coretypes.ipynb and III_segment_OT_2_plot_results.ipynb.

#### Segment-to-segment distance matrix computation

A distance matrix was computed between any pair of segments of all follicle and CL segments in young mice. For each segment, oocyte, granulosa or luteal spots, and theca spots were considered. A transport matrix from spots in any pair of segments was calculated with optimal transport, using the Sinkhorn algorithm implemented in moscot^92^ (epsilon=0, tau_a=1., tau_b=1., rank=20, initializer=k-means, min_iterations=5, max_iterations=200). This transport matrix was multiplied elementwise with the cost matrix of the optimal transport problem and the sum over all elements in this resulting matrix was used as a distance measure between the two segments.

#### Segment embedding and follicle pseudotime

This segment-to-segment distance matrix was treated as a segment feature space. The k-nearest neighbor graph of segments (k=15) was built from the top 50 PCAs computed and embedded as a UMAP. This embedding was generated both for the union of all follicle and CL segments in young mice, and for follicles in young mice alone. Diffusion pseudotime^148^ was then computed for follicles in young mice based on a manually defined root segment.

#### Trajectory projection of follicles from middle-aged and old mice

A distance matrix was calculated between all follicle and CL segments in young mice and middle and old mice, using the same optimal transport approach as above. Each segment from middle and old mice was then assigned to its closest neighbor in young mice based on this distance. Follicles from middle and old mice were then assigned the pseudotime coordinates from their closest neighbor follicle in young mice.

### Cell–cell communication analysis

Code underlying these analyses can be found in III.5_nichenet_ageYoung_stageAll_segmentFollicle_withOocyte_byCluster.Rmd and III_segment_OT_2_plot_results_v0.3.ipynb.

Cell-cell communication analysis was performed with MultiNicheNet^93^ on data from young mice, with follicle segments considered as samples for the purpose of the algorithm. Comparisons were performed between different follicle labels, using coarse cell type labels (preantral, antral, atretic). Oocyte, granulosa, theca, and immune spots were included in this analysis. Ligand-receptor pairs and receiver-sender cell type pairs specific to each follicle label were prioritized accordingly. Follicles that did not meet default cell type compositional thresholds were discarded (at least 5% of spots per segment mapping to each cell type). To characterize the communication states across all follicles, a summarized ligand-receptor score was calculated for each segment. This score was derived first by calculating the mean of normalized expression of receptor and ligand pairs for all cell types within a segment, log1p transforming, and then summed to generate the final segment specific ligand and receptor score.

### Immune infiltration analysis

To compare the infiltration of immune cells across segment types, immune spot composition was analyzed relative to distance from the segment center. For each scaled radial distance bin (calculated as described above) within individual segments, the proportion of immune spots was calculated by dividing the number of immune spots by the total number of spots within each bin. The average immune composition and standard error of the mean (SEM) for each distance bin were determined for every segment type to assess systematic variation with distance from the segment center. Tukey’s Honest Significant Difference (HSD) test was applied to evaluate the monotonic relationship between the radial bin index (independent variable) and the mean values of the feature of interest (dependent variable) for each segment type. To address potential bias from tissue sectioning, additional analysis was conducted on oocyte-containing follicles to confirm that slicing artifacts did not affect the evaluation of radial infiltration (**Extended Data Fig. 1j**). Statistical comparisons and p-values were calculated using the SciPy library. Radial bins with fewer than two data points were excluded from both statistical and visualization analyses.

### Age-associated differential gene expression analysis

#### Follicles

To analyze key hormone gene changes across young and old follicles, mean gene expression was calculated as described above. For each segment type (preantral, antral mitotic, antral luteinizing, and atretic follicles), stage (Estrus, Metestrus) and age, mean and SEM of gene expression were calculated for the hormone-related genes *Amhr2*, *Fshr*, *and Lhcgr*^47^ . Differential expression analysis between estrous stages was performed using two-sided t-test.

#### Oocytes

Primordial oocytes were removed from analysis due to the low representation in old animals. Genes were filtered to remove those expressed in less than 10% of oocytes. Differential expression analysis was performed using Scanpy on young and old oocytes, using log1p normalized data using t-test with overestimated variance and using the Benjamini-Hochberg correction method. Genes with a p-value <0.05 and a log2 fold change >1 were used for further analysis.

#### Stromal cells

Stromal cells including spots labeled as fibroblasts, stromal – nonsteroidogenic, and stromal – steroidogenic were used for differential expression analysis. Genes were filtered to remove those expressed in less than 10% of spots. Differential expression analysis was performed using Scanpy on young and old spots, using raw counts using t-test with overestimated variance and using the Benjamini-Hochberg correction method. Genes with a p-value <0.05 and a log2 fold change >1 were used for further analysis.

### Identifying multi-nucleated giant cells (MNGCs)

Code underlying these analyses can be found in II.5_registration_v0.8_all_pucks.ipynb and III_MNGC_dotplot_v0.1.ipynb.

#### Annotation of MNGCs in histology sections

Manual annotation of histology sections adjacent to sections used for Slide-seq analysis were used to classify the presence of multi-nucleated giant cells (MNGCs) based on pigmentation, size, and proximity to a growing follicle as previously described^11,33,114,115,152^. H&E stains with a high density of light brown clusters, generally surrounding growing follicles, were categorized as “visible MNGCs” (19 samples). Sections with lighter pigmentation and a lower density of clusters were categorized as “possible MNGCs” (13 samples). Samples lacking visible pigmentation were labeled as “no MNGCs.”

#### Segmentation of MNGCs in H&E and registration with spatial transcriptomics data

Samples deemed to contain MNGCs upon visual inspection of the H&E images were selected for segmentation. H&E images and spot transcriptomics were registered using a napari plugin in squidpy^137^, with anchor points manually defined at tissue features clearly visible in both (such as large oocytes or points on the edges of the tissue section). In this registered space, spots within MNGC regions were segmented based on histology and spot transcriptomics. For the analysis of immune cells in stroma and MNGC, all immune spots outside of follicle or corpus luteum segments were selected and classified as “stromal” (outside of MNGC segments) and “MNGC” (inside of MNGC segments).

### Co-localization analysis of segmented tissue structures

Code underlying these analyses can be found in III_segment_colocalization.ipynb. Within each puck, a spatial neighborhood graph of follicles was constructed to represent each follicle by the spatial center point of its constituent spots. Neighborhood enrichment was performed on this graph with Squidpy^137^ within homotypic follicles (preantral, antral, and atretic) with varying radius from 200 µm to 2000 µm in steps of 200 µm.

### Measurement of surface epithelium thickness

To validate the findings of increased abundance of surface epithelial cells in old animals, surface epithelium thickness was measured using ImageJ^153^. For each section, three thickness measurements were taken and averaged. A single section was measured for each ovary and plotted using GraphPad Prism version 10.0.0.

## Lead contact

Further information and requests for resources and reagents should be directed to and will be fulfilled by the lead contact, Hattie Chung

## Data availability

All sequencing data (Slide-seq) and expression count matrices (Slide-seq) will be deposited in Gene Expression Omnibus and the cellxgene portal prior to publication.

## Code availability

All code is available at https://github.com/HattieChungLab/ovarian_aging

## Author contributions

H.C. and J.G. designed the study. H.C., A.K., I.B., and S.M. conducted the pilot experiment and analysis. A.K., H.C., and T.C.T.L. conducted the mouse work. G.M. created in-house Slide-seq tiles with supervision from F.C. R.R., T.C.T.L., and H.C. conducted Slide-seq experiments. R.R. and T.C.T.L conducted histology experiments and imaging. R.R. processed all Slide-seq data. T.C.T.L. and A.K. conducted cell type clustering and annotation under supervision by H.C., D.S.F., A.K.S., and P.C.S. D.S.F. designed and implemented the segmentation pipeline, optimal transport, and cell-cell interaction analyses. D.S.F., R.R., and R.S. conducted computational histology analyses. H.C. and D.S.F. designed and supervised all other analyses conducted by T.C.T.L., A.K., S.S., A.M., and H.T. H.C. and J.G. supervised the overall project. T.C.T.L., D.S.F., A.K., R.R., S.S., and H.C. interpreted the results with input from all authors. T.C.T.L., A.K., D.S.F., J.G., and H.C. wrote the paper with input from all authors.

T.C.T.L. and A.K. contributed equally.

## Acknowledgments

This work was funded by a Pilot Grant from the Productive Health Global Consortium (formerly GCRLE) awarded to H.C. H.C. also acknowledges support from start-up funds from Yale University and the NIH (R35 GM160070).

T.C.T.L. is in part supported by the Productive Health Pilot Grant awarded to H.C. D.S.F. is supported by the Eric and Wendy Schmidt Center fellowship at the Broad Institute. J.L.G. and A.K. are supported by the NIH (R35 GM145305). We thank Anubhav Sinha for helpful discussions on HCR, and Sami Farhi and the Spatial Technology Platform for allowing us to use their Dragonfly. We thank Murray Tipping, Francesca Mazzoni, and Wenfei Kang at MSKCC. We thank the Broad Institute vivarium staff, especially Nathan Chan, for assistance with vaginal cytology. We thank Nicholas S. Moore for helpful feedback on the manuscript, and all members of the Chung and Garrison Labs and Francesca E. Duncan for helpful discussions.

## Competing interest declaration

A.K.S. reports compensation for consulting and/or scientific advisory board (SAB) membership from Honeycomb Biotechnologies, Cellarity, Danaher, Ochre Bio, Bio-Rad Laboratories, Relation Therapeutics, IntrECate biotherapeutics, Parabalis Medicines, Passkey Therapeutics and Dahlia Biosciences unrelated to this work. P.C.S. is a co-founder of Sherlock Biosciences and Delve Biosciences, a board member of Polaris Genomics, a former board member of Danaher Corporation, and a former scientific advisory board member of NextJenGane, and holds equity in the companies unrelated to this work. F.C. is a founder of Curio Bioscience and Doppler Bio. All other authors report no potential conflicts.

**Supplemental Information** Extended Data Figures 1-7 Extended Data Tables 1-6 Supplementary Files 1-3

---

**Extended Data Figure 1.**
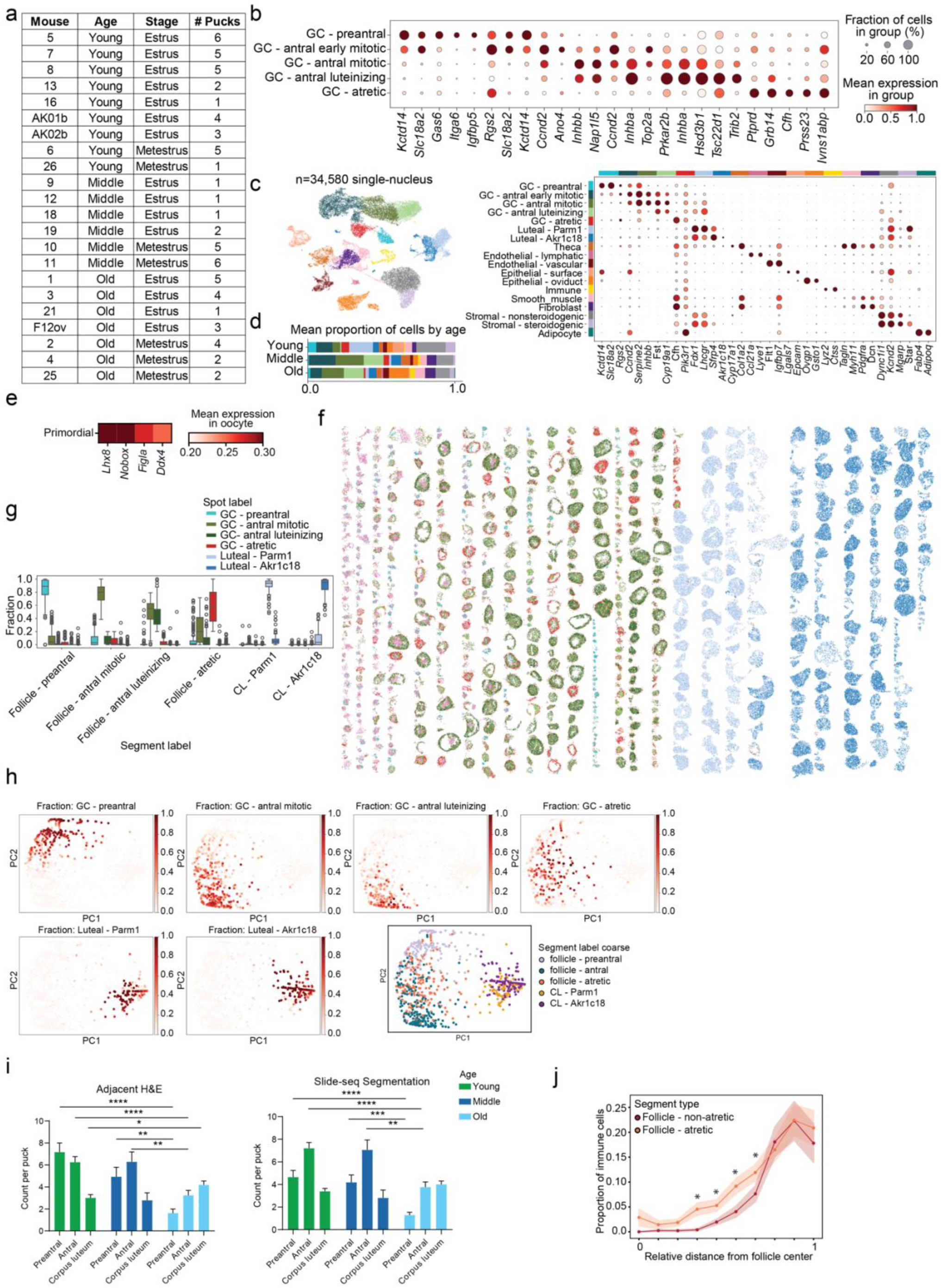
Classification and characteristics of segments driving folliculogenesis. **a.** Table of pucks sequenced per mouse, with age and estrous stage. **b.** Mean expression (color, scaled per gene) and fraction of expressing cells (dot size) of marker genes (columns) defining GC subclusters (rows). Top five differentially expressed genes per subcluster shown (Wilcoxon rank-sum test; min log fold-change ≥ 1). **c.** UMAP embedding of 34,580 nuclei from whole-ovary snRNA-seq colored by major cell type. Dot plot shows mean expression (color, log normalized counts) and fraction of expressing cells (dot size) of marker genes used to annotate granulosa, luteal, stromal, immune, endothelial, and epithelial populations. **d.** Mean cell type proportions across young, middle, and old mice, showing age-associated compositional shifts. **e.** Mean composite expression of primordial oocyte marker genes across primordial oocytes (*n*=57). **f.** Summary spatial plot of all detected follicle and CL segments. **g.** Boxplot of spot-level cell type composition (y axis, colors) across segment labels (x axis). Boxplot: center line indicates median, box bounds represent first and third quartiles, whiskers span from each quartile to the minimum or the maximum (1.5 interquartile range below 25% or above 75% quartiles); segment counts by type in **Extended Data Table 4**. **h.** PCA of segments based on cell type composition. Left: segments by spot counts. Right: segments colored by assigned type. **i.** Mean follicle counts from adjacent H&E and Slide-seq segmentation across age groups. Error bars: SEM. (\**P*<0.05, \*\**P*<0.01, \*\*\**P*<0.001, \*\*\*\**P*<0.0001 two-tailed t-test; all other bar groups, ns); young *n*=31, middle *n*=16, old *n*=21 pucks. **j.** Mean proportion of immune cells (y axis) across radial distance (x axis; 0-1) for each follicle type (color, lines); shading indicates SEM. Immune fractions differ between atretic and non-atretic follicles containing an oocyte (*P<*0.005, Tukey HSD test). Follicle counts by type in **Extended Data Table 4**.

**Extended Data Figure 2.**
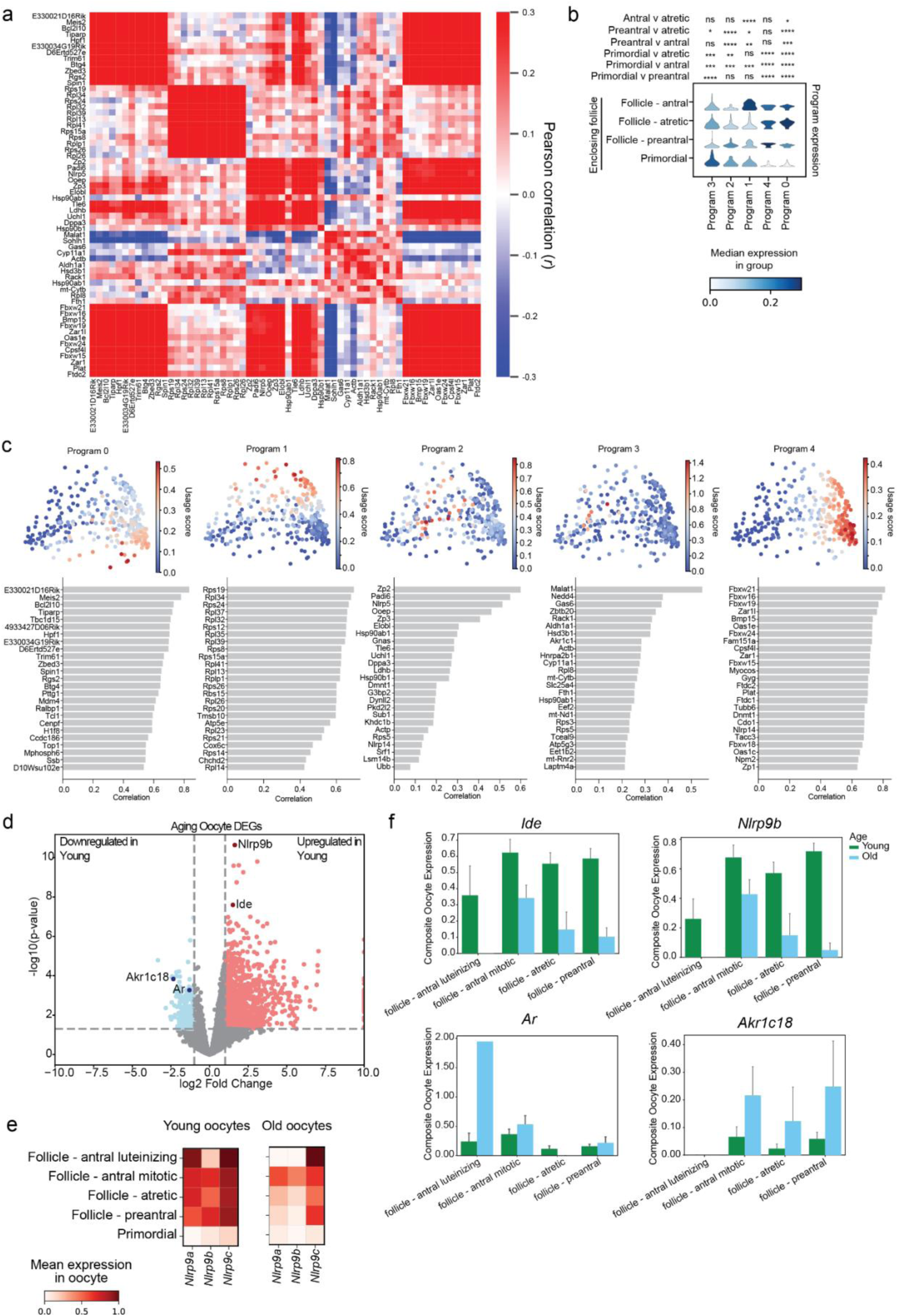
Evaluating oocyte transcriptomes using non-negative matrix factorization **a.** Pearson correlation (red/blue color) of pairwise gene expression across oocytes (rows and columns) based on the top 12 genes from each cNMF program, ordered by hierarchical clustering; *n*=358 oocytes. **b.** Median expression (color) and distribution (kernel density) of oocyte gene program scores (x axis) by enclosing follicle type (y axis); significance assessed by two-tailed tests (\**P*<0.05, \*\**P*<0.01, \*\*\**P*<0.001, \*\*\*\**P*<0.0001; ns). Oocyte counts by enclosing follicle type in **Extended Data Table 4**. **c.** PCA embedding of composite oocyte transcriptomes showing usage of cNMF programs and their top 20 contributing genes; *n*=358 oocytes. **d.** Differentially expressed genes (dots) between young (n=9) and old (n=7) oocytes. Volcano plot showing significance (y axis, -log_10_(P-value)) and log_2_ fold change (x axis). Significant genes (Benjamini-Hochberg FDR<5%, log2FC>1) colored; select genes annotated. Non-primordial oocyte counts by age in **Extended Data Table 4**. **e.** Mean expression (log normalized, variance-scaled) of *Nlrp9*-family genes (columns) in oocytes grouped by the enclosing follicle type (rows), in young (left, *n*=9) and old (right, *n*=7) mice. Oocyte counts by enclosing follicle type and age in **Extended Data Table 4**. **f.** Mean expression of selected differentially expressed genes in oocytes by follicle type for young (*n*=9) and old (*n*=7) mice. Error bars: SEM. Oocyte counts by enclosing follicle type and age in **Extended Data Table 4**.

**Extended Data Figure 3.**
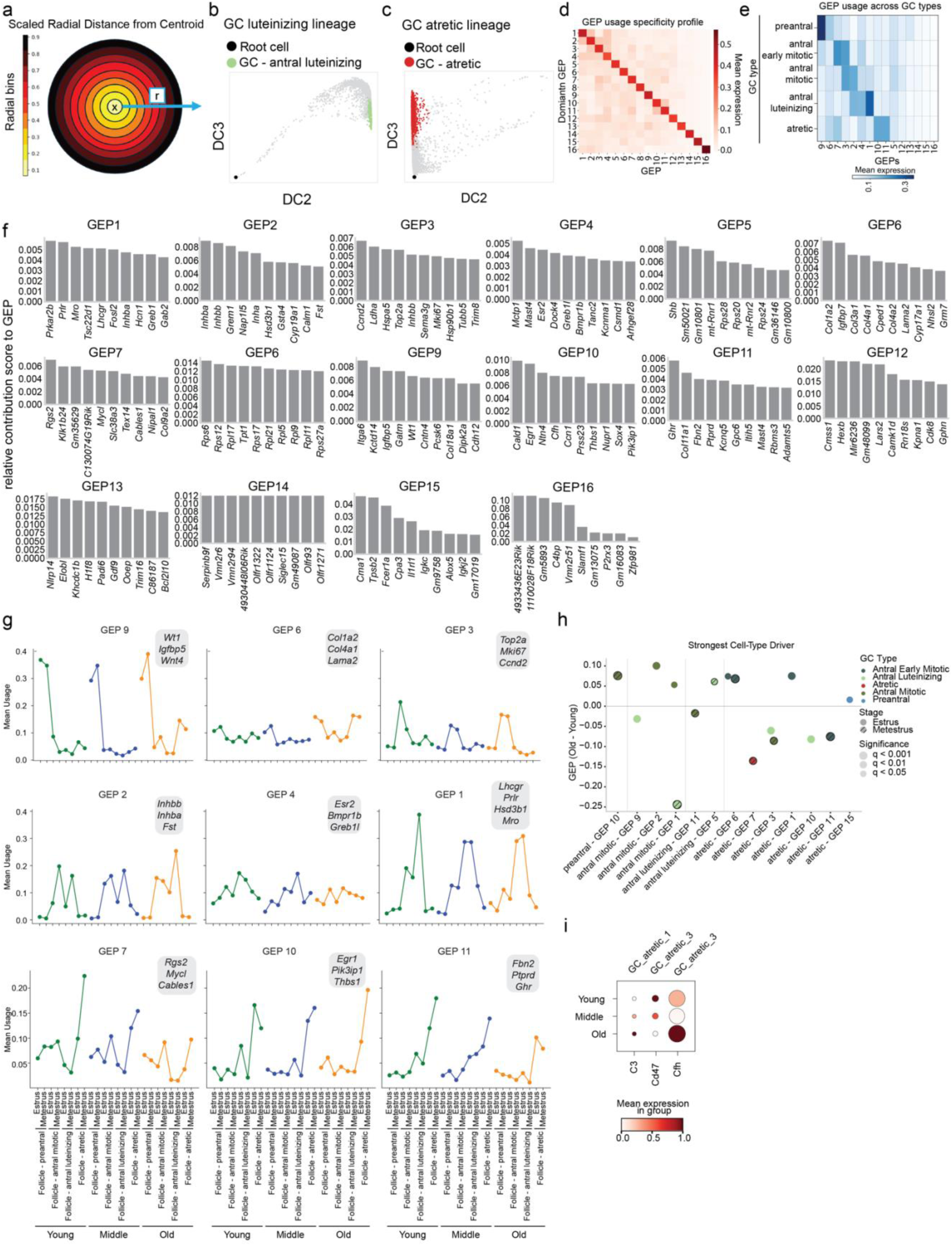
Spatial and molecular determinants of granulosa coordination in follicular niches. **a.** Schematic of radial analysis. Scaled radial distance (*r*) from the follicular centroid (“x”) was computed for each GC spot across 10 uniformly spaced, concentric bins were used to aggregate spot-level characteristics. **b,c.** Diffusion map embeddings (DC2, DC3) of GCs (spots) along luteinizing (**b**) and atretic (**c**) lineages, colored by terminal fate (“GC – antral luteinizing”, “GC – atretic”) and annotated with the root spot (black dot) used for diffusion pseudotime (DPT). **d.** Mean expression (red colorscale; log normalized, variance-scaled) of GC gene programs (rows, columns). GCs were grouped by their dominant program (highest usage); within-group mean program scores are shown. **e.** Mean usage of K=16 cNMF-derived gene expression programs (GEPs; columns) across GC subtypes (rows). Color intensity indicates the mean normalized program usage within each group. **f.** Top contributing genes (*x* axis) for each of the 16 GC GEPs (subplot) and their contribution scores (*y* axis). Contribution scores reflect gene weights within each program. **g.** Mean usage of selected GEPs by follicle type, estrous stage, and age. Each panel (GEP) lists top contributing genes for that GEP and shows the mean GEP usage across preantral, antral, and atretic follicles for young (green), middle (blue), and old (orange) mice. Follicle counts by stage and age in **Extended Data Table 4**. **h.** Dominant GC subtype drivers of age-associated GEP changes. For each significant follicle-level GEP (q < 0.05), the GC subtype with the largest age effect (Old − Young) within each estrous stage is shown. Points represent effect size (β[Age = Old]) for the most strongly associated cell type within the corresponding follicle segment and GEP. Colors denote the dominant GC subtype, while hatch marks distinguish stages (solid = Estrus, hatched = Metestrus). Point size encodes statistical significance level (q < 0.05, 0.01, 0.001). Positive effect sizes indicate higher GEP usage in old ovaries, and negative values indicate higher usage in young. Vertical separators delineate follicle type. **i.** Expression of selected immune-related genes in atretic GC clusters by age. Dot size indicates fraction of expressing cells; color indicates normalized mean expression level.

**Extended Data Figure 4.**
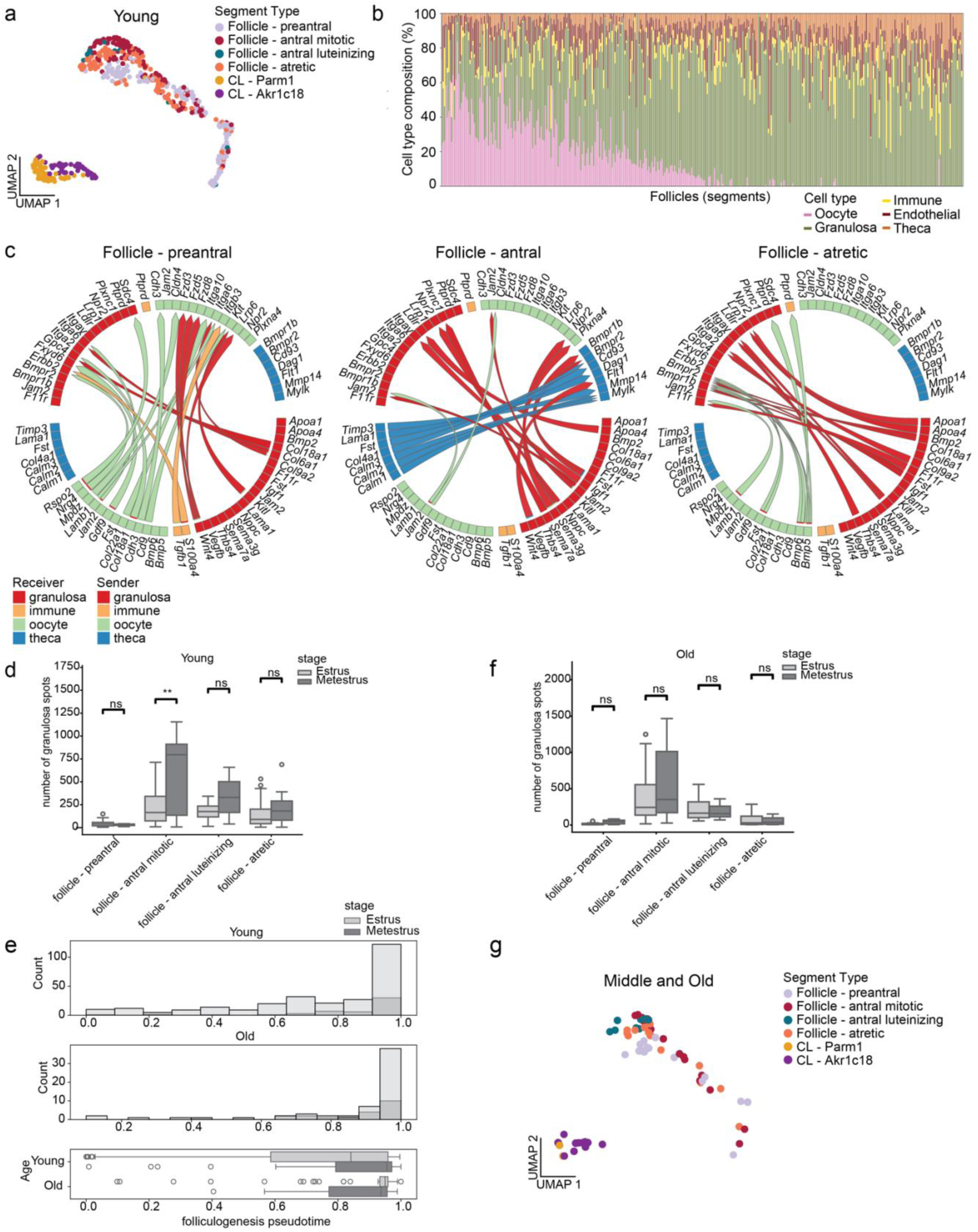
Age-associated temporal dysregulation of folliculogenesis. **a.** UMAP of all young segments (dots; *n*=472 follicles and corpora lutea) based on transport cost distances. **b. c.** Cell type composition (colors, y axis) of each follicle (bars, x axis) ordered by their folliculogenesis diffusion pseudotime as in Fig. 4e. **c.** Circos plot of the top receptor-ligand interactions enriched in preantral (left; *n*=144), antral (center; *n*=114), and atretic (right; *n*=109) follicles identified by MultiNicheNet. Interactions significantly enriched in each follicle type are shown by arrows starting from ligands (bottom half) of a “sender” cell type (color) to the corresponding receptor (top half) of a “receiver” cell type (color). **d.** Distribution of follicle sizes in young mice. Boxplot of the follicle size (number of granulosa spots; y axis) by follicle type (x axis) and stage (bar shading) (\*\**P*<0.005, two-tailed t-test of means). Boxplot: center line indicates the median, box bounds represent the first and third quartiles, and whiskers span from each quartile to the minimum or the maximum (1.5 interquartile range below 25% or above 75% quartiles). Follicle type by stage counts in **Extended Data Table 4**. **e.** Distribution of follicle pseudotime in young and old mice. Boxplot of follicle pseudotime in old and young across stages. Boxplot elements: same as in **d**. Follicle counts by stage and age in **Extended Data Table 4**. **f.** Distribution of follicle sizes in old mice. Boxplot of the follicle size (number of granulosa spots; y axis) by follicle type (x axis) and stage (bar shading) (ns, two-tailed t-test of means). Boxplot elements: same as in **d**. Follicle type by stage counts in **Extended Data Table 4**. **g.** UMAP of all middle and old segments (dots; *n*=190 follicles and corpora lutea) projected into the young segment trajectory based on their transport cost matrix.

**Extended Data Figure 5.**
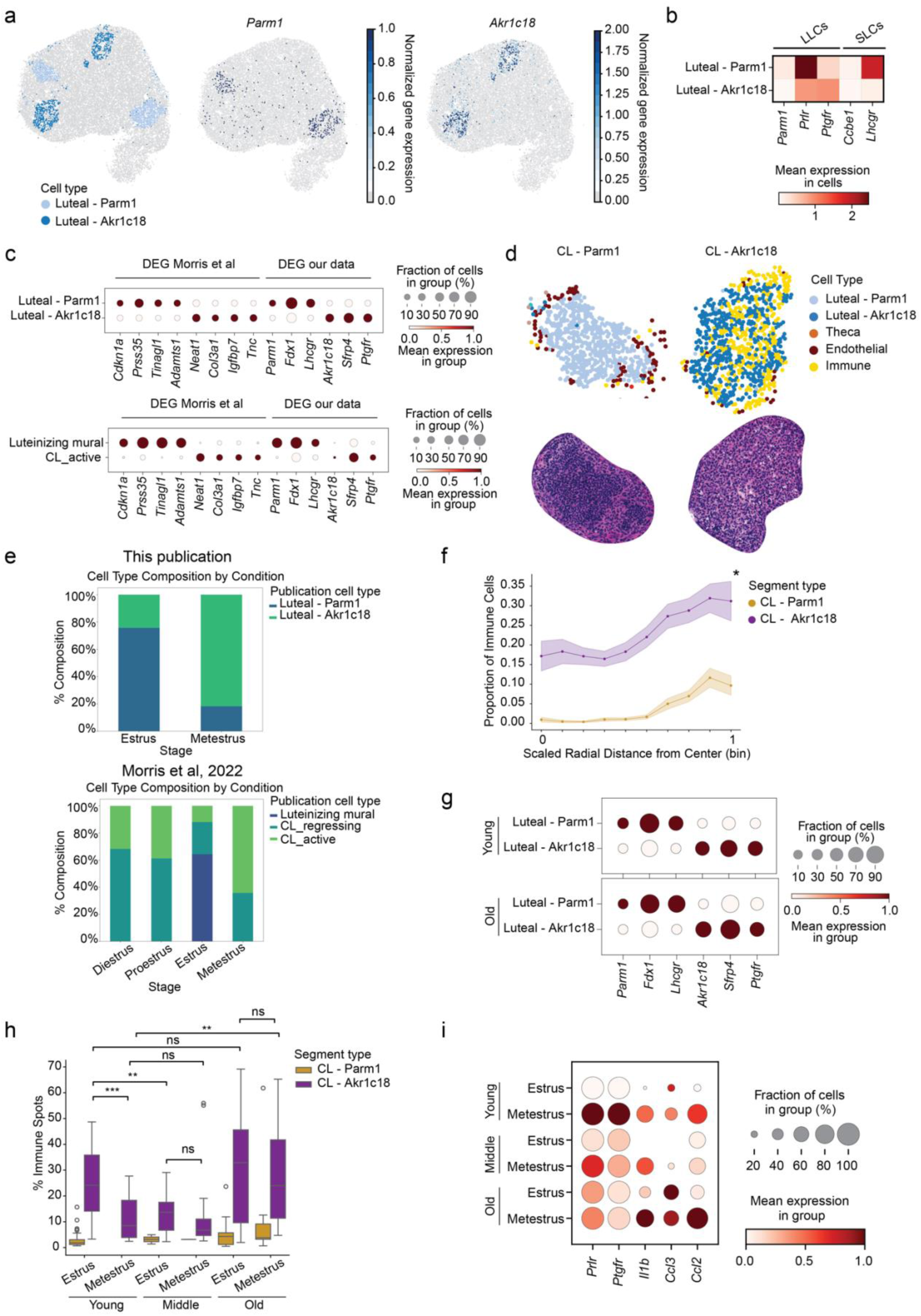
Dysregulation of corpora lutea clearance with age. **a.** Spatial visualization of luteal cell type markers in a representative puck. Top: early (light blue) and late (dark blue) luteal cell clusters. Middle and right: normalized expression of luteal subtype markers *Parm1* and *Akr1c18* (blue scale). All other cell types in gray. **b.** Mean expression of genes associated with large luteal cells (LLCs) and small luteal cells (SLCs) in previous literature. Spot counts in **Extended Data Table 4**. **c.** Differentially expressed genes (DEGs) of luteal clusters from Morris et al. and their corresponding expression in our dataset, showing high concordance between ‘Luteal - Parm1’ and ‘Luteinizing mural,’ and between ‘Luteal - Akr1c18’ and ‘Active CL.’ **d.** Representative spatial plots and H&E images of ‘CL - Parm1’ and ‘CL - Akr1c18’ structures. **e.** Relative proportions of luteal subtypes across estrus and metestrus in our dataset (top) compared with all 4 estrous stages in Morris et al. 2022^37^ (bottom). **f.** Immune cell proportions within each CL type (colored lines, y axis) as a function of the radial distance (x axis) from the CL center (0) to edge (1). Shaded lines represent the standard error of the mean (SEM) fraction across segments within each radial bin (*P<*0.005, Tukey HSD, all bins). CL type counts in **Extended Data Table 4**. **g.** Age-stratified gene expression showing conservation of luteal subtype marker expression in young (top) and old (bottom) ovaries, demonstrating that transcriptional identities are preserved with age despite shifts in proportional abundance. **h.** Immune cell abundance in CL subtypes. Boxplots show distribution of spot counts (y axis, log_10_ scale) of luteal (early and late combined), immune, and vascular cell types (color) across ‘CL – Parm1’ and ‘CL – Akr1c18’ (\*\**P*<0.01, \*\*\**P*<0.001, ns – not significant; two-tailed t-test). Center line indicates median, box bounds represent first and third quartiles, whiskers span from each quartile to the minimum or the maximum (1.5 interquartile range below 25% or above 75% quartiles). CL type counts by age and stage in **Extended Data Table 4**. **i.** Mean expression and proportion of CL - Akr1c18 segments (dot size) expressing select CL clearance and immune-recruitment genes (x-axis), grouped by age and stage (rows). CL – Akr1c18 counts by age and stage in **Extended Data Table 4**.

**Extended Data Figure 6.**
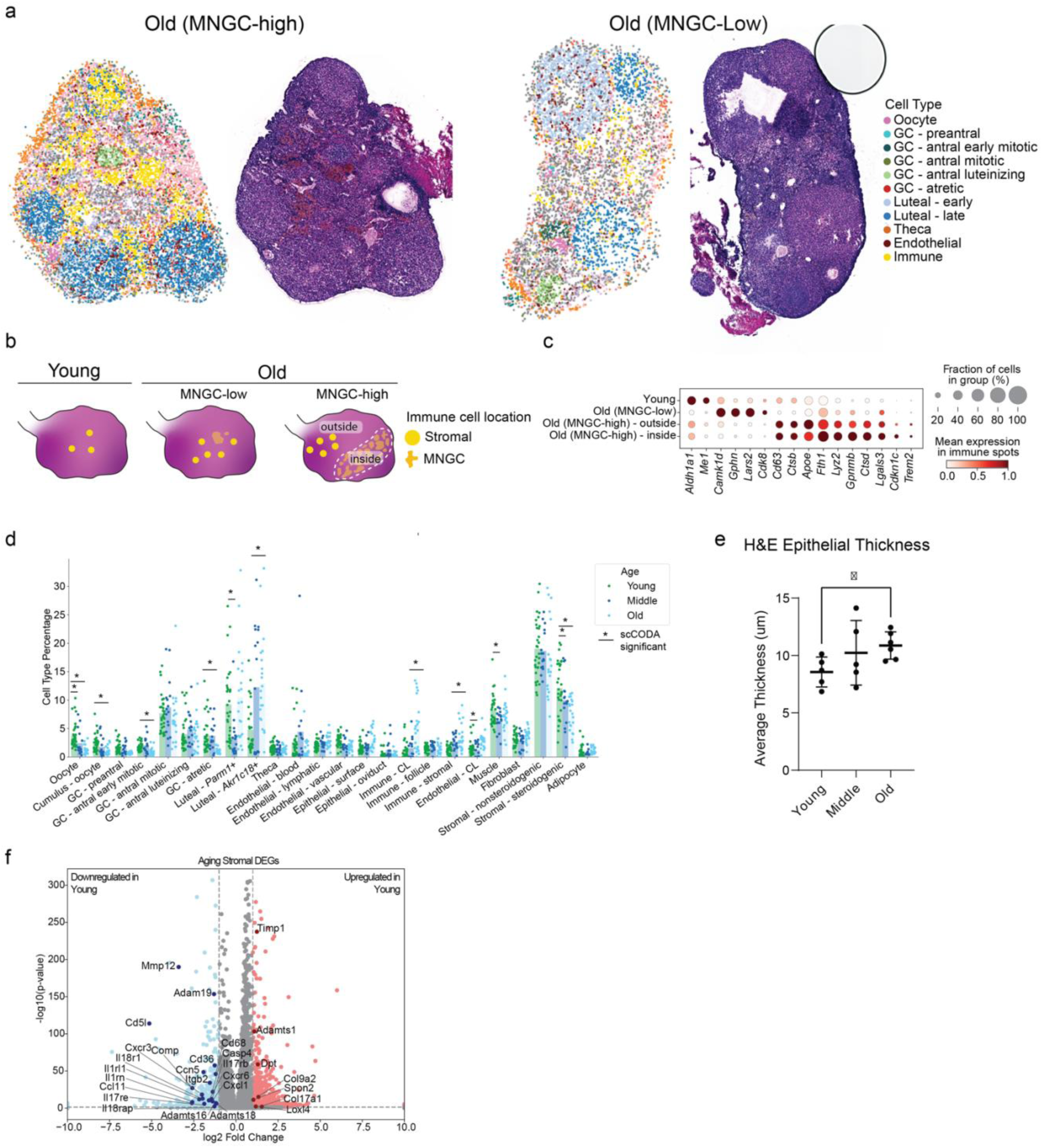
MNGC signatures in aging ovaries. **a.** Representative spatial plots and adjacent H&E sections for old animals categorized as ‘MNGC-high’ and ‘MNGC-low’. **b.** Schematic of MNGC transcriptional analysis. Stromal immune spots were grouped by age (young *vs* old) and MNGC content (low *vs* high), then examined for differentially expressed genes (DEGs) across these four categories. Mean expression (log normalized counts, dot color) and proportion of expressing stromal immune cells (dot size) of differentially expressed genes (columns) in each age-MNGC category (rows). **c.** DEGs in aging stromal cells (fibroblasts, stromal nonsteroidogenic, stromal steroidogenic). Significance (y axis, -log_10_(P-value)) and effect size as log_2_-fold change (x axis) for genes (dots) that are differentially expressed between young and old. Significant genes (color; Benjamini-Hochberg FDR<5%, log2FC>1) are annotated with select genes. Spot counts in **Extended Data Table 4**. **d.** Distribution of cell type composition (y axis) across pucks (dot) and the mean composition across pucks (bars) within each age group (color). Significant changes determined by scCODA indicated by black bars (FDR=0.05). Young *n*= 32 pucks, Middle *n*=16 pucks, Old *n*=21 pucks. **e.** Average surface epithelium thickness in adjacent H&E sections of young, middle, and old animals. Dots represent an H&E section, where three thickness measurements were taken and averaged. Bars represent mean, error bars SEM (\**P*<0.05, two-tailed t-test of means); *n*=5 pucks per age group. **f.** Differentially expressed genes in aging stromal cells (fibroblasts, stromal nonsteroidogenic, stromal steroidogenic). Significance (y axis, -log_10_(P-value)) and effect size as log_2_-fold change (x axis) for genes (dots) that are differentially expressed between young and old. Significant genes (color; Benjamini-Hochberg FDR<5%, log2FC>1) are annotated with select genes. Spot counts in **Extended Data Table 4**.

**Extended Data Figure 7.**
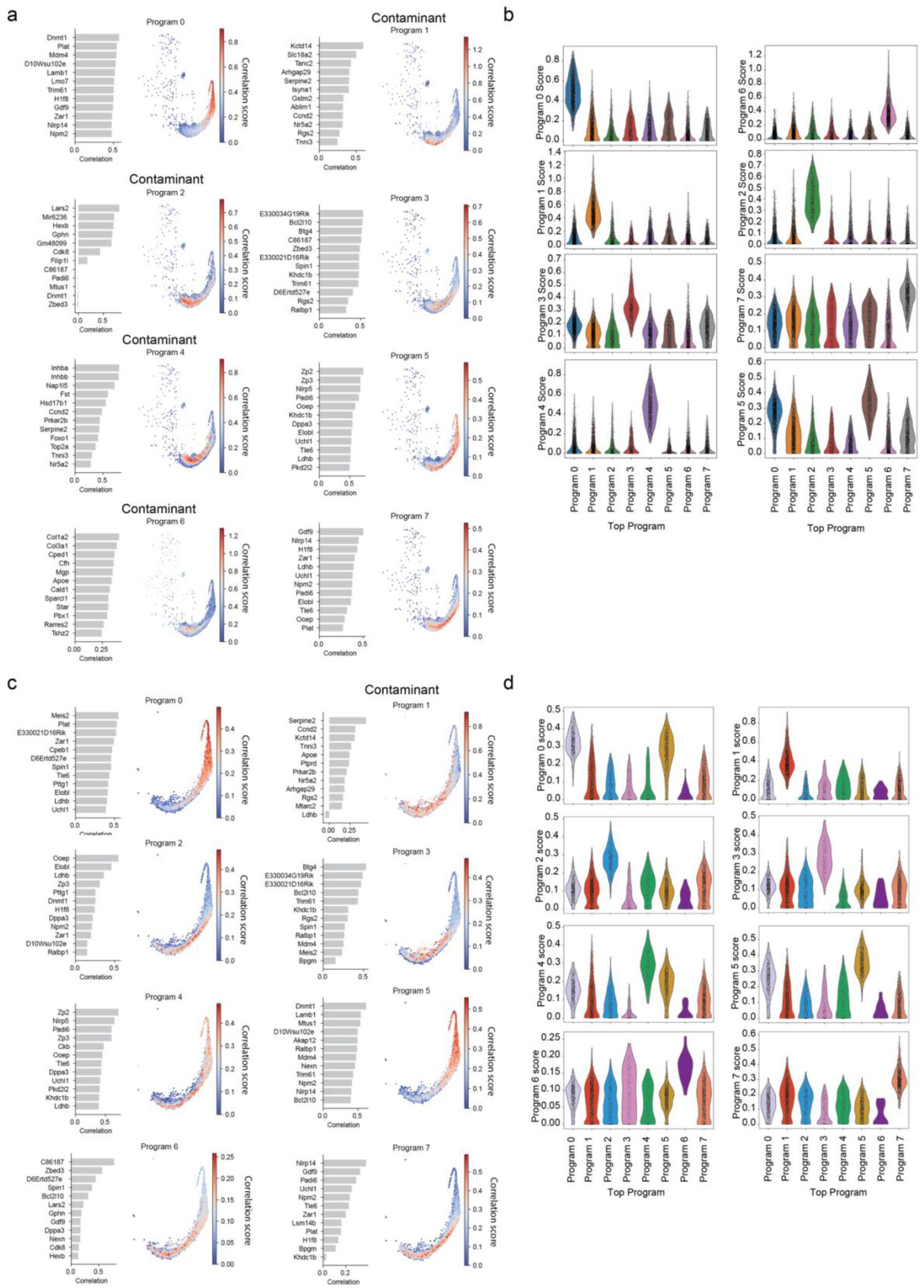
Identifying contaminated oocyte spots using non-negative matrix factorization. Two iterations of oocyte spot decontamination. **a-d.** Identifying contaminated oocyte spots through a two-step iteration, with the first iteration in **a,b** to remove contaminant spots, followed by a second iteration in **c,d** to further remove residual contaminants. **a,c.** Gene programs identified through NMF on oocyte spots using *k*=8 programs. Correlation scores (left, x axis) of the top 12 genes (left, y axis) and their program usage on a UMAP projection (right) of oocyte spots. Programs identified as contamination based on the expression of known cumulus and granulosa marker genes are labeled as ‘Contaminant’ (first iteration: programs 1, 2, 4, and 6; second iteration: program 1). **b,d.** Distribution of program scores (y axis, kernel density estimate) grouped by the top gene program (x axis) for each spot (dots). Distributions were used to manually set program-specific thresholds for removing contaminants. First iteration thresholds: Program 1: > 0.4, Program 2: > 0.3, Program 4: > 0.3, Program 6: > 0.4. Second iteration thresholds: Program 1: > 0.3.

