## Extended Data Table 1 for "Aging disrupts spatiotemporal coordination in the cycling murine ovary"

| Batch | Age | Mouse # | 4-Oct | 6-Oct | 9-Oct | 10-Oct | 11-Oct | 12-Oct | 13-Oct | 14-Oct | 16-Oct | 17-Oct | 18-Oct | 19-Oct | 20-Oct | 21-Oct | 22-Oct | 23-Oct | 24-Oct | 25-Oct | Final Harvest | Harvest Stage |
|---|---|---|---|---|---|---|---|---|---|---|---|---|---|---|---|---|---|---|---|---|---|---|
| Broad 2 | Old | 1 | M/D | M/D | M | M | M/D | D | D | P | E | | | | | | | | | | 16-Oct | E |
| | Old | 2 | E | M | M | M | M | D | D | D | P/E | E | E | M | | | | | | | 19-Oct | M |
| | Old | 3 | E | E/M | E | E | M | D | P | E | | | | | | | | | | | 14-Oct | E |
| | Old | 4 | E | M | E | E | M | D | D | D | M/D | D | D/P | rest | D/P | D/P | P | E | E | M | 21-Oct | M |
| | Young | 5 | E | M | E | M/D | D/P | P/E | P/E | E | | | | | | | | | | | 14-Oct | E |
| | Young | 6 | P | E | E/M | M | M | D | P | D | P | P/E | E | E | rest | M | | | | | 21-Oct | M |
| | Young | 7 | E | E/M | E | E/M | D | D | P/E | E/M | E | | | | | | | | | | 16-Oct | E |
| | Young | 8 | E | E/M | E | E | M | D | P | E | | | | | | | | | | | 14-Oct | E |
| | Middle | 9 | E | M | E | E/M | D | D | D/P | P | M/D | D | D | rest | D/P | E | | | | | 21-Oct | E |
| | Middle | 10 | | E/M | P | E | E/M | D | D | D/P | M/D | D | D/P | rest | D/P | P | P | E | E | M | 25-Oct | M |
| | Middle | 11 | M? | E/M | D | D/P | P/E | P/E | E | E | E | E/M | M | | | | | | | | 18-Oct | M |
| | Middle | 12 | | M | M | M | D | D | D/P | D | P/E | E | | | | | | | | | 17-Oct | E |
| Broad 1 | | | 25-Aug | 26-Aug | 27-Aug | 28-Aug | | | | | | | | | | | | | | | | |
| | Young | 13 | P | E | | | | | | | | | | | | | | | | | 26-Aug | E |
| | Young | 16 | E | | | | | | | | | | | | | | | | | | 25-Aug | E |
| | Middle | 18 | | | | E | | | | | | | | | | | | | | | 28-Aug | E |
| | Middle | 19 | P | E | | | | | | | | | | | | | | | | | 26-Aug | E |
| | Old | 21 | E | | | | | | | | | | | | | | | | | | 25-Aug | E |
| | Old | 25 | E | rest | rest | M | | | | | | | | | | | | | | | 28-Aug | M |
| | | | 20-Dec | 30-Mar | 4-Apr | 10-Apr | 20-Apr | | | | | | | | | | | | | | | |
| Buck | Young | AK01 | | M | M | E | | | | | | | | | | | | | | | | |
| | Young | AK02 | | E | M | D | E | | | | | | | | | | | | | | | |
| | Old | F12 | E | | | | | | | | | | | | | | | | | | | |
| | | | 11-Nov | 12-Nov | 13-Nov | 14-Nov | 15-Nov | 18-Nov | 19-Nov | 20-Nov | 21-Nov | 22-Nov | 25-Nov | 26-Nov | 27-Nov | 28-Nov | 29-Nov | 30-Nov | 1-Dec | | | |
| Yale | Young | 1.1 | M | D | P/E | E | M | E | E | M | | | | | | | | | | | 20-Nov | M |
| | Young | 1.2 | P | P | E | E | M | M/D | D/P | D/P | P/E | | | | | | | | | | 21-Nov | P/E |
| | Young | 1.3 | P/E | P/E | E | E | M | M | M/D | D/P | D/P | P | E | E/M | | | | | | | 26-Nov | E/M |
| | Young | 1.4 | E | E | M | D | D/P | M | M | D/P | D/P | P/E | M | D | P | E | E | M | D | | 1-Dec | D |
| | Middle | 2.1 | D/P | E | E | E | E | M/D | M/D | D | D/P | D/P | P/E | E | E | M | D | | | | 29-Nov | D |
| | Middle | 2.2 | P | E | E | E | M | D | D | D/P | D/P | D | P/E | P/E | E | | | | | | 27-Nov | E |
| | Middle | 2.3 | E | E | M | M | M | D | D/P | D/P | D/P | P | P/E | M | | | | | | | 26-Nov | M |
| | Middle | 2.4 | P/E | E | E | M | M/D | D | D | D/P | D/P | P/E | D/P | D/P | P | | | | | | 27-Nov | P |
| | | | 21-Jan | 22-Jan | 23-Jan | 24-Jan | 25-Jan | 26-Jan | 27-Jan | 28-Jan | 3-Feb | 4-Feb | 5-Feb | 6-Feb | 7-Feb | 8-Feb | | | | | | |
| | Young | O1 | D/P | P/E | E | E | M | D | P | | | | | | | | | | | | 27-Jan | P |
| | Young | O2 | M | M | M | D | P | P | E | M | E | E | M | D/P | | | | | | | 6-Feb | D/P |
| | Young | O3 | E | E | M | D | P | E | E | E | P | PE | M | D | P | E | | | | | 8-Feb | E |
| | Young | O4 | DP | DP | PE | PE | E | E | M | | | | | | | | | | | | 27-Jan | M |
