## Extended Data Table 2 for "Aging disrupts spatiotemporal coordination in the cycling murine ovary"

| Puck_alignment_run | Puck_name | Sample | Sample_puck_name | Mouse |
|---|---|---|---|---|
| 2023-06-20_Puck_230223_01 | Puck_230223_01 | AK01b | AK01b_230223_01 | AK01b |
| 2023-06-20_Puck_230406_01 | Puck_230406_01 | AK01b | AK01b_230406_01 | AK01b |
| 2023-06-20_Puck_230406_06 | Puck_230406_06 | AK01b | AK01b_230406_06 | AK01b |
| 2023-06-20_Puck_230406_08 | Puck_230406_08 | AK01b | AK01b_230406_08 | AK01b |
| 2023-06-20_Puck_230517_23 | Puck_230517_23 | F12ov | F12ov_230517_23 | F12ov |
| 2023-06-20_Puck_230517_33 | Puck_230517_33 | F12ov | F12ov_230517_33 | F12ov |
| 2023-06-20_Puck_230517_36 | Puck_230517_36 | F12ov | F12ov_230517_36 | F12ov |
| 2023-06-20_Puck_230517_37 | Puck_230517_37 | AK02b | AK02b_230517_37 | AK02b |
| 2023-06-20_Puck_230517_38 | Puck_230517_38 | AK02b | AK02b_230517_38 | AK02b |
| 2023-06-22_Puck_230517_39 | Puck_230517_39 | AK02b | AK02b_230517_39 | AK02b |
| 2023-09-05_Puck_230714_23 | Puck_230714_23 | 13YE | 13YE_Puck_230714_2[illegible] | 13 |
| 2023-09-05_Puck_230714_28 | Puck_230714_28 | 13YE | 13YE_Puck_230714_2[illegible] | 13 |
| 2023-09-05_Puck_230714_29 | Puck_230714_29 | 16YM | 16YM_Puck_230714_2[illegible] | 16 |
| 2023-09-05_Puck_230714_39 | Puck_230714_39 | 18ME | 18ME_Puck_230714_3[illegible] | 18 |
| 2023-09-27_Puck_230810_02 | Puck_230810_02 | 19ME | 19ME_230810_02 | 19 |
| 2023-09-27_Puck_230810_06 | Puck_230810_06 | 21OE | 21OE_230810_06 | 21 |
| 2023-09-27_Puck_230810_08 | Puck_230810_08 | 25OM | 25OM_230810_08 | 25 |
| 2023-09-27_Puck_230810_09 | Puck_230810_09 | 19ME | 19ME_230810_09 | 19 |
| 2023-09-27_Puck_230810_11 | Puck_230810_11 | 25OM | 25OM_230810_11 | 25 |
| 2023-11-28_A0029_035 | A0029_035 | 3OE | 3OE_A0029_035 | 3 |
| 2023-11-28_A0029_036 | A0029_036 | 8YE | 8YE_A0029_036 | 8 |
| 2023-11-28_A0029_043 | A0029_043 | 7YE | 7YE_A0029_043 | 7 |
| 2023-11-28_A0064_037 | A0064_037 | 10MM | 10MM_A0064_037 | 10 |
| 2023-11-28_A0064_038 | A0064_038 | 10MM | 10MM_A0064_038 | 10 |
| 2023-11-28_A0064_039 | A0064_039 | 6YM | 6YM_A0064_039 | 6 |
| 2023-11-28_A0064_042 | A0064_042 | 11MM | 11MM_A0064_042 | 11 |
| 2023-11-28_PM104_004 | PM104_004 | 8YE | 8YE_PM104_004 | 8 |
| 2023-11-28_Puck_230913_07 | Puck_230913_07 | 5YE | 5YE_230913_07 | 5 |
| 2023-12-17_A0029_020 | A0029_020 | 9ME | 9ME_A0029_020 | 9 |
| 2023-12-17_A0029_026 | A0029_026 | 12ME | 12ME_A0029_026 | 12 |
| 2023-12-17_A0029_028 | A0029_028 | 2OM | 2OM_A0029_028 | 2 |
| 2023-12-17_A0029_042 | A0029_042 | 7YE | 7YE_A0029_042 | 7 |
| 2023-12-17_A0029_047 | A0029_047 | 5YE | 5YE_A0029_047 | 5 |
| 2023-12-17_A0064_045 | A0064_045 | 11MM | 11MM_A0064_045 | 11 |
| 2023-12-17_PM104_001 | PM104_001 | 1OE | 1OE_PM104_001 | 1 |
| 2023-12-17_Puck_230913_06 | Puck_230913_06 | 1OE | 1OE_Puck_230913_06 | 1 |
| 2024-02-12_Puck_230807_03 | Puck_230807_03 | 1OE | 1OE_230807_03 | 1 |
| 2024-02-12_Puck_240108_03 | Puck_240108_03 | 1OE | 1OE_240108_03 | 1 |
| 2024-02-12_Puck_240108_10 | Puck_240108_10 | 8YE | 8YE_240108_10 | 8 |
| 2024-02-12_Puck_240108_11 | Puck_240108_11 | 8YE | 8YE_240108_11 | 8 |
| 2024-02-12_Puck_240108_14 | Puck_240108_14 | 3OE | 3OE_240108_14 | 3 |
| 2024-02-12_Puck_240108_16 | Puck_240108_16 | 3OE | 3OE_240108_16 | 3 |
| 2024-02-12_Puck_240108_17 | Puck_240108_17 | 3OE | 3OE_240108_17 | 3 |

| | | | | |
|---|---|---|---|---|
| 2024-02-13_Puck_230807_04 | Puck_230807_04 | 8YE | 8YE_230807_04 | 8 |
| 2024-02-13_Puck_240108_07 | Puck_240108_07 | 1OE | 1OE_240108_07 | 1 |
| 2024-02-27_Puck_240108_20 | Puck_240108_20 | 5YE | 5YE_240108_20 | 5 |
| 2024-02-27_Puck_240108_24 | Puck_240108_24 | 5YE | 5YE_240108_24 | 5 |
| 2024-02-27_Puck_240108_25 | Puck_240108_25 | 5YE | 5YE_240108_25 | 5 |
| 2024-02-27_Puck_240108_26 | Puck_240108_26 | 5YE | 5YE_240108_26 | 5 |
| 2024-02-27_Puck_240108_27 | Puck_240108_27 | 6YM | 6YM_240108_27 | 6 |
| 2024-02-27_Puck_240108_32 | Puck_240108_32 | 6YM | 6YM_240108_32 | 6 |
| 2024-02-27_Puck_240108_33 | Puck_240108_33 | 6YM | 6YM_240108_33 | 6 |
| 2024-02-27_Puck_240108_35 | Puck_240108_35 | 6YM | 6YM_240108_35 | 6 |
| 2024-02-27_Puck_240129_21 | Puck_240129_21 | 2OM | 2OM_240129_21 | 2 |
| 2024-02-27_Puck_240129_22 | Puck_240129_22 | 2OM | 2OM_240129_22 | 2 |
| 2024-02-27_Puck_240129_23 | Puck_240129_23 | 2OM | 2OM_240129_23 | 2 |
| 2024-03-02_Puck_240129_24 | Puck_240129_24 | 10MM | 10MM_240129_24 | 10 |
| 2024-03-02_Puck_240129_25 | Puck_240129_25 | 10MM | 10MM_240129_25 | 10 |
| 2024-03-02_Puck_240129_26 | Puck_240129_26 | 10MM | 10MM_240129_26 | 10 |
| 2024-03-02_Puck_240129_27 | Puck_240129_27 | 4OM | 4OM_240129_27 | 4 |
| 2024-03-02_Puck_240129_29 | Puck_240129_29 | 4OM | 4OM_240129_29 | 4 |
| 2024-03-02_Puck_240129_31 | Puck_240129_31 | 11MM | 11MM_240129_31 | 11 |
| 2024-03-02_Puck_240129_32 | Puck_240129_32 | 11MM | 11MM_240129_32 | 11 |
| 2024-03-02_Puck_240129_36 | Puck_240129_36 | 7YE | 7YE_240129_36 | 7 |
| 2024-03-02_Puck_240129_37 | Puck_240129_37 | 7YE | 7YE_240129_37 | 7 |
| 2024-03-04_Puck_240129_33 | Puck_240129_33 | 11MM | 11MM_240129_33 | 11 |
| 2024-03-04_Puck_240129_35 | Puck_240129_35 | 11MM | 11MM_240129_35 | 11 |
| 2024-03-21_Puck_230807_27 | Puck_230807_27 | 7YE | 7YE_230807_27 | 7 |
| 2025-03-13_Puck_230913_14 | Puck_230913_14 | 2.6YM | 26YM_250313_04 | 2.6 |

| Age | Stage | Mouse Batch | Puck Batch | MNGCs |
|---|---|---|---|---|
| Young | Estrus | Buck | V8 | |
| Young | Estrus | Buck | V8 | |
| Young | Estrus | Buck | V8 | |
| Young | Estrus | Buck | V8 | |
| Old | Estrus | Buck | V8 | visible |
| Old | Estrus | Buck | V8 | visible |
| Old | Estrus | Buck | V8 | visible |
| Young | Estrus | Buck | V8 | none |
| Young | Estrus | Buck | V8 | none |
| Young | Estrus | Buck | V8 | none |
| Young | Estrus | Broad1 | V8 | |
| Young | Estrus | Broad1 | V8 | |
| Young | Metestrus | Broad1 | V8 | |
| Middle | Estrus | Broad1 | V8 | |
| Middle | Estrus | Broad1 | V8 | visible |
| Old | Estrus | Broad1 | V8 | visible |
| Old | Metestrus | Broad1 | V8 | visible |
| Middle | Estrus | Broad1 | V8 | possible |
| Old | Metestrus | Broad1 | V8 | visible |
| Old | Estrus | Broad2 | Curio | visible |
| Young | Estrus | Broad2 | Curio | none |
| Young | Estrus | Broad2 | Curio | none |
| Middle | Metestrus | Broad2 | Curio | possible |
| Middle | Metestrus | Broad2 | Curio | |
| Young | Metestrus | Broad2 | Curio | none |
| Middle | Metestrus | Broad2 | Curio | none |
| Young | Estrus | Broad2 | Curio | none |
| Young | Estrus | Broad2 | V8 | none |
| Middle | Estrus | Broad2 | Curio | possible |
| Middle | Estrus | Broad2 | Curio | none |
| Old | Metestrus | Broad2 | Curio | visible |
| Young | Estrus | Broad2 | Curio | none |
| Young | Estrus | Broad2 | Curio | none |
| Middle | Metestrus | Broad2 | Curio | possible |
| Old | Estrus | Broad2 | Curio | visible |
| Old | Estrus | Broad2 | V8 | visible |
| Old | Estrus | Broad2 | V11 | visible |
| Old | Estrus | Broad2 | V11 | visible |
| Young | Estrus | Broad2 | V11 | none |
| Young | Estrus | Broad2 | V11 | none |
| Old | Estrus | Broad2 | V11 | visible |
| Old | Estrus | Broad2 | V11 | visible |
| Old | Estrus | Broad2 | V11 | possible |

| | | | | |
|---|---|---|---|---|
| Young | Estrus | Broad2 | V11 | none |
| Old | Estrus | Broad2 | V11 | visible |
| Young | Estrus | Broad2 | V11 | none |
| Young | Estrus | Broad2 | V11 | possible |
| Young | Estrus | Broad2 | V11 | none |
| Young | Estrus | Broad2 | V11 | none |
| Young | Metestrus | Broad2 | V11 | none |
| Young | Metestrus | Broad2 | V11 | none |
| Young | Metestrus | Broad2 | V11 | possible |
| Young | Metestrus | Broad2 | V11 | possible |
| Old | Metestrus | Broad2 | V11 | visible |
| Old | Metestrus | Broad2 | V11 | visible |
| Old | Metestrus | Broad2 | V11 | visible |
| Middle | Metestrus | Broad2 | V11 | possible |
| Middle | Metestrus | Broad2 | V11 | visible |
| Middle | Metestrus | Broad2 | V11 | possible |
| Old | Metestrus | Broad2 | V11 | visible |
| Old | Metestrus | Broad2 | V11 | visible |
| Middle | Metestrus | Broad2 | V11 | possible |
| Middle | Metestrus | Broad2 | V11 | visible |
| Young | Estrus | Broad2 | V11 | none |
| Young | Estrus | Broad2 | V11 | none |
| Middle | Metestrus | Broad2 | V11 | visible |
| Middle | Metestrus | Broad2 | V11 | possible |
| Young | Estrus | Broad2 | V11 | possible |
| Young | Metestrus | Yale | V8 | none |
