## Extended Data Table 3 for "Aging disrupts spatiotemporal coordination in the cycling murine ovary"

| group | names | scores | logfoldchang | pvals | pvals_adj |
|---|---|---|---|---|---|
| Oocyte | Nlrp14 | 138.57684 | 7.7596674 | 0 | 0 |
| Oocyte | H1f8 | 135.11133 | 8.09594 | 0 | 0 |
| Oocyte | Khdc1b | 134.12206 | 7.939358 | 0 | 0 |
| Oocyte | Gdf9 | 134.03876 | 7.675353 | 0 | 0 |
| Oocyte | Elobl | 130.0231 | 8.028568 | 0 | 0 |
| Oocyte | Padi6 | 122.224846 | 7.2938905 | 0 | 0 |
| Oocyte | Ldhb | 120.28983 | 4.3392134 | 0 | 0 |
| Oocyte | Bpgm | 115.822426 | 5.7837625 | 0 | 0 |
| Oocyte | Bcl2l10 | 114.81437 | 7.750009 | 0 | 0 |
| Oocyte | E330034G19Rik | 114.551315 | 7.8326144 | 0 | 0 |
| Oocyte | Ooep | 114.279144 | 7.5795107 | 0 | 0 |
| Oocyte | Uchl1 | 113.13571 | 4.727344 | 0 | 0 |
| Oocyte | Npm2 | 112.835365 | 7.4162664 | 0 | 0 |
| Oocyte | Zbed3 | 112.37182 | 6.5699754 | 0 | 0 |
| Oocyte | Zar1 | 111.44981 | 7.7513843 | 0 | 0 |
| Oocyte | C86187 | 111.089134 | 7.7305946 | 0 | 0 |
| Oocyte | Trim61 | 111.07985 | 7.791505 | 0 | 0 |
| Oocyte | Dnmt1 | 110.49408 | 5.378522 | 0 | 0 |
| Oocyte | Btg4 | 109.90709 | 7.738044 | 0 | 0 |
| Oocyte | D10Wsu102e | 109.212 | 5.470587 | 0 | 0 |
| Oocyte | Tle6 | 109.074234 | 5.8108563 | 0 | 0 |
| Oocyte | Plat | 108.37091 | 5.4818826 | 0 | 0 |
| Oocyte | Lsm14b | 106.9101 | 4.8601975 | 0 | 0 |
| Oocyte | Spin1 | 105.829414 | 4.6145573 | 0 | 0 |
| Oocyte | Zp3 | 102.520096 | 7.608677 | 0 | 0 |
| Oocyte | Nlrp5 | 99.82572 | 7.3207736 | 0 | 0 |
| Oocyte | Dppa3 | 99.363976 | 7.4817634 | 0 | 0 |
| Oocyte | Rdx | 98.35037 | 3.292572 | 0 | 0 |
| Oocyte | E330021D16Rik | 97.896126 | 7.723927 | 0 | 0 |
| Oocyte | Zp2 | 97.55553 | 7.52774 | 0 | 0 |
| Oocyte | D6Ertd527e | 96.640755 | 7.6661315 | 0 | 0 |
| Oocyte | Mdm4 | 95.42125 | 3.8648503 | 0 | 0 |
| Oocyte | Pkd2l2 | 93.61737 | 6.699923 | 0 | 0 |
| Oocyte | Ralbp1 | 90.66719 | 3.592547 | 0 | 0 |
| Oocyte | Rgs2 | 88.49591 | 2.8531818 | 0 | 0 |
| Oocyte | Tcl1 | 86.97503 | 7.5227113 | 0 | 0 |
| Oocyte | Tacc3 | 86.49844 | 5.041936 | 0 | 0 |
| Oocyte | Fbxw24 | 84.97516 | 7.389504 | 0 | 0 |
| Oocyte | Smc4 | 82.74213 | 3.1220324 | 0 | 0 |
| Oocyte | Birc5 | 82.67748 | 4.4921694 | 0 | 0 |
| Oocyte | Mphosph6 | 82.4105 | 5.182598 | 0 | 0 |
| Oocyte | Oas1e | 82.40819 | 7.5612483 | 0 | 0 |

| | | | | | |
|---|---|---|---|---|---|
| Oocyte | Oas1c | 81.08471 | 7.045629 | 0 | 0 |
| Oocyte | Pttg1 | 80.1759 | 4.1173615 | 0 | 0 |
| Oocyte | Ftdc2 | 79.012856 | 7.357887 | 0 | 0 |
| Oocyte | Skp1 | 78.44528 | 2.5488536 | 0 | 0 |
| Oocyte | 4933427D06Rik | 78.22448 | 7.060156 | 0 | 0 |
| Oocyte | Cenpf | 78.06711 | 3.920782 | 0 | 0 |
| Oocyte | Tet3 | 76.70717 | 3.8793397 | 0 | 0 |
| Oocyte | G3bp2 | 74.400955 | 2.7784424 | 0 | 0 |
| Cumulus - oocyte | Nlrp14 | 37.862267 | 3.7233071 | 0 | 0 |
| Cumulus - oocyte | Gdf9 | 32.090218 | 3.457762 | 6.04E-226 | 2.35E-222 |
| Cumulus - oocyte | H1f8 | 31.72521 | 3.5890296 | 6.98E-221 | 2.38E-217 |
| Cumulus - oocyte | Khdc1b | 30.987469 | 3.5521703 | 7.95E-211 | 1.97E-207 |
| Cumulus - oocyte | Elobl | 29.519753 | 3.577476 | 1.61E-191 | 3.65E-188 |
| Cumulus - oocyte | Padi6 | 28.327158 | 3.4210894 | 1.60E-176 | 3.36E-173 |
| Cumulus - oocyte | Ldhb | 28.2838 | 1.6778482 | 5.47E-176 | 1.07E-172 |
| Cumulus - oocyte | Rgs2 | 26.737812 | 1.3302977 | 1.71E-157 | 2.46E-154 |
| Cumulus - oocyte | Serpine2 | 25.761976 | 1.141802 | 2.37E-146 | 2.94E-143 |
| Cumulus - oocyte | Bcl2l10 | 23.704372 | 3.4697587 | 3.25E-124 | 2.86E-121 |
| Cumulus - oocyte | Ooep | 22.941534 | 3.380773 | 1.79E-116 | 1.48E-113 |
| Cumulus - oocyte | E330034G19Rik | 22.037012 | 3.365433 | 1.27E-107 | 9.14E-105 |
| Cumulus - oocyte | Npm2 | 21.306675 | 3.1971607 | 9.84E-101 | 6.72E-98 |
| Cumulus - oocyte | Btg4 | 21.1354 | 3.3973775 | 3.76E-99 | 2.50E-96 |
| Cumulus - oocyte | Zbed3 | 21.080246 | 2.8318288 | 1.21E-98 | 7.85E-96 |
| Cumulus - oocyte | C86187 | 20.8093 | 3.3783762 | 3.57E-96 | 2.26E-93 |
| Cumulus - oocyte | Bpgm | 20.79263 | 2.2787554 | 5.05E-96 | 3.13E-93 |
| Cumulus - oocyte | Uchl1 | 20.669922 | 1.6913954 | 6.46E-95 | 3.92E-92 |
| Cumulus - oocyte | Dnmt1 | 20.0943 | 2.0825527 | 8.28E-90 | 4.71E-87 |
| Cumulus - oocyte | Kctd14 | 19.927752 | 1.5640953 | 2.34E-88 | 1.30E-85 |
| Cumulus - oocyte | Trim61 | 19.533169 | 3.2194579 | 5.74E-85 | 3.13E-82 |
| Cumulus - oocyte | D10Wsu102e | 19.323635 | 2.2060473 | 3.40E-83 | 1.72E-80 |
| Cumulus - oocyte | Tle6 | 19.272646 | 2.368876 | 9.12E-83 | 4.44E-80 |
| Cumulus - oocyte | Lsm14b | 19.046103 | 1.8451849 | 7.08E-81 | 3.33E-78 |
| Cumulus - oocyte | Plat | 18.529346 | 2.0958529 | 1.20E-76 | 5.27E-74 |
| Cumulus - oocyte | Zar1 | 18.042263 | 3.0741348 | 9.07E-73 | 3.81E-70 |
| Cumulus - oocyte | Nlrp5 | 17.351902 | 3.173918 | 1.91E-67 | 7.78E-65 |
| Cumulus - oocyte | Spin1 | 16.997982 | 1.596622 | 8.50E-65 | 3.27E-62 |
| Cumulus - oocyte | E330021D16Rik | 16.536419 | 3.191036 | 2.01E-61 | 7.50E-59 |
| Cumulus - oocyte | Smc4 | 16.25359 | 1.1388115 | 2.11E-59 | 7.67E-57 |
| Cumulus - oocyte | Dppa3 | 16.219685 | 3.1791472 | 3.66E-59 | 1.32E-56 |
| Cumulus - oocyte | Ccnd2 | 16.182312 | 1.0092741 | 6.72E-59 | 2.35E-56 |
| Cumulus - oocyte | Zp2 | 15.903568 | 3.055164 | 5.99E-57 | 2.07E-54 |
| Cumulus - oocyte | Zp3 | 15.864555 | 2.9775925 | 1.12E-56 | 3.76E-54 |
| Cumulus - oocyte | D6Ertd527e | 15.058323 | 3.134122 | 3.04E-51 | 9.34E-49 |

| | | | | | |
|---|---|---|---|---|---|
| Cumulus - oocyte | Birc5 | 14.338953 | 1.8564986 | 1.25E-46 | 3.34E-44 |
| Cumulus - oocyte | Pkd2l2 | 13.421602 | 2.7068586 | 4.52E-41 | 1.10E-38 |
| Cumulus - oocyte | Cenpf | 13.410044 | 1.5207646 | 5.28E-41 | 1.28E-38 |
| Cumulus - oocyte | mt-Rnr1 | 13.127324 | 0.30671492 | 2.30E-39 | 5.40E-37 |
| Cumulus - oocyte | Mdm4 | 12.81313 | 1.0903937 | 1.38E-37 | 3.15E-35 |
| Cumulus - oocyte | Top2a | 12.806562 | 1.2231499 | 1.51E-37 | 3.40E-35 |
| Cumulus - oocyte | Tacc3 | 12.711658 | 1.9281787 | 5.09E-37 | 1.12E-34 |
| Cumulus - oocyte | Ralbp1 | 12.703961 | 1.0607136 | 5.62E-37 | 1.22E-34 |
| Cumulus - oocyte | Npm1 | 11.8678665 | 0.5126431 | 1.74E-32 | 3.37E-30 |
| Cumulus - oocyte | Tnni3 | 11.608393 | 0.8144207 | 3.74E-31 | 6.75E-29 |
| Cumulus - oocyte | Inhbb | 11.591727 | 0.91446215 | 4.54E-31 | 8.15E-29 |
| Cumulus - oocyte | Rbbp7 | 11.457436 | 0.59458214 | 2.16E-30 | 3.80E-28 |
| Cumulus - oocyte | Ivns1abp | 11.420017 | 0.69829756 | 3.32E-30 | 5.81E-28 |
| Cumulus - oocyte | Tubb5 | 11.327781 | 0.5152863 | 9.56E-30 | 1.65E-27 |
| Cumulus - oocyte | Tcl1 | 11.224647 | 2.8282166 | 3.09E-29 | 5.27E-27 |
| GC - preantral | Kctd14 | 92.02202 | 4.9467297 | 0 | 0 |
| GC - preantral | Slc18a2 | 64.00603 | 5.125036 | 0 | 0 |
| GC - preantral | Rgs2 | 58.256195 | 2.6725667 | 0 | 0 |
| GC - preantral | Tenm4 | 56.218266 | 2.581112 | 0 | 0 |
| GC - preantral | Serpine2 | 55.278725 | 2.3850675 | 0 | 0 |
| GC - preantral | Isyna1 | 45.389824 | 2.8079946 | 0 | 0 |
| GC - preantral | Arhgap29 | 45.31936 | 2.9277802 | 0 | 0 |
| GC - preantral | Tanc2 | 45.12575 | 2.7382746 | 0 | 0 |
| GC - preantral | Itga6 | 41.163704 | 3.620754 | 0 | 0 |
| GC - preantral | Gm2044 | 40.377747 | 3.5685387 | 0 | 0 |
| GC - preantral | Igfbp5 | 38.40136 | 3.299891 | 0 | 0 |
| GC - preantral | Tnni3 | 38.061756 | 2.1875615 | 0 | 0 |
| GC - preantral | Ldhb | 37.957363 | 2.1290429 | 0 | 0 |
| GC - preantral | Wt1 | 35.628197 | 3.3439558 | 5.13E-278 | 6.67E-275 |
| GC - preantral | Ablim1 | 35.232563 | 2.3516667 | 6.35E-272 | 7.88E-269 |
| GC - preantral | Syne2 | 35.229195 | 1.4746648 | 7.15E-272 | 8.48E-269 |
| GC - preantral | Ivns1abp | 34.95912 | 1.780117 | 9.41E-268 | 1.07E-264 |
| GC - preantral | Ccnd2 | 33.032513 | 1.8061237 | 2.77E-239 | 2.70E-236 |
| GC - preantral | Dag1 | 32.492542 | 1.787986 | 1.36E-231 | 1.28E-228 |
| GC - preantral | St3gal5 | 31.787188 | 1.7268206 | 9.73E-222 | 8.86E-219 |
| GC - preantral | Rap2b | 30.858429 | 2.9311874 | 4.32E-209 | 3.68E-206 |
| GC - preantral | Emx2 | 30.788418 | 4.3838086 | 3.74E-208 | 3.10E-205 |
| GC - preantral | Vcan | 30.62694 | 2.5136874 | 5.36E-206 | 4.30E-203 |
| GC - preantral | Tut4 | 29.657316 | 1.6737617 | 2.73E-193 | 2.13E-190 |
| GC - preantral | Amhr2 | 29.645607 | 2.5453794 | 3.86E-193 | 2.93E-190 |
| GC - preantral | Gatm | 29.17937 | 3.1013432 | 3.54E-187 | 2.61E-184 |
| GC - preantral | Nr5a2 | 27.915636 | 1.4275843 | 1.72E-171 | 1.24E-168 |
| GC - preantral | Aff3 | 27.84045 | 2.5899284 | 1.41E-170 | 9.84E-168 |

| | | | | | |
|---|---|---|---|---|---|
| GC - preantral | Gtf2a1 | 27.613789 | 2.8850198 | 7.60E-168 | 5.19E-165 |
| GC - preantral | Tubb5 | 27.585785 | 1.1988851 | 1.65E-167 | 1.10E-164 |
| GC - preantral | Myo10 | 26.029972 | 1.9638733 | 2.27E-149 | 1.38E-146 |
| GC - preantral | Dpf3 | 25.749504 | 2.578361 | 3.26E-146 | 1.94E-143 |
| GC - preantral | Arhgap31 | 25.487759 | 2.590524 | 2.69E-143 | 1.57E-140 |
| GC - preantral | Npm1 | 25.308317 | 1.0689498 | 2.59E-141 | 1.44E-138 |
| GC - preantral | Myo1b | 24.747917 | 2.1624675 | 3.26E-135 | 1.78E-132 |
| GC - preantral | Gstm2 | 24.632366 | 1.312443 | 5.69E-134 | 3.04E-131 |
| GC - preantral | Mras | 24.481968 | 2.042135 | 2.30E-132 | 1.21E-129 |
| GC - preantral | Hsp90ab1 | 24.257544 | 0.7666055 | 5.50E-130 | 2.84E-127 |
| GC - preantral | Foxl2 | 24.239038 | 1.7629167 | 8.63E-130 | 4.36E-127 |
| GC - preantral | Gls | 24.161463 | 1.7518662 | 5.66E-129 | 2.81E-126 |
| GC - preantral | Rpl4 | 24.080826 | 0.8011575 | 3.97E-128 | 1.94E-125 |
| GC - preantral | Igf1r | 23.974836 | 1.454651 | 5.09E-127 | 2.40E-124 |
| GC - preantral | Sytl4 | 23.71676 | 2.61964 | 2.42E-124 | 1.12E-121 |
| GC - preantral | Fus | 23.714287 | 0.9007561 | 2.57E-124 | 1.17E-121 |
| GC - preantral | Pcsk6 | 23.190395 | 3.813756 | 5.69E-119 | 2.55E-116 |
| GC - preantral | Gas6 | 23.110325 | 0.99128836 | 3.65E-118 | 1.61E-115 |
| GC - preantral | Hmgcs2 | 22.509027 | 1.1702616 | 3.39E-112 | 1.42E-109 |
| GC - preantral | Nnat | 22.329903 | 2.8512137 | 1.89E-110 | 7.83E-108 |
| GC - preantral | Synpo | 21.924551 | 1.7636058 | 1.52E-106 | 6.08E-104 |
| GC - preantral | Ncl | 21.774952 | 0.869449 | 4.01E-105 | 1.59E-102 |
| GC - antral early mitotic | Serpine2 | 111.67321 | 4.331063 | 0 | 0 |
| GC - antral early mitotic | Rgs2 | 94.48482 | 3.803297 | 0 | 0 |
| GC - antral early mitotic | Inha | 89.92043 | 3.381332 | 0 | 0 |
| GC - antral early mitotic | Hspa5 | 82.54195 | 2.7899487 | 0 | 0 |
| GC - antral early mitotic | Ccnd2 | 82.07996 | 3.497613 | 0 | 0 |
| GC - antral early mitotic | Gja1 | 80.65014 | 3.0232227 | 0 | 0 |
| GC - antral early mitotic | Slc18a2 | 74.75208 | 5.232702 | 0 | 0 |
| GC - antral early mitotic | Ctsl | 73.50867 | 2.9056509 | 0 | 0 |
| GC - antral early mitotic | Ivns1abp | 73.497406 | 3.1202261 | 0 | 0 |
| GC - antral early mitotic | Hsp90b1 | 71.8132 | 2.2725244 | 0 | 0 |
| GC - antral early mitotic | Tenm4 | 70.53179 | 2.8818953 | 0 | 0 |
| GC - antral early mitotic | Tubb5 | 68.81417 | 2.5283232 | 0 | 0 |
| GC - antral early mitotic | Nr5a2 | 63.43372 | 2.6744235 | 0 | 0 |
| GC - antral early mitotic | Kctd14 | 62.51869 | 3.391245 | 0 | 0 |
| GC - antral early mitotic | Foxo1 | 62.16405 | 2.7416484 | 0 | 0 |
| GC - antral early mitotic | Npm1 | 58.896725 | 2.14097 | 0 | 0 |
| GC - antral early mitotic | Pdia6 | 56.619965 | 2.2934215 | 0 | 0 |
| GC - antral early mitotic | Top2a | 56.212578 | 3.2794523 | 0 | 0 |
| GC - antral early mitotic | Cyb5a | 55.518368 | 2.136932 | 0 | 0 |
| GC - antral early mitotic | Tut4 | 55.36597 | 2.5260007 | 0 | 0 |
| GC - antral early mitotic | Ncl | 54.221245 | 1.9006169 | 0 | 0 |

| | | | | | |
|---|---|---|---|---|---|
| GC - antral early mitotic | Tanc2 | 53.504063 | 2.7657359 | 0 | 0 |
| GC - antral early mitotic | Cnmd | 53.11756 | 3.3649836 | 0 | 0 |
| GC - antral early mitotic | Map1b | 52.440384 | 2.2947102 | 0 | 0 |
| GC - antral early mitotic | Rps5 | 52.189278 | 1.5753896 | 0 | 0 |
| GC - antral early mitotic | Rpl4 | 51.644638 | 1.5806428 | 0 | 0 |
| GC - antral early mitotic | Tuba1b | 51.50555 | 2.305075 | 0 | 0 |
| GC - antral early mitotic | Scd2 | 51.39899 | 1.9723318 | 0 | 0 |
| GC - antral early mitotic | Carhsp1 | 50.54568 | 2.8601892 | 0 | 0 |
| GC - antral early mitotic | Hnrnpab | 50.532406 | 1.81982 | 0 | 0 |
| GC - antral early mitotic | Pdia3 | 50.46718 | 1.8808961 | 0 | 0 |
| GC - antral early mitotic | Ano4 | 50.145706 | 4.4269443 | 0 | 0 |
| GC - antral early mitotic | Mras | 49.436428 | 3.0079958 | 0 | 0 |
| GC - antral early mitotic | Hnrnpa2b1 | 48.571083 | 1.6609243 | 0 | 0 |
| GC - antral early mitotic | Fst | 48.25111 | 2.4193811 | 0 | 0 |
| GC - antral early mitotic | Fam13a | 47.84372 | 2.756897 | 0 | 0 |
| GC - antral early mitotic | Tshz1 | 47.84005 | 2.3295426 | 0 | 0 |
| GC - antral early mitotic | Rbbp7 | 47.313553 | 1.9023976 | 0 | 0 |
| GC - antral early mitotic | Ptbp3 | 46.327614 | 2.4692056 | 0 | 0 |
| GC - antral early mitotic | Tnni3 | 45.597694 | 2.263213 | 0 | 0 |
| GC - antral early mitotic | Calr | 45.345257 | 1.7545683 | 0 | 0 |
| GC - antral early mitotic | Arhgap29 | 45.23688 | 2.630122 | 0 | 0 |
| GC - antral early mitotic | Ank3 | 45.083885 | 3.4659722 | 0 | 0 |
| GC - antral early mitotic | Ranbp1 | 44.65644 | 2.0550113 | 0 | 0 |
| GC - antral early mitotic | Sfpq | 44.36217 | 1.832469 | 0 | 0 |
| GC - antral early mitotic | Smc4 | 44.126926 | 2.2482011 | 0 | 0 |
| GC - antral early mitotic | Hsp90ab1 | 43.843624 | 1.3390768 | 0 | 0 |
| GC - antral early mitotic | Txnl1 | 43.691135 | 2.1613028 | 0 | 0 |
| GC - antral early mitotic | Apoa4 | 43.36369 | 3.6799242 | 0 | 0 |
| GC - antral early mitotic | Htra1 | 43.031647 | 2.2046273 | 0 | 0 |
| GC - antral mitotic | Inhba | 235.8038 | 5.381864 | 0 | 0 |
| GC - antral mitotic | Inha | 234.6913 | 4.219585 | 0 | 0 |
| GC - antral mitotic | Serpine2 | 228.79854 | 4.304275 | 0 | 0 |
| GC - antral mitotic | Inhbb | 212.59544 | 5.431075 | 0 | 0 |
| GC - antral mitotic | Nap1l5 | 193.07161 | 4.8636518 | 0 | 0 |
| GC - antral mitotic | Gja1 | 191.08249 | 3.4380753 | 0 | 0 |
| GC - antral mitotic | Prkar2b | 168.57257 | 3.311303 | 0 | 0 |
| GC - antral mitotic | Fst | 162.1494 | 3.9925952 | 0 | 0 |
| GC - antral mitotic | Hspa5 | 152.42531 | 2.4356747 | 0 | 0 |
| GC - antral mitotic | Hsd17b1 | 144.54456 | 4.0299892 | 0 | 0 |
| GC - antral mitotic | Tnni3 | 144.34163 | 3.4331348 | 0 | 0 |
| GC - antral mitotic | Ccnd2 | 143.48074 | 3.2419353 | 0 | 0 |
| GC - antral mitotic | Cyb5a | 141.07938 | 2.605162 | 0 | 0 |
| GC - antral mitotic | Hsp90b1 | 137.7603 | 1.9833899 | 0 | 0 |

| | | | | | |
|---|---|---|---|---|---|
| GC - antral mitotic | Foxo1 | 128.93433 | 2.8522542 | 0 | 0 |
| GC - antral mitotic | Nr5a2 | 125.30609 | 2.661765 | 0 | 0 |
| GC - antral mitotic | Tubb5 | 109.7225 | 2.0061834 | 0 | 0 |
| GC - antral mitotic | Top2a | 109.11523 | 3.451202 | 0 | 0 |
| GC - antral mitotic | Jak1 | 104.2108 | 2.2382078 | 0 | 0 |
| GC - antral mitotic | Socs2 | 103.63872 | 2.4571862 | 0 | 0 |
| GC - antral mitotic | Calm1 | 103.48919 | 1.7327436 | 0 | 0 |
| GC - antral mitotic | Fam13a | 103.01721 | 3.0944934 | 0 | 0 |
| GC - antral mitotic | Ctsl | 98.85503 | 1.9819456 | 0 | 0 |
| GC - antral mitotic | Ivns1abp | 97.28776 | 2.136324 | 0 | 0 |
| GC - antral mitotic | Map1b | 96.12268 | 2.1452832 | 0 | 0 |
| GC - antral mitotic | Trib2 | 95.37835 | 2.3220909 | 0 | 0 |
| GC - antral mitotic | Pdia6 | 90.50775 | 1.8639292 | 0 | 0 |
| GC - antral mitotic | Scd2 | 89.48118 | 1.7002639 | 0 | 0 |
| GC - antral mitotic | Mid1ip1 | 89.12681 | 2.4411027 | 0 | 0 |
| GC - antral mitotic | Hnrnpab | 88.91579 | 1.542981 | 0 | 0 |
| GC - antral mitotic | Htra1 | 88.17139 | 2.3118744 | 0 | 0 |
| GC - antral mitotic | Tut4 | 86.002464 | 2.0538375 | 0 | 0 |
| GC - antral mitotic | Hnrnpa2b1 | 84.827225 | 1.3449802 | 0 | 0 |
| GC - antral mitotic | Tuba1b | 83.95703 | 1.9111165 | 0 | 0 |
| GC - antral mitotic | Akr1c14 | 82.84276 | 2.2709813 | 0 | 0 |
| GC - antral mitotic | Ext1 | 82.78666 | 1.9737055 | 0 | 0 |
| GC - antral mitotic | St3gal4 | 82.60229 | 2.3573034 | 0 | 0 |
| GC - antral mitotic | Calr | 82.51841 | 1.5720153 | 0 | 0 |
| GC - antral mitotic | Tenm4 | 81.26572 | 1.714403 | 0 | 0 |
| GC - antral mitotic | Smc4 | 79.33759 | 2.0946 | 0 | 0 |
| GC - antral mitotic | Tceal9 | 77.62618 | 1.244258 | 0 | 0 |
| GC - antral mitotic | Cnmd | 76.610695 | 2.8233426 | 0 | 0 |
| GC - antral mitotic | Pdia3 | 75.88776 | 1.4003567 | 0 | 0 |
| GC - antral mitotic | Rgs2 | 75.71393 | 1.6189079 | 0 | 0 |
| GC - antral mitotic | Bmpr2 | 74.86105 | 1.8478587 | 0 | 0 |
| GC - antral mitotic | Obsl1 | 73.06354 | 2.2153866 | 0 | 0 |
| GC - antral mitotic | Cyp19a1 | 71.05652 | 3.3666153 | 0 | 0 |
| GC - antral mitotic | Csrp2 | 70.889565 | 1.9542636 | 0 | 0 |
| GC - antral mitotic | Mctp1 | 67.32655 | 2.68258 | 0 | 0 |
| GC - antral mitotic | Tox2 | 66.90588 | 3.1286318 | 0 | 0 |
| GC - antral luteinizing | Inhba | 230.95398 | 6.683851 | 0 | 0 |
| GC - antral luteinizing | Prkar2b | 217.0511 | 5.1954923 | 0 | 0 |
| GC - antral luteinizing | Inha | 214.73003 | 4.8096523 | 0 | 0 |
| GC - antral luteinizing | Gja1 | 203.17473 | 4.4585967 | 0 | 0 |
| GC - antral luteinizing | Nr5a2 | 177.5845 | 4.3348207 | 0 | 0 |
| GC - antral luteinizing | Serpine2 | 175.44627 | 4.068784 | 0 | 0 |
| GC - antral luteinizing | Fst | 151.98297 | 4.3087955 | 0 | 0 |

| | | | | | |
|---|---|---|---|---|---|
| GC - antral luteinizing | Foxo1 | 147.13487 | 3.7276232 | 0 | 0 |
| GC - antral luteinizing | Trib2 | 147.1004 | 3.8814077 | 0 | 0 |
| GC - antral luteinizing | Nap1l5 | 144.47357 | 4.1946316 | 0 | 0 |
| GC - antral luteinizing | Cyb5a | 137.67822 | 3.0483382 | 0 | 0 |
| GC - antral luteinizing | Hsd17b1 | 136.11241 | 4.224363 | 0 | 0 |
| GC - antral luteinizing | Tnni3 | 133.247 | 3.631725 | 0 | 0 |
| GC - antral luteinizing | Inhbb | 132.99641 | 4.0143356 | 0 | 0 |
| GC - antral luteinizing | Ext1 | 129.55579 | 3.4723647 | 0 | 0 |
| GC - antral luteinizing | Fam13a | 124.99301 | 4.054291 | 0 | 0 |
| GC - antral luteinizing | Bmpr2 | 124.62182 | 3.3465722 | 0 | 0 |
| GC - antral luteinizing | Greb1 | 121.97891 | 2.983712 | 0 | 0 |
| GC - antral luteinizing | Socs2 | 121.9508 | 3.253474 | 0 | 0 |
| GC - antral luteinizing | Jak1 | 116.16634 | 2.8793283 | 0 | 0 |
| GC - antral luteinizing | Mid1ip1 | 112.149025 | 3.3776016 | 0 | 0 |
| GC - antral luteinizing | Tsc22d1 | 112.02571 | 2.011179 | 0 | 0 |
| GC - antral luteinizing | Cyp19a1 | 109.97819 | 4.955878 | 0 | 0 |
| GC - antral luteinizing | St3gal4 | 106.81892 | 3.3466227 | 0 | 0 |
| GC - antral luteinizing | Plxnc1 | 106.2097 | 3.9608104 | 0 | 0 |
| GC - antral luteinizing | Sema5a | 104.78826 | 3.1950276 | 0 | 0 |
| GC - antral luteinizing | Crim1 | 104.51305 | 3.5397398 | 0 | 0 |
| GC - antral luteinizing | Greb1l | 102.78335 | 3.2950654 | 0 | 0 |
| GC - antral luteinizing | Hspa5 | 98.83115 | 2.0563009 | 0 | 0 |
| GC - antral luteinizing | Calm1 | 98.34921 | 2.048312 | 0 | 0 |
| GC - antral luteinizing | Mro | 98.196175 | 4.492536 | 0 | 0 |
| GC - antral luteinizing | Ivns1abp | 96.1435 | 2.476744 | 0 | 0 |
| GC - antral luteinizing | Me2 | 95.31968 | 3.0715275 | 0 | 0 |
| GC - antral luteinizing | St3gal5 | 94.66595 | 2.5033383 | 0 | 0 |
| GC - antral luteinizing | Bmpr1b | 91.90276 | 3.3531845 | 0 | 0 |
| GC - antral luteinizing | Wapl | 91.70346 | 2.5369442 | 0 | 0 |
| GC - antral luteinizing | Tanc2 | 90.58887 | 2.8570256 | 0 | 0 |
| GC - antral luteinizing | Map1b | 90.25386 | 2.3595095 | 0 | 0 |
| GC - antral luteinizing | Srbd1 | 89.38602 | 3.6830118 | 0 | 0 |
| GC - antral luteinizing | Hsp90b1 | 88.81635 | 1.7732698 | 0 | 0 |
| GC - antral luteinizing | Fat1 | 88.02708 | 3.3747146 | 0 | 0 |
| GC - antral luteinizing | Slc26a7 | 86.83709 | 4.2918057 | 0 | 0 |
| GC - antral luteinizing | Obsl1 | 86.4943 | 2.9107263 | 0 | 0 |
| GC - antral luteinizing | Gab2 | 85.5669 | 2.5245957 | 0 | 0 |
| GC - antral luteinizing | Cnmd | 85.54356 | 3.4122422 | 0 | 0 |
| GC - antral luteinizing | Mctp1 | 84.43837 | 3.5773587 | 0 | 0 |
| GC - antral luteinizing | Akr1c14 | 83.440704 | 2.6377053 | 0 | 0 |
| GC - antral luteinizing | Csrp2 | 83.34424 | 2.6147032 | 0 | 0 |
| GC - antral luteinizing | Tut4 | 79.98341 | 2.233951 | 0 | 0 |
| GC - antral luteinizing | Fndc3b | 78.80742 | 2.1986938 | 0 | 0 |

| | | | | | |
|---|---|---|---|---|---|
| GC - atretic | Ivns1abp | 121.68908 | 3.7619798 | 0 | 0 |
| GC - atretic | Tenm4 | 106.650566 | 3.2117853 | 0 | 0 |
| GC - atretic | Ptprd | 105.55792 | 3.2220578 | 0 | 0 |
| GC - atretic | Foxo1 | 104.54444 | 3.3309078 | 0 | 0 |
| GC - atretic | Rgs2 | 102.41105 | 3.1097207 | 0 | 0 |
| GC - atretic | Grb14 | 101.19847 | 3.116863 | 0 | 0 |
| GC - atretic | Prss23 | 94.672066 | 4.0333953 | 0 | 0 |
| GC - atretic | Serpine2 | 93.9858 | 2.7276905 | 0 | 0 |
| GC - atretic | Ctsl | 92.436485 | 2.6853414 | 0 | 0 |
| GC - atretic | Tdrd5 | 89.130806 | 3.8642044 | 0 | 0 |
| GC - atretic | Mast4 | 88.761024 | 3.1879692 | 0 | 0 |
| GC - atretic | Gja1 | 88.3886 | 2.455628 | 0 | 0 |
| GC - atretic | Syne2 | 81.44591 | 2.1997766 | 0 | 0 |
| GC - atretic | Nr5a2 | 79.10611 | 2.5345526 | 0 | 0 |
| GC - atretic | Kcnq5 | 74.93492 | 4.604549 | 0 | 0 |
| GC - atretic | Sema5a | 74.48898 | 2.9000313 | 0 | 0 |
| GC - atretic | Bmpr2 | 73.858185 | 2.5617127 | 0 | 0 |
| GC - atretic | Esr2 | 71.05576 | 3.44445 | 0 | 0 |
| GC - atretic | Inha | 70.799095 | 1.973545 | 0 | 0 |
| GC - atretic | Sox4 | 70.3181 | 2.502486 | 0 | 0 |
| GC - atretic | Bmpr1b | 64.07706 | 2.9832945 | 0 | 0 |
| GC - atretic | Plxdc2 | 64.07054 | 2.3540864 | 0 | 0 |
| GC - atretic | Crim1 | 62.770428 | 2.7845004 | 0 | 0 |
| GC - atretic | Adamts2 | 62.27009 | 3.0589685 | 0 | 0 |
| GC - atretic | Ccn2 | 61.908787 | 4.6260543 | 0 | 0 |
| GC - atretic | Thbs1 | 61.53373 | 3.3916116 | 0 | 0 |
| GC - atretic | Ank3 | 61.436653 | 3.6340134 | 0 | 0 |
| GC - atretic | Arhgef28 | 61.185047 | 3.3878715 | 0 | 0 |
| GC - atretic | Tceal9 | 61.173386 | 1.5106273 | 0 | 0 |
| GC - atretic | Cfh | 59.85932 | 1.9129349 | 0 | 0 |
| GC - atretic | Pik3r1 | 58.684074 | 2.3276062 | 0 | 0 |
| GC - atretic | Akr1c14 | 58.527313 | 2.3149714 | 0 | 0 |
| GC - atretic | Tshz1 | 58.186825 | 2.1796284 | 0 | 0 |
| GC - atretic | Zfp385b | 56.94041 | 3.8400192 | 0 | 0 |
| GC - atretic | Map4k4 | 55.742855 | 2.077838 | 0 | 0 |
| GC - atretic | Tanc2 | 55.616337 | 2.2970219 | 0 | 0 |
| GC - atretic | Prkar2b | 55.46149 | 1.679147 | 0 | 0 |
| GC - atretic | Tut4 | 54.87235 | 1.9623878 | 0 | 0 |
| GC - atretic | Gtf2e2 | 54.58922 | 2.74084 | 0 | 0 |
| GC - atretic | Csmd1 | 54.40376 | 2.3548295 | 0 | 0 |
| GC - atretic | Tmtc2 | 54.336018 | 2.6187787 | 0 | 0 |
| GC - atretic | Cyb5a | 53.858635 | 1.5646852 | 0 | 0 |
| GC - atretic | Fbn2 | 53.57897 | 3.3067532 | 0 | 0 |

| | | | | | |
|---|---|---|---|---|---|
| GC - atretic | Me2 | 53.40572 | 2.2954397 | 0 | 0 |
| GC - atretic | Pde7b | 53.018616 | 3.0991278 | 0 | 0 |
| GC - atretic | Ghr | 52.89546 | 3.0670652 | 0 | 0 |
| GC - atretic | Cnmd | 52.009884 | 2.7676165 | 0 | 0 |
| GC - atretic | Itm2b | 51.53574 | 1.3299278 | 0 | 0 |
| GC - atretic | 8030451A03Rik | 51.18955 | 2.909589 | 0 | 0 |
| GC - atretic | Itih5 | 50.8234 | 2.642921 | 0 | 0 |
| Luteal - Parm1 | Cyp11a1 | 225.83612 | 2.7798793 | 0 | 0 |
| Luteal - Parm1 | Fdx1 | 188.78944 | 2.743119 | 0 | 0 |
| Luteal - Parm1 | Lhcgr | 165.22925 | 3.680088 | 0 | 0 |
| Luteal - Parm1 | Prlr | 158.98679 | 2.8672068 | 0 | 0 |
| Luteal - Parm1 | Idh1 | 158.3661 | 2.9124706 | 0 | 0 |
| Luteal - Parm1 | Scarb1 | 155.01588 | 3.3191059 | 0 | 0 |
| Luteal - Parm1 | Runx2 | 154.7865 | 4.0405307 | 0 | 0 |
| Luteal - Parm1 | Star | 154.30652 | 2.8928435 | 0 | 0 |
| Luteal - Parm1 | Cst8 | 152.99878 | 4.1058064 | 0 | 0 |
| Luteal - Parm1 | Fkbp5 | 146.064 | 2.7352653 | 0 | 0 |
| Luteal - Parm1 | Gm2a | 143.20952 | 3.193982 | 0 | 0 |
| Luteal - Parm1 | Ltbp1 | 139.16104 | 3.2645268 | 0 | 0 |
| Luteal - Parm1 | Heg1 | 133.8226 | 3.3091743 | 0 | 0 |
| Luteal - Parm1 | Hsd17b7 | 130.54662 | 4.3205447 | 0 | 0 |
| Luteal - Parm1 | Sfrp4 | 124.00355 | 2.5521078 | 0 | 0 |
| Luteal - Parm1 | Sgk1 | 123.85697 | 3.7052426 | 0 | 0 |
| Luteal - Parm1 | Ephx2 | 119.867424 | 2.1201415 | 0 | 0 |
| Luteal - Parm1 | Nrn1 | 119.01924 | 3.4762568 | 0 | 0 |
| Luteal - Parm1 | Tsc22d1 | 117.40232 | 1.611382 | 0 | 0 |
| Luteal - Parm1 | Hsd3b1 | 116.607124 | 1.5448831 | 0 | 0 |
| Luteal - Parm1 | Aplp2 | 113.34277 | 2.0641413 | 0 | 0 |
| Luteal - Parm1 | Smarca1 | 113.10965 | 2.5069065 | 0 | 0 |
| Luteal - Parm1 | Sparc | 106.095665 | 1.7494707 | 0 | 0 |
| Luteal - Parm1 | Slc25a30 | 103.19917 | 2.77379 | 0 | 0 |
| Luteal - Parm1 | Nr5a2 | 101.49839 | 2.1738243 | 0 | 0 |
| Luteal - Parm1 | Npc2 | 101.427 | 2.0303195 | 0 | 0 |
| Luteal - Parm1 | Elovl5 | 101.1888 | 2.2169588 | 0 | 0 |
| Luteal - Parm1 | Atp6v0e2 | 101.09766 | 3.1347349 | 0 | 0 |
| Luteal - Parm1 | Aif1l | 100.946205 | 3.4360702 | 0 | 0 |
| Luteal - Parm1 | Scd2 | 100.243385 | 1.8817962 | 0 | 0 |
| Luteal - Parm1 | Hmgcs1 | 97.80588 | 2.2879703 | 0 | 0 |
| Luteal - Parm1 | Acsl4 | 96.00112 | 2.7298322 | 0 | 0 |
| Luteal - Parm1 | Prdx2 | 95.461426 | 1.8501306 | 0 | 0 |
| Luteal - Parm1 | Me1 | 93.56885 | 1.6113523 | 0 | 0 |
| Luteal - Parm1 | Mrfap1 | 91.89915 | 2.1467001 | 0 | 0 |
| Luteal - Parm1 | Acly | 87.92007 | 2.423954 | 0 | 0 |

| | | | | | |
|---|---|---|---|---|---|
| Luteal - Parm1 | Prss35 | 87.52366 | 1.8804314 | 0 | 0 |
| Luteal - Parm1 | Sod1 | 86.47987 | 1.4579163 | 0 | 0 |
| Luteal - Parm1 | Aebp1 | 85.80524 | 1.9052328 | 0 | 0 |
| Luteal - Parm1 | Rora | 85.45372 | 1.8289915 | 0 | 0 |
| Luteal - Parm1 | Fam126a | 83.63355 | 2.53162 | 0 | 0 |
| Luteal - Parm1 | Ctnnb1 | 82.99316 | 1.4918522 | 0 | 0 |
| Luteal - Parm1 | Msi2 | 81.5997 | 1.685373 | 0 | 0 |
| Luteal - Parm1 | Avpi1 | 81.46971 | 2.9005265 | 0 | 0 |
| Luteal - Parm1 | Dhcr7 | 81.46523 | 3.500877 | 0 | 0 |
| Luteal - Parm1 | Cemip | 81.16727 | 2.890587 | 0 | 0 |
| Luteal - Parm1 | Bst2 | 81.030014 | 1.6095892 | 0 | 0 |
| Luteal - Parm1 | Slc6a6 | 80.931404 | 2.6749814 | 0 | 0 |
| Luteal - Parm1 | Akr1cl | 80.43795 | 1.071181 | 0 | 0 |
| Luteal - Parm1 | Prxl2b | 79.77633 | 2.8977115 | 0 | 0 |
| Luteal - Akr1c18 | Sfrp4 | 303.63223 | 5.80309 | 0 | 0 |
| Luteal - Akr1c18 | Ephx2 | 254.54741 | 3.9714992 | 0 | 0 |
| Luteal - Akr1c18 | S100a6 | 215.40668 | 4.3177834 | 0 | 0 |
| Luteal - Akr1c18 | Akr1c18 | 213.27672 | 6.0173335 | 0 | 0 |
| Luteal - Akr1c18 | Malat1 | 195.97409 | 1.4322782 | 0 | 0 |
| Luteal - Akr1c18 | Nupr1 | 193.568 | 4.4007063 | 0 | 0 |
| Luteal - Akr1c18 | Ptgfr | 173.211 | 4.787237 | 0 | 0 |
| Luteal - Akr1c18 | Clu | 170.1842 | 2.4804301 | 0 | 0 |
| Luteal - Akr1c18 | Ybx1 | 159.69658 | 2.2714052 | 0 | 0 |
| Luteal - Akr1c18 | Cyp11a1 | 156.42538 | 2.0029633 | 0 | 0 |
| Luteal - Akr1c18 | Ftl1 | 144.48743 | 1.8735844 | 0 | 0 |
| Luteal - Akr1c18 | Pdia4 | 138.11012 | 3.188327 | 0 | 0 |
| Luteal - Akr1c18 | Tnc | 137.5091 | 4.6746507 | 0 | 0 |
| Luteal - Akr1c18 | Far1 | 124.345856 | 3.4034722 | 0 | 0 |
| Luteal - Akr1c18 | Gadd45a | 121.26407 | 3.46273 | 0 | 0 |
| Luteal - Akr1c18 | Star | 118.194214 | 2.0408874 | 0 | 0 |
| Luteal - Akr1c18 | Ltbp1 | 116.30762 | 2.8149004 | 0 | 0 |
| Luteal - Akr1c18 | Frmd5 | 113.93301 | 2.2560568 | 0 | 0 |
| Luteal - Akr1c18 | Gm2a | 110.11949 | 2.447274 | 0 | 0 |
| Luteal - Akr1c18 | S100a4 | 109.57716 | 4.5475574 | 0 | 0 |
| Luteal - Akr1c18 | Anxa2 | 109.44234 | 2.1907856 | 0 | 0 |
| Luteal - Akr1c18 | AU020206 | 107.856224 | 1.7260654 | 0 | 0 |
| Luteal - Akr1c18 | Lgmn | 107.77001 | 3.4643004 | 0 | 0 |
| Luteal - Akr1c18 | Lipg | 101.181305 | 3.878742 | 0 | 0 |
| Luteal - Akr1c18 | Gpx3 | 100.860596 | 2.2169313 | 0 | 0 |
| Luteal - Akr1c18 | Aplp2 | 99.46232 | 1.7264562 | 0 | 0 |
| Luteal - Akr1c18 | Wnt10b | 94.67194 | 3.3531184 | 0 | 0 |
| Luteal - Akr1c18 | Onecut2 | 93.04961 | 6.2271733 | 0 | 0 |
| Luteal - Akr1c18 | Cd63 | 91.4775 | 1.3842065 | 0 | 0 |

| | | | | | |
|---|---|---|---|---|---|
| Luteal - Akr1c18 | Idh1 | 90.13872 | 1.6493249 | 0 | 0 |
| Luteal - Akr1c18 | Kcnma1 | 87.08397 | 2.4673789 | 0 | 0 |
| Luteal - Akr1c18 | Runx2 | 85.755455 | 2.4306207 | 0 | 0 |
| Luteal - Akr1c18 | Neat1 | 85.51368 | 1.5430443 | 0 | 0 |
| Luteal - Akr1c18 | Fkbp5 | 84.38572 | 1.5803432 | 0 | 0 |
| Luteal - Akr1c18 | Uchl1 | 83.83878 | 2.7010727 | 0 | 0 |
| Luteal - Akr1c18 | Cemip | 83.634674 | 3.126249 | 0 | 0 |
| Luteal - Akr1c18 | Phf20l1 | 83.60402 | 2.294769 | 0 | 0 |
| Luteal - Akr1c18 | Tsc22d1 | 81.26306 | 1.0359223 | 0 | 0 |
| Luteal - Akr1c18 | Maf | 80.9411 | 2.9232275 | 0 | 0 |
| Luteal - Akr1c18 | Timp1 | 79.30808 | 3.1177168 | 0 | 0 |
| Luteal - Akr1c18 | Cdkn1a | 77.366035 | 3.4775748 | 0 | 0 |
| Luteal - Akr1c18 | Ramp1 | 75.66601 | 3.9200506 | 0 | 0 |
| Luteal - Akr1c18 | Fdx1 | 73.03247 | 1.0758759 | 0 | 0 |
| Luteal - Akr1c18 | Psap | 72.53437 | 1.304739 | 0 | 0 |
| Luteal - Akr1c18 | Epdr1 | 72.340515 | 2.6106613 | 0 | 0 |
| Luteal - Akr1c18 | Pcyt1a | 72.06237 | 2.425841 | 0 | 0 |
| Luteal - Akr1c18 | Sparc | 71.31212 | 1.0864953 | 0 | 0 |
| Luteal - Akr1c18 | Sema3d | 68.42016 | 4.176602 | 0 | 0 |
| Luteal - Akr1c18 | Bzw1 | 68.29028 | 1.5603215 | 0 | 0 |
| Luteal - Akr1c18 | Wdr41 | 66.99327 | 2.7814631 | 0 | 0 |
| Theca | Col1a2 | 84.96591 | 3.3704832 | 0 | 0 |
| Theca | Igfbp7 | 65.566185 | 2.3381317 | 0 | 0 |
| Theca | Col3a1 | 56.633022 | 2.3212903 | 0 | 0 |
| Theca | Prkar2b | 55.16619 | 2.2159045 | 0 | 0 |
| Theca | Col4a1 | 54.40028 | 2.6970541 | 0 | 0 |
| Theca | Cyp17a1 | 53.09798 | 3.7148209 | 0 | 0 |
| Theca | Smoc2 | 47.87451 | 2.5042095 | 0 | 0 |
| Theca | Inhba | 47.604797 | 2.150309 | 0 | 0 |
| Theca | Inha | 44.65387 | 1.6788551 | 0 | 0 |
| Theca | Gas6 | 41.152008 | 1.52581 | 0 | 0 |
| Theca | Ptch1 | 40.621914 | 3.1708682 | 0 | 0 |
| Theca | Tpm1 | 40.113132 | 1.4768498 | 0 | 0 |
| Theca | Trib2 | 39.109787 | 1.886103 | 0 | 0 |
| Theca | Cped1 | 37.32256 | 1.7304207 | 7.07E-305 | 1.38E-301 |
| Theca | Tsc22d1 | 36.6847 | 1.2156522 | 1.28E-294 | 2.33E-291 |
| Theca | Fads2 | 36.5935 | 2.0292683 | 3.63E-293 | 6.19E-290 |
| Theca | Cald1 | 34.868317 | 1.5246679 | 2.25E-266 | 3.61E-263 |
| Theca | Gja1 | 34.474903 | 1.3159355 | 1.91E-260 | 2.89E-257 |
| Theca | Gab2 | 33.827023 | 1.7266265 | 7.90E-251 | 1.14E-247 |
| Theca | Serpine2 | 33.764374 | 1.3292468 | 6.58E-250 | 8.98E-247 |
| Theca | Acta2 | 33.60865 | 1.6895698 | 1.25E-247 | 1.63E-244 |
| Theca | Lama2 | 33.508286 | 2.8840554 | 3.65E-246 | 4.53E-243 |

| | | | | | |
|---|---|---|---|---|---|
| Theca | Col4a2 | 33.094795 | 2.3844187 | 3.53E-240 | 4.19E-237 |
| Theca | Rplp0 | 32.744587 | 1.1359155 | 3.63E-235 | 4.12E-232 |
| Theca | Nedd4 | 32.371975 | 1.1747274 | 6.81E-230 | 7.44E-227 |
| Theca | Enpep | 32.31391 | 5.0810647 | 4.46E-229 | 4.68E-226 |
| Theca | Rbbp7 | 31.694515 | 1.2722708 | 1.85E-220 | 1.87E-217 |
| Theca | Rack1 | 31.090933 | 1.0655743 | 3.19E-212 | 3.11E-209 |
| Theca | Pbx1 | 30.28861 | 1.275908 | 1.62E-201 | 1.52E-198 |
| Theca | Lamb1 | 30.279327 | 1.9856865 | 2.15E-201 | 1.95E-198 |
| Theca | Rpl4 | 30.106056 | 1.0155101 | 4.04E-199 | 3.56E-196 |
| Theca | Serpinh1 | 29.539635 | 1.313179 | 8.92E-192 | 7.61E-189 |
| Theca | Acsbg1 | 29.304407 | 1.0856049 | 9.11E-189 | 7.54E-186 |
| Theca | Mid1ip1 | 29.084976 | 1.6567719 | 5.56E-186 | 4.47E-183 |
| Theca | Fbn1 | 28.91284 | 2.708784 | 8.23E-184 | 6.42E-181 |
| Theca | Aldh1a1 | 28.64299 | 1.1377951 | 1.96E-180 | 1.49E-177 |
| Theca | AU020206 | 28.544529 | 1.0552368 | 3.28E-179 | 2.42E-176 |
| Theca | Nr2f2 | 28.444439 | 1.5944098 | 5.71E-178 | 4.10E-175 |
| Theca | Rpl8 | 27.866888 | 0.9666902 | 6.72E-171 | 4.59E-168 |
| Theca | Rps5 | 27.58661 | 0.9458564 | 1.61E-167 | 1.07E-164 |
| Theca | Eef2 | 27.092937 | 0.9393777 | 1.19E-161 | 7.75E-159 |
| Theca | Eef1b2 | 26.769348 | 0.94707716 | 7.35E-158 | 4.67E-155 |
| Theca | Gstm2 | 26.640345 | 1.1919861 | 2.32E-156 | 1.40E-153 |
| Theca | Npm1 | 26.55238 | 0.97943467 | 2.41E-155 | 1.43E-152 |
| Theca | Hsp90ab1 | 26.113861 | 0.913405 | 2.54E-150 | 1.47E-147 |
| Theca | Rps3 | 25.42919 | 0.9009656 | 1.20E-142 | 6.82E-140 |
| Theca | Septin11 | 25.19264 | 1.6960386 | 4.82E-140 | 2.69E-137 |
| Theca | Hspa5 | 25.178831 | 0.92398286 | 6.83E-140 | 3.73E-137 |
| Theca | Gramd1b | 25.07798 | 1.0850565 | 8.65E-139 | 4.63E-136 |
| Theca | Gpc3 | 24.964779 | 3.4686115 | 1.48E-137 | 7.75E-135 |
| Endothelial - blood | Hbb-bs | 150.60548 | 8.217234 | 0 | 0 |
| Endothelial - blood | Hba-a1 | 93.34336 | 8.589102 | 0 | 0 |
| Endothelial - blood | Hba-a2 | 79.4823 | 8.584898 | 0 | 0 |
| Endothelial - blood | Hbb-bt | 72.096306 | 8.220582 | 0 | 0 |
| Endothelial - blood | mt-Rnr2 | 42.261467 | 0.5753819 | 0 | 0 |
| Endothelial - blood | Aldh1a1 | 32.22558 | 0.88544095 | 7.74E-228 | 3.02E-224 |
| Endothelial - blood | Cyp11a1 | 29.620453 | 0.70100635 | 8.15E-193 | 2.78E-189 |
| Endothelial - blood | Ftl1 | 26.53241 | 0.6542333 | 4.10E-155 | 1.24E-151 |
| Endothelial - blood | Adh1 | 24.8508 | 1.2333741 | 2.53E-136 | 6.29E-133 |
| Endothelial - blood | Tmsb4x | 23.88012 | 0.71696687 | 4.93E-126 | 9.61E-123 |
| Endothelial - blood | Cmss1 | 23.408596 | 0.7255947 | 3.49E-121 | 6.36E-118 |
| Endothelial - blood | Fdx1 | 23.386644 | 0.6231622 | 5.84E-121 | 9.97E-118 |
| Endothelial - blood | mt-Nd1 | 22.74636 | 0.34970087 | 1.56E-114 | 2.50E-111 |
| Endothelial - blood | Rn18s | 22.471241 | 0.3982228 | 7.93E-112 | 1.14E-108 |
| Endothelial - blood | Fth1 | 21.470688 | 0.32345447 | 2.93E-102 | 3.80E-99 |

| | | | | | |
|---|---|---|---|---|---|
| Endothelial - blood | Mt1 | 20.549347 | 0.87170005 | 7.80E-94 | 8.87E-91 |
| Endothelial - blood | Rps14 | 19.814232 | 0.3518313 | 2.24E-87 | 2.36E-84 |
| Endothelial - blood | Dbi | 18.460327 | 0.76762486 | 4.31E-76 | 3.79E-73 |
| Endothelial - blood | Rpl32 | 17.350317 | 0.5330678 | 1.96E-67 | 1.49E-64 |
| Endothelial - blood | Rps24 | 16.75013 | 0.5737398 | 5.65E-63 | 3.76E-60 |
| Endothelial - blood | Hsd3b1 | 16.456501 | 0.33132184 | 7.53E-61 | 4.78E-58 |
| Endothelial - blood | Mgarp | 16.29601 | 0.50692004 | 1.05E-59 | 6.25E-57 |
| Endothelial - blood | Rps19 | 15.72229 | 0.6036641 | 1.06E-55 | 5.70E-53 |
| Endothelial - blood | Rps20 | 15.492661 | 0.4705885 | 3.89E-54 | 2.00E-51 |
| Endothelial - blood | Gstm1 | 14.897462 | 0.5994664 | 3.42E-50 | 1.46E-47 |
| Endothelial - blood | Dnajc15 | 14.245463 | 0.6242697 | 4.78E-46 | 1.81E-43 |
| Endothelial - blood | Hspe1 | 14.155477 | 0.7628872 | 1.73E-45 | 6.37E-43 |
| Endothelial - blood | Rpl41 | 13.91766 | 0.80208594 | 4.95E-44 | 1.73E-41 |
| Endothelial - blood | Rps8 | 13.774485 | 0.59064716 | 3.63E-43 | 1.22E-40 |
| Endothelial - blood | Rplp1 | 13.008574 | 0.51637894 | 1.09E-38 | 3.28E-36 |
| Endothelial - blood | Ubb | 12.883867 | 0.4548912 | 5.55E-38 | 1.51E-35 |
| Endothelial - blood | Rpl18 | 12.615705 | 0.90086824 | 1.73E-36 | 4.41E-34 |
| Endothelial - blood | Rps21 | 12.393809 | 0.47071472 | 2.82E-35 | 6.76E-33 |
| Endothelial - blood | Rpl13 | 11.638933 | 0.58652943 | 2.61E-31 | 5.32E-29 |
| Endothelial - blood | Cox6c | 11.557501 | 0.55269325 | 6.76E-31 | 1.35E-28 |
| Endothelial - blood | Tmsb10 | 11.552717 | 0.7604103 | 7.15E-31 | 1.42E-28 |
| Endothelial - blood | Me1 | 11.223091 | 0.370855 | 3.14E-29 | 5.76E-27 |
| Endothelial - blood | Rpl23 | 10.982289 | 0.5394156 | 4.65E-28 | 8.04E-26 |
| Endothelial - blood | Atp5j2 | 10.970913 | 0.46368414 | 5.27E-28 | 9.06E-26 |
| Endothelial - blood | Myl6 | 10.952134 | 0.53718394 | 6.49E-28 | 1.11E-25 |
| Endothelial - blood | Rps12 | 10.836996 | 0.9035631 | 2.30E-27 | 3.83E-25 |
| Endothelial - blood | Gstm2 | 10.804656 | 0.57979333 | 3.27E-27 | 5.41E-25 |
| Endothelial - blood | Rps18 | 10.653135 | 0.56172085 | 1.69E-26 | 2.69E-24 |
| Endothelial - blood | Rpl34 | 10.635913 | 0.56581396 | 2.03E-26 | 3.16E-24 |
| Endothelial - blood | Chchd2 | 10.206519 | 0.38614494 | 1.85E-24 | 2.60E-22 |
| Endothelial - blood | Mgst1 | 10.195063 | 0.4487711 | 2.09E-24 | 2.91E-22 |
| Endothelial - blood | Tpt1 | 10.183278 | 1.0283798 | 2.35E-24 | 3.25E-22 |
| Endothelial - blood | Atp5e | 10.050038 | 0.60630786 | 9.18E-24 | 1.21E-21 |
| Endothelial - blood | Rpl11 | 10.020704 | 0.9358377 | 1.24E-23 | 1.60E-21 |
| Endothelial - blood | Mt2 | 9.875886 | 0.75803447 | 5.30E-23 | 6.60E-21 |
| Endothelial - lymphatic | Ccl21a | 119.79292 | 6.9681244 | 0 | 0 |
| Endothelial - lymphatic | Mmrn1 | 63.529144 | 7.0366254 | 0 | 0 |
| Endothelial - lymphatic | Selenop | 49.675453 | 2.0428574 | 0 | 0 |
| Endothelial - lymphatic | Tmsb4x | 47.89389 | 1.3607448 | 0 | 0 |
| Endothelial - lymphatic | Igfbp4 | 38.90515 | 1.8207476 | 0 | 0 |
| Endothelial - lymphatic | Cavin2 | 37.112785 | 3.558803 | 1.75E-301 | 4.77E-298 |
| Endothelial - lymphatic | Lyve1 | 37.04634 | 6.1935735 | 2.06E-300 | 5.11E-297 |
| Endothelial - lymphatic | Aqp1 | 36.2988 | 5.599985 | 1.69E-288 | 3.85E-285 |

| | | | | | |
|---|---|---|---|---|---|
| Endothelial - lymphatic | Gng11 | 35.16705 | 2.8782961 | 6.38E-271 | 1.24E-267 |
| Endothelial - lymphatic | Tagln | 34.151604 | 1.8006167 | 1.27E-255 | 2.30E-252 |
| Endothelial - lymphatic | Fth1 | 33.34882 | 0.6008673 | 7.58E-244 | 1.15E-240 |
| Endothelial - lymphatic | Dcn | 33.063793 | 1.6711383 | 9.86E-240 | 1.42E-236 |
| Endothelial - lymphatic | Fgl2 | 32.322094 | 3.202932 | 3.42E-229 | 4.25E-226 |
| Endothelial - lymphatic | Acta2 | 31.602108 | 1.5785072 | 3.45E-219 | 3.93E-216 |
| Endothelial - lymphatic | Sparcl1 | 29.623402 | 1.3873335 | 7.47E-193 | 7.55E-190 |
| Endothelial - lymphatic | Stab1 | 28.67978 | 3.3226233 | 6.82E-181 | 6.21E-178 |
| Endothelial - lymphatic | Rarres2 | 28.601442 | 1.453231 | 6.45E-180 | 5.33E-177 |
| Endothelial - lymphatic | Mgp | 27.99853 | 1.1710901 | 1.69E-172 | 1.28E-169 |
| Endothelial - lymphatic | Cfh | 26.918869 | 1.143784 | 1.32E-159 | 9.25E-157 |
| Endothelial - lymphatic | Ifitm3 | 26.073946 | 1.5033188 | 7.20E-150 | 4.68E-147 |
| Endothelial - lymphatic | Myh11 | 24.357807 | 1.5659077 | 4.79E-131 | 2.78E-128 |
| Endothelial - lymphatic | Egfl7 | 23.774736 | 2.3313031 | 6.10E-125 | 3.47E-122 |
| Endothelial - lymphatic | Tpm2 | 23.277702 | 1.3336031 | 7.46E-120 | 4.07E-117 |
| Endothelial - lymphatic | mt-Rnr2 | 23.275742 | 0.2815563 | 7.81E-120 | 4.18E-117 |
| Endothelial - lymphatic | Prox1 | 22.85708 | 5.422326 | 1.24E-115 | 6.52E-113 |
| Endothelial - lymphatic | H2-D1 | 21.91171 | 1.1895918 | 2.01E-106 | 9.30E-104 |
| Endothelial - lymphatic | Fxyd6 | 20.859362 | 2.142819 | 1.25E-96 | 5.43E-94 |
| Endothelial - lymphatic | Ahnak | 19.91705 | 1.0438594 | 2.90E-88 | 1.16E-85 |
| Endothelial - lymphatic | Myl9 | 19.819805 | 1.2886448 | 2.01E-87 | 7.95E-85 |
| Endothelial - lymphatic | Igfbp7 | 19.504831 | 0.66481376 | 9.99E-85 | 3.79E-82 |
| Endothelial - lymphatic | Reln | 19.215664 | 4.5338426 | 2.74E-82 | 9.71E-80 |
| Endothelial - lymphatic | B2m | 19.154444 | 0.9348467 | 8.89E-82 | 3.11E-79 |
| Endothelial - lymphatic | Rhoj | 18.611048 | 1.9657022 | 2.61E-77 | 8.81E-75 |
| Endothelial - lymphatic | Gnas | 18.24846 | 0.43643296 | 2.13E-74 | 6.92E-72 |
| Endothelial - lymphatic | Timp3 | 17.962013 | 1.0201641 | 3.87E-72 | 1.20E-69 |
| Endothelial - lymphatic | Mylk | 17.911312 | 1.1299436 | 9.62E-72 | 2.92E-69 |
| Endothelial - lymphatic | Timp2 | 17.863316 | 1.1843324 | 2.28E-71 | 6.83E-69 |
| Endothelial - lymphatic | Tgm2 | 17.771664 | 1.8742006 | 1.17E-70 | 3.48E-68 |
| Endothelial - lymphatic | Flt4 | 17.656101 | 4.9825654 | 9.13E-70 | 2.68E-67 |
| Endothelial - lymphatic | Kdr | 17.52714 | 2.7969813 | 8.89E-69 | 2.58E-66 |
| Endothelial - lymphatic | Nrp2 | 17.398523 | 1.7573571 | 8.47E-68 | 2.43E-65 |
| Endothelial - lymphatic | Cald1 | 17.051672 | 0.8026649 | 3.40E-65 | 9.56E-63 |
| Endothelial - lymphatic | Ltbp4 | 16.682175 | 2.3110101 | 1.77E-62 | 4.73E-60 |
| Endothelial - lymphatic | Prss23 | 16.480545 | 1.386934 | 5.06E-61 | 1.30E-58 |
| Endothelial - lymphatic | H2-K1 | 16.23999 | 0.74830663 | 2.63E-59 | 6.47E-57 |
| Endothelial - lymphatic | Nfib | 16.126469 | 1.3722293 | 1.66E-58 | 4.02E-56 |
| Endothelial - lymphatic | Tgfbr2 | 15.841678 | 2.3139553 | 1.60E-56 | 3.71E-54 |
| Endothelial - lymphatic | Apoe | 15.676114 | 0.54480404 | 2.20E-55 | 4.97E-53 |
| Endothelial - lymphatic | Elk3 | 15.416895 | 2.3883026 | 1.26E-53 | 2.65E-51 |
| Endothelial - lymphatic | Ogn | 15.174722 | 1.5455668 | 5.20E-52 | 1.05E-49 |
| Endothelial - vascular | Igfbp7 | 109.41886 | 2.433846 | 0 | 0 |

| | | | | | |
|---|---|---|---|---|---|
| Endothelial - vascular | Sparc | 94.49793 | 1.9327084 | 0 | 0 |
| Endothelial - vascular | Col4a1 | 78.7142 | 2.6641078 | 0 | 0 |
| Endothelial - vascular | Tmsb4x | 62.492363 | 1.3553244 | 0 | 0 |
| Endothelial - vascular | Vim | 61.802635 | 1.6468667 | 0 | 0 |
| Endothelial - vascular | Egfl7 | 54.5514 | 3.179414 | 0 | 0 |
| Endothelial - vascular | Col4a2 | 53.943424 | 2.6189983 | 0 | 0 |
| Endothelial - vascular | Rbp1 | 51.490314 | 1.8179331 | 0 | 0 |
| Endothelial - vascular | Mgp | 49.6096 | 1.4432538 | 0 | 0 |
| Endothelial - vascular | Actb | 48.823437 | 0.8487363 | 0 | 0 |
| Endothelial - vascular | Flt1 | 43.409607 | 4.212952 | 0 | 0 |
| Endothelial - vascular | Epas1 | 40.55967 | 2.23639 | 0 | 0 |
| Endothelial - vascular | Sptbn1 | 40.270134 | 1.145241 | 0 | 0 |
| Endothelial - vascular | Myh9 | 39.765682 | 1.4128939 | 0 | 0 |
| Endothelial - vascular | Ehd4 | 39.702526 | 2.3682206 | 0 | 0 |
| Endothelial - vascular | B2m | 39.555832 | 1.2063878 | 0 | 0 |
| Endothelial - vascular | Tcf4 | 38.47479 | 0.99468064 | 0 | 0 |
| Endothelial - vascular | Ptprb | 38.27807 | 3.2092009 | 0 | 0 |
| Endothelial - vascular | Cald1 | 38.098503 | 1.1359094 | 0 | 0 |
| Endothelial - vascular | Ifitm3 | 38.069874 | 1.4657496 | 0 | 0 |
| Endothelial - vascular | H2-K1 | 37.75637 | 1.0687488 | 0 | 0 |
| Endothelial - vascular | Tpm4 | 36.901886 | 1.4084103 | 4.31E-298 | 5.12E-295 |
| Endothelial - vascular | Col3a1 | 36.71838 | 1.0872482 | 3.72E-295 | 4.23E-292 |
| Endothelial - vascular | Col1a2 | 35.391186 | 1.0397793 | 2.33E-274 | 2.55E-271 |
| Endothelial - vascular | Sparcl1 | 35.32959 | 1.1542175 | 2.06E-273 | 2.17E-270 |
| Endothelial - vascular | Cd93 | 34.299873 | 3.390948 | 7.88E-258 | 7.97E-255 |
| Endothelial - vascular | Igfbp4 | 34.021793 | 1.1710949 | 1.06E-253 | 1.03E-250 |
| Endothelial - vascular | Cd34 | 32.181355 | 2.6646957 | 3.22E-227 | 2.83E-224 |
| Endothelial - vascular | Nfib | 31.64344 | 1.6573145 | 9.33E-220 | 7.96E-217 |
| Endothelial - vascular | Tagln2 | 31.238598 | 1.6920002 | 3.19E-214 | 2.64E-211 |
| Endothelial - vascular | Pcp4l1 | 31.005713 | 2.1790693 | 4.51E-211 | 3.63E-208 |
| Endothelial - vascular | Kdr | 30.703106 | 3.1651864 | 5.17E-207 | 4.04E-204 |
| Endothelial - vascular | H2-D1 | 30.493183 | 1.0874588 | 3.21E-204 | 2.37E-201 |
| Endothelial - vascular | Piezo2 | 29.69311 | 2.272397 | 9.42E-194 | 6.77E-191 |
| Endothelial - vascular | Tm4sf1 | 28.738482 | 2.8884196 | 1.26E-181 | 8.83E-179 |
| Endothelial - vascular | Calm1 | 28.62224 | 0.64753294 | 3.55E-180 | 2.43E-177 |
| Endothelial - vascular | Anxa2 | 28.382359 | 0.82807213 | 3.34E-177 | 2.22E-174 |
| Endothelial - vascular | Tpm1 | 28.11989 | 0.67980605 | 5.60E-174 | 3.64E-171 |
| Endothelial - vascular | Cdh5 | 27.698963 | 3.417246 | 7.19E-169 | 4.56E-166 |
| Endothelial - vascular | Pecam1 | 27.372177 | 3.3475425 | 5.88E-165 | 3.65E-162 |
| Endothelial - vascular | Tuba1a | 27.362339 | 1.0230755 | 7.70E-165 | 4.67E-162 |
| Endothelial - vascular | Ctnnb1 | 27.326164 | 0.66765976 | 2.07E-164 | 1.23E-161 |
| Endothelial - vascular | Ctla2a | 27.254368 | 2.648689 | 1.48E-163 | 8.57E-161 |
| Endothelial - vascular | Gnai2 | 27.101084 | 1.1008061 | 9.56E-162 | 5.44E-159 |

| | | | | | |
|---|---|---|---|---|---|
| Endothelial - vascular | Slc9a3r2 | 26.995691 | 2.3210626 | 1.66E-160 | 9.25E-158 |
| Endothelial - vascular | Ets1 | 26.88938 | 2.798763 | 2.92E-159 | 1.60E-156 |
| Endothelial - vascular | Ly6e | 26.767656 | 1.8050773 | 7.69E-158 | 4.12E-155 |
| Endothelial - vascular | Smarca2 | 26.725718 | 0.8600918 | 2.37E-157 | 1.24E-154 |
| Endothelial - vascular | Septin7 | 26.640282 | 0.8730803 | 2.32E-156 | 1.19E-153 |
| Endothelial - vascular | Rgs5 | 26.464949 | 2.8178124 | 2.46E-154 | 1.24E-151 |
| Epithelial - surface | Lgals7 | 52.706436 | 6.5851555 | 0 | 0 |
| Epithelial - surface | Kctd14 | 51.21717 | 3.0549095 | 0 | 0 |
| Epithelial - surface | Tmsb4x | 45.111492 | 1.3814371 | 0 | 0 |
| Epithelial - surface | mt-Rnr2 | 42.400455 | 0.59853137 | 0 | 0 |
| Epithelial - surface | Krt19 | 35.02481 | 6.0052333 | 9.43E-269 | 1.43E-265 |
| Epithelial - surface | Ly6e | 32.27191 | 2.8984063 | 1.73E-228 | 2.25E-225 |
| Epithelial - surface | Epcam | 31.328123 | 5.9650683 | 1.93E-215 | 2.11E-212 |
| Epithelial - surface | Igfbp5 | 30.945688 | 2.8527467 | 2.90E-210 | 2.94E-207 |
| Epithelial - surface | Ildr2 | 29.794872 | 4.250794 | 4.55E-195 | 3.88E-192 |
| Epithelial - surface | Krt18 | 28.572716 | 4.9776587 | 1.47E-179 | 1.14E-176 |
| Epithelial - surface | Krt7 | 28.273022 | 5.964536 | 7.42E-176 | 5.33E-173 |
| Epithelial - surface | Mt2 | 26.579386 | 1.8133031 | 1.18E-155 | 7.46E-153 |
| Epithelial - surface | Plxna4 | 26.281591 | 4.148121 | 3.11E-152 | 1.93E-149 |
| Epithelial - surface | Rarres2 | 24.414574 | 1.4338856 | 1.20E-131 | 5.74E-129 |
| Epithelial - surface | Wt1 | 24.011364 | 2.6001782 | 2.12E-127 | 9.17E-125 |
| Epithelial - surface | Rbbp7 | 23.812777 | 1.0999002 | 2.46E-125 | 1.03E-122 |
| Epithelial - surface | mt-Rnr1 | 21.860752 | 0.46683505 | 6.14E-106 | 2.15E-103 |
| Epithelial - surface | Upk1b | 21.451208 | 6.4419804 | 4.45E-102 | 1.48E-99 |
| Epithelial - surface | Atp1b1 | 21.159933 | 2.8541238 | 2.24E-99 | 7.18E-97 |
| Epithelial - surface | Crip1 | 20.878742 | 2.628034 | 8.36E-97 | 2.59E-94 |
| Epithelial - surface | Lmo7 | 20.714985 | 3.1404629 | 2.54E-95 | 7.79E-93 |
| Epithelial - surface | Upk3b | 20.145325 | 6.2030067 | 2.96E-90 | 8.24E-88 |
| Epithelial - surface | Mt1 | 19.45225 | 0.90167534 | 2.79E-84 | 7.12E-82 |
| Epithelial - surface | F11r | 19.43611 | 1.8175446 | 3.82E-84 | 9.66E-82 |
| Epithelial - surface | Lsr | 19.306852 | 2.2954462 | 4.70E-83 | 1.16E-80 |
| Epithelial - surface | Lhx9 | 19.08338 | 5.8610606 | 3.47E-81 | 8.39E-79 |
| Epithelial - surface | Unc45b | 18.95559 | 4.7479954 | 3.97E-80 | 9.35E-78 |
| Epithelial - surface | Lgals2 | 17.428152 | 5.8341365 | 5.04E-68 | 9.98E-66 |
| Epithelial - surface | Lgr5 | 17.074854 | 3.6475978 | 2.28E-65 | 4.24E-63 |
| Epithelial - surface | Ppfibp2 | 16.826586 | 3.1314957 | 1.56E-63 | 2.84E-61 |
| Epithelial - surface | Magi1 | 16.755602 | 1.8574182 | 5.15E-63 | 9.14E-61 |
| Epithelial - surface | Txnip | 16.677023 | 0.88482016 | 1.93E-62 | 3.39E-60 |
| Epithelial - surface | Igfbp6 | 15.244013 | 1.6303867 | 1.80E-52 | 2.63E-50 |
| Epithelial - surface | Sdc4 | 14.642908 | 1.368132 | 1.50E-48 | 1.96E-46 |
| Epithelial - surface | Adamts19 | 14.617218 | 2.3063457 | 2.18E-48 | 2.82E-46 |
| Epithelial - surface | Cald1 | 14.61657 | 0.81135404 | 2.20E-48 | 2.84E-46 |
| Epithelial - surface | Syne2 | 14.610558 | 0.6151786 | 2.41E-48 | 3.08E-46 |

| | | | | | |
|---|---|---|---|---|---|
| Epithelial - surface | Hsd17b2 | 14.348537 | 5.879675 | 1.09E-46 | 1.33E-44 |
| Epithelial - surface | Aldh1a2 | 14.258567 | 2.1160944 | 3.97E-46 | 4.81E-44 |
| Epithelial - surface | Creg1 | 13.74262 | 1.2827067 | 5.64E-43 | 6.23E-41 |
| Epithelial - surface | Gng13 | 13.543669 | 5.3381214 | 8.64E-42 | 9.28E-40 |
| Epithelial - surface | Nrg1 | 13.509174 | 4.435493 | 1.38E-41 | 1.48E-39 |
| Epithelial - surface | Emx2 | 13.479434 | 2.8995047 | 2.07E-41 | 2.20E-39 |
| Epithelial - surface | Pcnx2 | 13.4195 | 5.3424325 | 4.65E-41 | 4.92E-39 |
| Epithelial - surface | Isyna1 | 13.069394 | 1.1548326 | 4.93E-39 | 4.89E-37 |
| Epithelial - surface | Ablim1 | 12.709218 | 1.1150554 | 5.26E-37 | 4.83E-35 |
| Epithelial - surface | Cgn | 12.68993 | 3.5660107 | 6.72E-37 | 6.16E-35 |
| Epithelial - surface | Cryl1 | 12.316505 | 1.7599102 | 7.38E-35 | 6.28E-33 |
| Epithelial - surface | Zfp36l1 | 12.252739 | 1.0607172 | 1.62E-34 | 1.37E-32 |
| Epithelial - surface | Rnf213 | 11.9297285 | 1.5006133 | 8.28E-33 | 6.69E-31 |
| Epithelial - oviduct | Ovgp1 | 80.816986 | 9.5285225 | 0 | 0 |
| Epithelial - oviduct | Gsto1 | 38.618744 | 4.4248424 | 0 | 0 |
| Epithelial - oviduct | Alcam | 28.995975 | 5.209503 | 7.39E-185 | 3.37E-181 |
| Epithelial - oviduct | Plet1 | 27.142353 | 7.2425857 | 3.12E-162 | 1.06E-158 |
| Epithelial - oviduct | S100g | 24.348099 | 5.9284763 | 6.07E-131 | 1.66E-127 |
| Epithelial - oviduct | C3 | 24.106401 | 4.0857882 | 2.14E-128 | 5.32E-125 |
| Epithelial - oviduct | Cbr2 | 22.599115 | 6.8594894 | 4.42E-113 | 9.29E-110 |
| Epithelial - oviduct | Rnf128 | 19.67726 | 3.911194 | 3.38E-86 | 5.76E-83 |
| Epithelial - oviduct | Tmsb4x | 19.617899 | 1.0217398 | 1.09E-85 | 1.75E-82 |
| Epithelial - oviduct | Lcn2 | 18.481575 | 7.1142087 | 2.91E-76 | 3.45E-73 |
| Epithelial - oviduct | mt-Rnr2 | 18.138767 | 0.43883926 | 1.58E-73 | 1.65E-70 |
| Epithelial - oviduct | Krt19 | 17.908487 | 4.9019895 | 1.01E-71 | 9.87E-69 |
| Epithelial - oviduct | Krt18 | 17.710104 | 4.6027336 | 3.50E-70 | 3.19E-67 |
| Epithelial - oviduct | Emb | 17.06158 | 3.7803714 | 2.87E-65 | 2.12E-62 |
| Epithelial - oviduct | Ltf | 15.867255 | 6.102934 | 1.07E-56 | 7.29E-54 |
| Epithelial - oviduct | Dcxr | 15.187909 | 4.088226 | 4.25E-52 | 2.58E-49 |
| Epithelial - oviduct | Ehf | 14.6707 | 4.5892887 | 9.93E-49 | 5.53E-46 |
| Epithelial - oviduct | Padi1 | 13.739396 | 7.5749636 | 5.90E-43 | 2.93E-40 |
| Epithelial - oviduct | Fxyd4 | 13.4250345 | 7.468564 | 4.31E-41 | 2.03E-38 |
| Epithelial - oviduct | Ncald | 12.118143 | 2.6428416 | 8.47E-34 | 3.12E-31 |
| Epithelial - oviduct | Slc14a1 | 11.979647 | 6.957094 | 4.54E-33 | 1.65E-30 |
| Epithelial - oviduct | Chdh | 11.854272 | 4.756749 | 2.04E-32 | 7.35E-30 |
| Epithelial - oviduct | Cyb5r3 | 11.186956 | 1.3200219 | 4.72E-29 | 1.45E-26 |
| Epithelial - oviduct | Gstm2 | 11.1704445 | 1.0066792 | 5.69E-29 | 1.73E-26 |
| Epithelial - oviduct | Chpt1 | 10.787552 | 1.9270846 | 3.94E-27 | 1.11E-24 |
| Epithelial - oviduct | Atp1b1 | 10.439107 | 2.426037 | 1.64E-25 | 4.36E-23 |
| Epithelial - oviduct | Spint2 | 10.216194 | 1.9693184 | 1.68E-24 | 4.24E-22 |
| Epithelial - oviduct | Krt7 | 10.039156 | 4.1649313 | 1.03E-23 | 2.55E-21 |
| Epithelial - oviduct | Wwc1 | 9.898967 | 4.3996215 | 4.21E-23 | 1.03E-20 |
| Epithelial - oviduct | Cd24a | 9.610157 | 4.680915 | 7.24E-22 | 1.66E-19 |

| | | | | | |
|---|---|---|---|---|---|
| Epithelial - oviduct | Epcam | 9.543332 | 3.8667943 | 1.38E-21 | 3.05E-19 |
| Epithelial - oviduct | mt-Rnr1 | 9.521297 | 0.2878097 | 1.71E-21 | 3.71E-19 |
| Epithelial - oviduct | Cbs | 9.395905 | 3.9697826 | 5.67E-21 | 1.15E-18 |
| Epithelial - oviduct | Ifitm3 | 9.133282 | 1.0680207 | 6.65E-20 | 1.24E-17 |
| Epithelial - oviduct | Hpgds | 9.095736 | 3.8237793 | 9.39E-20 | 1.73E-17 |
| Epithelial - oviduct | Mecom | 9.092205 | 3.9148204 | 9.71E-20 | 1.78E-17 |
| Epithelial - oviduct | Ezr | 9.042979 | 1.8928052 | 1.52E-19 | 2.74E-17 |
| Epithelial - oviduct | Slc25a48 | 9.034571 | 4.6456957 | 1.65E-19 | 2.92E-17 |
| Epithelial - oviduct | Bcat1 | 8.987521 | 1.2045314 | 2.53E-19 | 4.43E-17 |
| Epithelial - oviduct | Upk1a | 8.949367 | 7.39447 | 3.58E-19 | 6.18E-17 |
| Epithelial - oviduct | Ivns1abp | 8.901699 | 0.8597595 | 5.50E-19 | 9.39E-17 |
| Epithelial - oviduct | Gpx1 | 8.791915 | 0.79285496 | 1.47E-18 | 2.46E-16 |
| Epithelial - oviduct | Cd9 | 8.728523 | 1.6703556 | 2.58E-18 | 4.22E-16 |
| Epithelial - oviduct | Slc39a4 | 8.340885 | 6.036488 | 7.37E-17 | 1.13E-14 |
| Epithelial - oviduct | Mmp7 | 8.267454 | 8.7219095 | 1.37E-16 | 2.06E-14 |
| Epithelial - oviduct | Wfdc2 | 8.249608 | 7.2059608 | 1.59E-16 | 2.35E-14 |
| Epithelial - oviduct | Ctsb | 8.19778 | 0.6452234 | 2.45E-16 | 3.52E-14 |
| Epithelial - oviduct | Cdkl1 | 8.147486 | 5.5365195 | 3.72E-16 | 5.28E-14 |
| Epithelial - oviduct | Ppp2r2b | 7.647705 | 3.9426258 | 2.05E-14 | 2.56E-12 |
| Epithelial - oviduct | Esr1 | 7.641411 | 1.6584959 | 2.15E-14 | 2.68E-12 |
| Immune | Apoe | 228.93802 | 4.5163417 | 0 | 0 |
| Immune | Fth1 | 175.79587 | 2.235694 | 0 | 0 |
| Immune | Lyz2 | 172.19327 | 4.7626305 | 0 | 0 |
| Immune | Ctss | 154.85817 | 4.7176247 | 0 | 0 |
| Immune | Ctsd | 144.88217 | 3.3097193 | 0 | 0 |
| Immune | Ftl1 | 142.32292 | 2.1349585 | 0 | 0 |
| Immune | Gpnmb | 133.57133 | 5.0670953 | 0 | 0 |
| Immune | Psap | 132.68803 | 2.6612105 | 0 | 0 |
| Immune | Ctsb | 130.55074 | 2.66829 | 0 | 0 |
| Immune | Lgals3 | 125.6545 | 3.7558107 | 0 | 0 |
| Immune | Tmsb4x | 108.198456 | 1.8633521 | 0 | 0 |
| Immune | H2-D1 | 101.56525 | 2.728712 | 0 | 0 |
| Immune | Selenop | 95.40794 | 2.363476 | 0 | 0 |
| Immune | H2-Ab1 | 89.24523 | 3.502211 | 0 | 0 |
| Immune | Cd63 | 84.1151 | 1.4817092 | 0 | 0 |
| Immune | Actb | 74.69865 | 0.9359435 | 0 | 0 |
| Immune | C1qb | 72.58774 | 3.6204407 | 0 | 0 |
| Immune | H2-Eb1 | 72.299126 | 3.5047238 | 0 | 0 |
| Immune | C1qc | 70.73727 | 3.609047 | 0 | 0 |
| Immune | Mpeg1 | 70.20938 | 4.2205033 | 0 | 0 |
| Immune | Itm2b | 69.57464 | 1.189496 | 0 | 0 |
| Immune | B2m | 68.48095 | 1.7429675 | 0 | 0 |
| Immune | Cd74 | 67.115425 | 3.5870466 | 0 | 0 |

| | | | | | |
|---|---|---|---|---|---|
| Immune | Tyrobp | 63.566418 | 3.6637802 | 0 | 0 |
| Immune | Cyba | 62.558125 | 2.766433 | 0 | 0 |
| Immune | Cst3 | 61.825962 | 1.2511761 | 0 | 0 |
| Immune | H2-Aa | 59.995636 | 3.5291102 | 0 | 0 |
| Immune | Ctsz | 57.770454 | 1.9639715 | 0 | 0 |
| Immune | Creg1 | 54.02674 | 2.1537428 | 0 | 0 |
| Immune | Trf | 53.818493 | 2.4062865 | 0 | 0 |
| Immune | Atp6v0d2 | 52.666916 | 4.5742984 | 0 | 0 |
| Immune | H2-K1 | 52.661 | 1.2799135 | 0 | 0 |
| Immune | C1qa | 49.28391 | 3.401137 | 0 | 0 |
| Immune | Dcn | 49.131027 | 1.504947 | 0 | 0 |
| Immune | Mmp12 | 48.61711 | 4.6193814 | 0 | 0 |
| Immune | Laptm5 | 47.16281 | 3.889396 | 0 | 0 |
| Immune | Cotl1 | 46.158047 | 2.4832492 | 0 | 0 |
| Immune | Igfbp7 | 45.682224 | 0.9200975 | 0 | 0 |
| Immune | Ccl6 | 42.97188 | 3.7543216 | 0 | 0 |
| Immune | Lipa | 42.90018 | 2.5127456 | 0 | 0 |
| Immune | Mgp | 42.25851 | 1.059151 | 0 | 0 |
| Immune | Grn | 41.855076 | 1.6566406 | 0 | 0 |
| Immune | Pltp | 41.484936 | 3.1014335 | 0 | 0 |
| Immune | Igfbp4 | 40.09157 | 1.21504 | 0 | 0 |
| Immune | Fabp4 | 40.006557 | 2.7647567 | 0 | 0 |
| Immune | Sparcl1 | 39.61081 | 1.1278509 | 0 | 0 |
| Immune | Lcp1 | 39.407677 | 3.1428242 | 0 | 0 |
| Immune | Fcer1g | 38.33984 | 3.4276211 | 0 | 0 |
| Immune | Trem2 | 38.08352 | 4.060093 | 0 | 0 |
| Immune | Cd68 | 37.199223 | 4.0686393 | 7.02E-303 | 3.04E-300 |
| Muscle | Acta2 | 149.91876 | 3.7353427 | 0 | 0 |
| Muscle | Tagln | 135.97072 | 3.732134 | 0 | 0 |
| Muscle | Myh11 | 134.9389 | 4.1365156 | 0 | 0 |
| Muscle | Tpm2 | 119.54884 | 3.3724048 | 0 | 0 |
| Muscle | Mylk | 111.12312 | 3.209537 | 0 | 0 |
| Muscle | Col1a2 | 109.88088 | 2.4439726 | 0 | 0 |
| Muscle | Tpm1 | 103.85892 | 1.9135811 | 0 | 0 |
| Muscle | Myl9 | 103.27801 | 3.1990287 | 0 | 0 |
| Muscle | Cald1 | 88.275055 | 2.0601866 | 0 | 0 |
| Muscle | Sparcl1 | 83.19668 | 2.0960193 | 0 | 0 |
| Muscle | Actg2 | 82.01994 | 3.9240544 | 0 | 0 |
| Muscle | Tmsb4x | 79.099396 | 1.3405995 | 0 | 0 |
| Muscle | Cfh | 77.21339 | 1.7667751 | 0 | 0 |
| Muscle | Rarres2 | 76.80736 | 2.0967293 | 0 | 0 |
| Muscle | Dmd | 75.03622 | 2.9014335 | 0 | 0 |
| Muscle | Col3a1 | 69.71865 | 1.6559489 | 0 | 0 |

| | | | | | |
|---|---|---|---|---|---|
| Muscle | Dcn | 64.99552 | 1.9073125 | 0 | 0 |
| Muscle | Mgp | 63.958824 | 1.5148184 | 0 | 0 |
| Muscle | Flna | 63.577778 | 1.9728523 | 0 | 0 |
| Muscle | Igfbp7 | 63.09691 | 1.1844332 | 0 | 0 |
| Muscle | Ogn | 58.229866 | 2.8580468 | 0 | 0 |
| Muscle | Csmd1 | 56.539932 | 1.9965938 | 0 | 0 |
| Muscle | Cped1 | 50.350456 | 1.3324242 | 0 | 0 |
| Muscle | Lpp | 46.79441 | 1.2907001 | 0 | 0 |
| Muscle | Myl6 | 46.687496 | 1.1210136 | 0 | 0 |
| Muscle | Sparc | 45.090267 | 0.7578368 | 0 | 0 |
| Muscle | Csrp1 | 44.131863 | 2.6555464 | 0 | 0 |
| Muscle | Cnn1 | 43.398525 | 4.0920696 | 0 | 0 |
| Muscle | Serpinf1 | 42.3823 | 2.3163424 | 0 | 0 |
| Muscle | Rbpms | 39.75588 | 1.7484477 | 0 | 0 |
| Muscle | Actb | 38.825245 | 0.40697405 | 0 | 0 |
| Muscle | Fstl1 | 38.086765 | 1.9100637 | 0 | 0 |
| Muscle | Rbp1 | 37.230663 | 1.2248917 | 2.18E-303 | 9.75E-301 |
| Muscle | Igfbp6 | 36.77507 | 1.9291314 | 4.62E-296 | 1.94E-293 |
| Muscle | Ndrg2 | 36.073887 | 1.1962545 | 5.82E-285 | 2.27E-282 |
| Muscle | Prkg1 | 35.963745 | 2.2810867 | 3.09E-283 | 1.19E-280 |
| Muscle | Igfbp4 | 35.090496 | 1.0779469 | 9.41E-270 | 3.57E-267 |
| Muscle | Gsn | 34.761555 | 1.8878206 | 9.27E-265 | 3.47E-262 |
| Muscle | Sfrp1 | 34.56379 | 1.9169804 | 8.85E-262 | 3.22E-259 |
| Muscle | Col1a1 | 34.43318 | 2.056205 | 8.04E-260 | 2.89E-257 |
| Muscle | Lgals1 | 34.396557 | 0.68441635 | 2.84E-259 | 1.01E-256 |
| Muscle | Itih5 | 34.177402 | 1.667383 | 5.24E-256 | 1.81E-253 |
| Muscle | Ckb | 33.7798 | 1.7027016 | 3.90E-250 | 1.32E-247 |
| Muscle | Vim | 33.620632 | 0.82506526 | 8.38E-248 | 2.79E-245 |
| Muscle | Des | 33.590904 | 4.0111084 | 2.28E-247 | 7.49E-245 |
| Muscle | Ifitm3 | 33.26204 | 1.1871313 | 1.37E-242 | 4.34E-240 |
| Muscle | Irag1 | 33.158894 | 2.597327 | 4.22E-241 | 1.32E-238 |
| Muscle | Ahnak | 33.101143 | 0.99662524 | 2.86E-240 | 8.88E-238 |
| Muscle | Ppp1r12a | 32.975563 | 1.07511 | 1.82E-238 | 5.58E-236 |
| Muscle | Lmod1 | 32.96621 | 3.4528797 | 2.48E-238 | 7.52E-236 |
| Fibroblast | Malat1 | 77.79906 | 0.99548477 | 0 | 0 |
| Fibroblast | Rad51b | 76.96225 | 3.4395661 | 0 | 0 |
| Fibroblast | Cped1 | 56.159424 | 1.992125 | 0 | 0 |
| Fibroblast | Col1a2 | 46.88947 | 1.5818402 | 0 | 0 |
| Fibroblast | Cfh | 42.17324 | 1.4482436 | 0 | 0 |
| Fibroblast | Sparcl1 | 41.2387 | 1.5662547 | 0 | 0 |
| Fibroblast | Zbtb20 | 40.20208 | 1.0656797 | 0 | 0 |
| Fibroblast | Lama2 | 38.712154 | 2.8623848 | 0 | 0 |
| Fibroblast | Ptprd | 38.43185 | 1.355002 | 0 | 0 |

| | | | | | |
|---|---|---|---|---|---|
| Fibroblast | Sox5 | 36.815086 | 1.6919355 | 1.06E-296 | 1.38E-293 |
| Fibroblast | Pdgfra | 36.594883 | 2.7001147 | 3.45E-293 | 4.28E-290 |
| Fibroblast | Tshz2 | 35.832127 | 1.2855432 | 3.49E-281 | 3.97E-278 |
| Fibroblast | Csmd1 | 35.31029 | 1.8082458 | 4.08E-273 | 4.29E-270 |
| Fibroblast | Dcn | 34.657013 | 1.497509 | 3.50E-263 | 3.30E-260 |
| Fibroblast | Pbx1 | 34.414516 | 1.1583956 | 1.53E-259 | 1.39E-256 |
| Fibroblast | Rora | 32.30194 | 1.2114886 | 6.57E-229 | 4.98E-226 |
| Fibroblast | Rbms3 | 31.619278 | 2.1055849 | 2.01E-219 | 1.40E-216 |
| Fibroblast | Zeb1 | 29.71348 | 1.835921 | 5.14E-194 | 2.99E-191 |
| Fibroblast | Bicc1 | 29.51204 | 1.9625945 | 2.02E-191 | 1.15E-188 |
| Fibroblast | Xist | 27.826427 | 1.3226922 | 2.08E-170 | 1.11E-167 |
| Fibroblast | Auts2 | 27.175335 | 1.3273754 | 1.27E-162 | 6.55E-160 |
| Fibroblast | Plxdc2 | 26.89391 | 1.256711 | 2.59E-159 | 1.24E-156 |
| Fibroblast | Mgp | 26.78256 | 0.9558696 | 5.16E-158 | 2.39E-155 |
| Fibroblast | Cacnb2 | 26.758854 | 2.103646 | 9.74E-158 | 4.43E-155 |
| Fibroblast | Bnc2 | 26.68877 | 1.6073161 | 6.35E-157 | 2.84E-154 |
| Fibroblast | Sned1 | 25.442593 | 2.340673 | 8.53E-143 | 3.42E-140 |
| Fibroblast | Adamts19 | 25.2289 | 2.7546623 | 1.93E-140 | 7.64E-138 |
| Fibroblast | Dpp6 | 24.788013 | 2.84451 | 1.21E-135 | 4.58E-133 |
| Fibroblast | Ablim1 | 24.296211 | 1.3977396 | 2.15E-130 | 7.93E-128 |
| Fibroblast | 9530026P05Rik | 24.027248 | 1.8207654 | 1.44E-127 | 5.12E-125 |
| Fibroblast | Ogt | 23.572908 | 1.2802286 | 7.31E-123 | 2.49E-120 |
| Fibroblast | Ccdc141 | 23.260735 | 2.5771277 | 1.11E-119 | 3.73E-117 |
| Fibroblast | Pde7b | 23.22385 | 1.9027486 | 2.61E-119 | 8.71E-117 |
| Fibroblast | Zfpm2 | 22.67705 | 1.6398765 | 7.55E-114 | 2.34E-111 |
| Fibroblast | Rarres2 | 22.369724 | 1.0230047 | 7.76E-111 | 2.33E-108 |
| Fibroblast | Tcf4 | 22.344095 | 0.7467318 | 1.38E-110 | 4.09E-108 |
| Fibroblast | Tpm1 | 22.334986 | 0.6654011 | 1.69E-110 | 4.96E-108 |
| Fibroblast | Itih5 | 21.96111 | 1.558556 | 6.78E-107 | 1.93E-104 |
| Fibroblast | Sptbn1 | 21.960629 | 0.83363485 | 6.85E-107 | 1.93E-104 |
| Fibroblast | Nfia | 21.92795 | 0.9381369 | 1.41E-106 | 3.92E-104 |
| Fibroblast | Wt1 | 21.768421 | 2.0664618 | 4.62E-105 | 1.25E-102 |
| Fibroblast | Zfp36l1 | 21.748116 | 1.257886 | 7.20E-105 | 1.93E-102 |
| Fibroblast | Ptprk | 21.67653 | 2.1381364 | 3.42E-104 | 9.06E-102 |
| Fibroblast | Kcnt2 | 21.313385 | 1.900151 | 8.53E-101 | 2.16E-98 |
| Fibroblast | Pard3b | 21.074306 | 1.666265 | 1.37E-98 | 3.43E-96 |
| Fibroblast | Frem1 | 20.669897 | 3.1335683 | 6.47E-95 | 1.52E-92 |
| Fibroblast | Airn | 20.488855 | 1.5601522 | 2.71E-93 | 6.32E-91 |
| Fibroblast | Gas6 | 20.306507 | 0.5935277 | 1.13E-91 | 2.56E-89 |
| Fibroblast | Igfbp7 | 19.965578 | 0.58294505 | 1.10E-88 | 2.44E-86 |
| Fibroblast | Txnip | 19.947369 | 0.7620849 | 1.58E-88 | 3.48E-86 |
| Stromal - nonsteroidogenic | Aldh1a1 | 140.3374 | 1.5496122 | 0 | 0 |
| Stromal - nonsteroidogenic | mt-Rnr2 | 124.41688 | 0.64037085 | 0 | 0 |

| | | | | | |
|---|---|---|---|---|---|
| Stromal - nonsteroidogenic | Malat1 | 105.863884 | 0.4286431 | 0 | 0 |
| Stromal - nonsteroidogenic | Hsd3b1 | 104.61402 | 0.7132228 | 0 | 0 |
| Stromal - nonsteroidogenic | Akr1cl | 104.043076 | 0.5364099 | 0 | 0 |
| Stromal - nonsteroidogenic | Rn18s | 94.405136 | 0.6199027 | 0 | 0 |
| Stromal - nonsteroidogenic | mt-Rnr1 | 87.731674 | 0.7092463 | 0 | 0 |
| Stromal - nonsteroidogenic | Cyp11a1 | 78.02096 | 0.75642216 | 0 | 0 |
| Stromal - nonsteroidogenic | Mgarp | 72.80232 | 0.89900285 | 0 | 0 |
| Stromal - nonsteroidogenic | Gas6 | 63.692486 | 0.92267543 | 0 | 0 |
| Stromal - nonsteroidogenic | Dnajc15 | 62.04214 | 1.0890762 | 0 | 0 |
| Stromal - nonsteroidogenic | Dync1i1 | 60.06025 | 1.939754 | 0 | 0 |
| Stromal - nonsteroidogenic | Adh1 | 56.529274 | 1.1985763 | 0 | 0 |
| Stromal - nonsteroidogenic | Kcnd2 | 51.20437 | 1.5460876 | 0 | 0 |
| Stromal - nonsteroidogenic | Me1 | 46.96481 | 0.62446547 | 0 | 0 |
| Stromal - nonsteroidogenic | mt-Nd1 | 44.578693 | 0.14521728 | 0 | 0 |
| Stromal - nonsteroidogenic | Acsbg1 | 42.043003 | 0.60629547 | 0 | 0 |
| Stromal - nonsteroidogenic | Nckap5 | 38.970867 | 1.5605751 | 0 | 0 |
| Stromal - nonsteroidogenic | Mt1 | 38.29227 | 0.7276889 | 0 | 0 |
| Stromal - nonsteroidogenic | Abcb1b | 36.51504 | 0.94134563 | 6.40E-292 | 6.72E-290 |
| Stromal - nonsteroidogenic | Gstm1 | 36.32782 | 0.6272166 | 5.89E-289 | 6.09E-287 |
| Stromal - nonsteroidogenic | Cmss1 | 34.91047 | 0.46666458 | 5.16E-267 | 4.92E-265 |
| Stromal - nonsteroidogenic | Dlgap1 | 34.74501 | 1.4769312 | 1.65E-264 | 1.54E-262 |
| Stromal - nonsteroidogenic | Lars2 | 31.567123 | 0.53823006 | 1.04E-218 | 7.50E-217 |
| Stromal - nonsteroidogenic | Cped1 | 30.047855 | 0.68883806 | 2.33E-198 | 1.51E-196 |
| Stromal - nonsteroidogenic | mt-Nd4 | 28.664227 | 0.19290891 | 1.07E-180 | 6.19E-179 |
| Stromal - nonsteroidogenic | Coro2a | 28.402937 | 1.8689148 | 1.86E-177 | 1.05E-175 |
| Stromal - nonsteroidogenic | Neat1 | 27.312738 | 0.49070966 | 2.99E-164 | 1.51E-162 |
| Stromal - nonsteroidogenic | Mir6236 | 26.726871 | 0.6900505 | 2.29E-157 | 1.11E-155 |
| Stromal - nonsteroidogenic | Lsamp | 26.582191 | 1.7250082 | 1.09E-155 | 5.20E-154 |
| Stromal - nonsteroidogenic | Grm7 | 25.854052 | 1.916065 | 2.19E-147 | 9.73E-146 |
| Stromal - nonsteroidogenic | Tbx3 | 23.569586 | 1.1426493 | 7.91E-123 | 2.87E-121 |
| Stromal - nonsteroidogenic | A530020G20Rik | 23.56373 | 1.4606925 | 9.08E-123 | 3.29E-121 |
| Stromal - nonsteroidogenic | Gramd1b | 22.931673 | 0.52364653 | 2.25E-116 | 7.67E-115 |
| Stromal - nonsteroidogenic | Gm48099 | 21.061329 | 0.6021529 | 1.80E-98 | 5.18E-97 |
| Stromal - nonsteroidogenic | Gphn | 21.046333 | 0.50956076 | 2.47E-98 | 7.08E-97 |
| Stromal - nonsteroidogenic | Tcaf1 | 20.974373 | 0.50714296 | 1.12E-97 | 3.19E-96 |
| Stromal - nonsteroidogenic | Rerg | 20.85812 | 1.1025907 | 1.29E-96 | 3.64E-95 |
| Stromal - nonsteroidogenic | Camk1d | 20.593006 | 0.23173104 | 3.17E-94 | 8.67E-93 |
| Stromal - nonsteroidogenic | Cacnb2 | 20.406885 | 1.088911 | 1.45E-92 | 3.91E-91 |
| Stromal - nonsteroidogenic | Rbm47 | 19.68696 | 0.84986466 | 2.79E-86 | 7.03E-85 |
| Stromal - nonsteroidogenic | Hexb | 19.415009 | 0.4781316 | 5.76E-84 | 1.42E-82 |
| Stromal - nonsteroidogenic | Cacna1d | 19.366272 | 1.501786 | 1.49E-83 | 3.63E-82 |
| Stromal - nonsteroidogenic | Fdx1 | 18.631678 | 0.16498023 | 1.78E-77 | 4.06E-76 |
| Stromal - nonsteroidogenic | Ephx1 | 18.54373 | 0.7757365 | 9.16E-77 | 2.06E-75 |

| | | | | | |
|---|---|---|---|---|---|
| Stromal - nonsteroidogenic | Ppm1e | 18.540195 | 1.1539676 | 9.79E-77 | 2.20E-75 |
| Stromal - nonsteroidogenic | Snhg11 | 18.506372 | 1.8468436 | 1.83E-76 | 4.11E-75 |
| Stromal - nonsteroidogenic | Mt2 | 18.132505 | 0.66336435 | 1.77E-73 | 3.82E-72 |
| Stromal - nonsteroidogenic | mt-Cytb | 17.717808 | -0.0266183 | 3.06E-70 | 6.30E-69 |
| Stromal - nonsteroidogenic | Pcsk5 | 17.37637 | 0.7019813 | 1.25E-67 | 2.49E-66 |
| Stromal - steroidogenic | Aldh1a1 | 271.48846 | 3.5628538 | 0 | 0 |
| Stromal - steroidogenic | Hsd3b1 | 239.5961 | 2.3376887 | 0 | 0 |
| Stromal - steroidogenic | Cyp11a1 | 193.94449 | 2.2614405 | 0 | 0 |
| Stromal - steroidogenic | Mgarp | 186.23442 | 2.6088464 | 0 | 0 |
| Stromal - steroidogenic | Akr1cl | 172.49146 | 1.7025665 | 0 | 0 |
| Stromal - steroidogenic | Adh1 | 155.45901 | 3.14048 | 0 | 0 |
| Stromal - steroidogenic | mt-Nd1 | 150.00072 | 1.554775 | 0 | 0 |
| Stromal - steroidogenic | Gas6 | 146.90208 | 2.2730477 | 0 | 0 |
| Stromal - steroidogenic | Me1 | 146.43889 | 2.1348605 | 0 | 0 |
| Stromal - steroidogenic | mt-Nd4 | 142.52731 | 1.7362952 | 0 | 0 |
| Stromal - steroidogenic | Acsbg1 | 134.47955 | 2.0138664 | 0 | 0 |
| Stromal - steroidogenic | Dnajc15 | 133.6498 | 2.314094 | 0 | 0 |
| Stromal - steroidogenic | Gstm1 | 126.22605 | 2.085989 | 0 | 0 |
| Stromal - steroidogenic | mt-Cytb | 125.99438 | 1.4222904 | 0 | 0 |
| Stromal - steroidogenic | Fdx1 | 120.59125 | 1.6354501 | 0 | 0 |
| Stromal - steroidogenic | Tcaf1 | 112.241356 | 2.1609502 | 0 | 0 |
| Stromal - steroidogenic | Mt1 | 105.38594 | 1.8861451 | 0 | 0 |
| Stromal - steroidogenic | Cpe | 104.218704 | 1.4474863 | 0 | 0 |
| Stromal - steroidogenic | mt-Nd2 | 102.15865 | 1.6732209 | 0 | 0 |
| Stromal - steroidogenic | Fth1 | 95.646286 | 1.0358169 | 0 | 0 |
| Stromal - steroidogenic | Serinc3 | 94.02164 | 1.4111005 | 0 | 0 |
| Stromal - steroidogenic | Prss35 | 93.82107 | 1.8827006 | 0 | 0 |
| Stromal - steroidogenic | Gstm2 | 92.068756 | 1.8765117 | 0 | 0 |
| Stromal - steroidogenic | Pank1 | 91.030106 | 2.0533948 | 0 | 0 |
| Stromal - steroidogenic | Tmem176b | 81.53556 | 1.6582894 | 0 | 0 |
| Stromal - steroidogenic | Mgst1 | 80.57467 | 1.4085798 | 0 | 0 |
| Stromal - steroidogenic | Hmgcs2 | 78.23772 | 1.556036 | 0 | 0 |
| Stromal - steroidogenic | Abcb1b | 76.24983 | 1.8492812 | 0 | 0 |
| Stromal - steroidogenic | Hao2 | 74.32883 | 2.0616918 | 0 | 0 |
| Stromal - steroidogenic | Sqstm1 | 71.70652 | 1.1615993 | 0 | 0 |
| Stromal - steroidogenic | Ephx1 | 71.48805 | 2.5058122 | 0 | 0 |
| Stromal - steroidogenic | Tmem176a | 69.70066 | 1.6867398 | 0 | 0 |
| Stromal - steroidogenic | Gramd1b | 67.99662 | 1.3187337 | 0 | 0 |
| Stromal - steroidogenic | Dbi | 67.934204 | 1.2232742 | 0 | 0 |
| Stromal - steroidogenic | Smarca1 | 67.46701 | 1.3896446 | 0 | 0 |
| Stromal - steroidogenic | Star | 66.63807 | 1.1419593 | 0 | 0 |
| Stromal - steroidogenic | Map1lc3a | 63.766083 | 1.3574257 | 0 | 0 |
| Stromal - steroidogenic | Dhrs7 | 63.464764 | 1.9794605 | 0 | 0 |

| | | | | | |
|---|---|---|---|---|---|
| Stromal - steroidogenic | mt-Rnr2 | 63.131603 | 0.520502 | 0 | 0 |
| Stromal - steroidogenic | Mt2 | 61.354668 | 1.8203807 | 0 | 0 |
| Stromal - steroidogenic | Pcolce | 61.06586 | 1.5655198 | 0 | 0 |
| Stromal - steroidogenic | Prlr | 60.812458 | 1.0416623 | 0 | 0 |
| Stromal - steroidogenic | mt-Nd5 | 59.9259 | 1.3270347 | 0 | 0 |
| Stromal - steroidogenic | Atp5g3 | 59.25851 | 0.8594829 | 0 | 0 |
| Stromal - steroidogenic | Ptp4a2 | 57.513214 | 0.9247502 | 0 | 0 |
| Stromal - steroidogenic | Prxl2a | 57.228756 | 2.223332 | 0 | 0 |
| Stromal - steroidogenic | Fxyd1 | 56.315746 | 1.8634471 | 0 | 0 |
| Stromal - steroidogenic | Fads2 | 55.930805 | 1.5693986 | 0 | 0 |
| Stromal - steroidogenic | Fdxr | 55.779175 | 1.1708719 | 0 | 0 |
| Stromal - steroidogenic | Tmem86a | 55.2797 | 1.2767742 | 0 | 0 |
| Adipocyte | Fabp4 | 47.123272 | 5.2728643 | 0 | 0 |
| Adipocyte | C3 | 45.23833 | 5.476862 | 0 | 0 |
| Adipocyte | Car3 | 44.918682 | 7.1059775 | 0 | 0 |
| Adipocyte | Cfd | 41.5505 | 7.296378 | 0 | 0 |
| Adipocyte | Dcn | 36.996 | 2.760905 | 1.33E-299 | 6.04E-296 |
| Adipocyte | Gsn | 32.294353 | 3.4751 | 8.40E-229 | 2.55E-225 |
| Adipocyte | Tmsb4x | 27.318598 | 1.2141579 | 2.55E-164 | 5.36E-161 |
| Adipocyte | Dpt | 22.279335 | 4.4788713 | 5.86E-110 | 6.67E-107 |
| Adipocyte | mt-Rnr2 | 21.737265 | 0.31146607 | 9.12E-105 | 8.30E-102 |
| Adipocyte | Col1a2 | 15.38708 | 1.200266 | 2.00E-53 | 6.82E-51 |
| Adipocyte | Plac9a | 15.331216 | 3.237062 | 4.73E-53 | 1.59E-50 |
| Adipocyte | Mgp | 12.9279585 | 0.98792505 | 3.13E-38 | 5.77E-36 |
| Adipocyte | Col3a1 | 11.846567 | 0.99083006 | 2.24E-32 | 3.11E-30 |
| Adipocyte | Igfbp4 | 11.516046 | 1.115048 | 1.10E-30 | 1.35E-28 |
| Adipocyte | Serping1 | 11.513063 | 2.2127998 | 1.13E-30 | 1.39E-28 |
| Adipocyte | Igfbp6 | 11.4111 | 1.8093334 | 3.68E-30 | 4.41E-28 |
| Adipocyte | Ifi27l2a | 10.643711 | 2.646647 | 1.87E-26 | 1.78E-24 |
| Adipocyte | Adipoq | 10.422298 | 7.208054 | 1.96E-25 | 1.77E-23 |
| Adipocyte | Lpl | 9.668352 | 2.1714013 | 4.11E-22 | 2.98E-20 |
| Adipocyte | Tagln | 9.659389 | 1.1022464 | 4.49E-22 | 3.25E-20 |
| Adipocyte | Apoe | 9.350609 | 0.55775905 | 8.71E-21 | 5.76E-19 |
| Adipocyte | Retn | 9.051332 | 6.2795086 | 1.41E-19 | 8.72E-18 |
| Adipocyte | Tpm2 | 8.746252 | 1.049061 | 2.21E-18 | 1.23E-16 |
| Adipocyte | Trf | 8.727027 | 1.4746932 | 2.61E-18 | 1.45E-16 |
| Adipocyte | Ifitm3 | 8.286905 | 1.0107193 | 1.16E-16 | 5.67E-15 |
| Adipocyte | Efemp1 | 8.230559 | 1.9004503 | 1.86E-16 | 8.93E-15 |
| Adipocyte | Ccdc80 | 8.229879 | 2.2061052 | 1.87E-16 | 8.96E-15 |
| Adipocyte | Igkc | 8.195513 | 2.9019916 | 2.50E-16 | 1.18E-14 |
| Adipocyte | H2-Ab1 | 8.035316 | 1.3035195 | 9.33E-16 | 4.25E-14 |
| Adipocyte | Cidec | 7.929874 | 7.12181 | 2.19E-15 | 9.61E-14 |
| Adipocyte | Lum | 7.801228 | 1.9312426 | 6.13E-15 | 2.58E-13 |

| | | | | | |
|---|---|---|---|---|---|
| Adipocyte | Hp | 7.6758823 | 4.040653 | 1.64E-14 | 6.66E-13 |
| Adipocyte | Igfbp7 | 6.6796346 | 0.40733767 | 2.40E-11 | 7.12E-10 |
| Adipocyte | Pck1 | 6.5855193 | 7.332969 | 4.53E-11 | 1.31E-09 |
| Adipocyte | Igfbp3 | 6.356131 | 2.9704149 | 2.07E-10 | 5.51E-09 |
| Adipocyte | C4b | 6.312701 | 3.2016253 | 2.74E-10 | 7.16E-09 |
| Adipocyte | Cdo1 | 6.2969246 | 2.7851288 | 3.04E-10 | 7.89E-09 |
| Adipocyte | Rarres2 | 6.0602627 | 0.7302251 | 1.36E-09 | 3.25E-08 |
| Adipocyte | Lsp1 | 5.897734 | 2.2479088 | 3.69E-09 | 8.43E-08 |
| Adipocyte | H2-Eb1 | 5.80677 | 1.2562672 | 6.37E-09 | 1.42E-07 |
| Adipocyte | Fth1 | 5.525892 | -0.1049803 | 3.28E-08 | 6.62E-07 |
| Adipocyte | C1s1 | 5.5028887 | 0.9335777 | 3.74E-08 | 7.46E-07 |
| Adipocyte | Gpc3 | 5.4235487 | 2.0587864 | 5.84E-08 | 1.15E-06 |
| Adipocyte | Penk | 5.3881803 | 2.6423528 | 7.12E-08 | 1.38E-06 |
| Adipocyte | Galnt15 | 5.298632 | 1.7635528 | 1.17E-07 | 2.19E-06 |
| Adipocyte | Spon1 | 5.2379684 | 2.0948427 | 1.62E-07 | 2.97E-06 |
| Adipocyte | Cyp2e1 | 4.9832497 | 5.5859513 | 6.25E-07 | 1.05E-05 |
| Adipocyte | Ighm | 4.955585 | 3.459496 | 7.21E-07 | 1.20E-05 |
| Adipocyte | Celf2 | 4.897533 | 1.2830071 | 9.70E-07 | 1.59E-05 |
| Adipocyte | Ghr | 4.8607597 | 1.0936961 | 1.17E-06 | 1.88E-05 |
