## Extended Data Table 4 for "Aging disrupts spatiotemporal coordination in the cycling murine ovary"

**Spot counts - All cell types**

(Fig. 1b,c; ED Fig. 3b, 3c, 3d, 5b, 5c, 5e)

| Cell type | *n* |
|---|---|
| Adipocyte | 6105 |
| Cumulus - oocyte | 10053 |
| Endothelial - blood | 14405 |
| Endothelial - lymphatic | 15099 |
| Endothelial - vascular | 27221 |
| Epithelial - oviduct | 4439 |
| Epithelial - surface | 12681 |
| Fibroblast | 20456 |
| GC - antral early mitotic | 8861 |
| GC - antral luteinizing | 26169 |
| GC - antral mitotic | 44900 |
| GC - atretic | 17538 |
| GC - preantral | 7922 |
| Immune | 41879 |
| Luteal - Akr1c18 | 60102 |
| Luteal - Parm1 | 45812 |
| Muscle | 48294 |
| Oocyte | 14107 |
| Stromal - nonsteroidogenic | 106442 |
| Stromal - steroidogenic | 65694 |
| Theca | 9695 |

**Puck counts by age**

(Fig. 1d, 5b, 5g, 7b, 7f, ED Fig. 1i, 6d)

| Age | *n* |
|---|---|
| Young | 32 |
| Middle | 16 |
| Old | 21 |

**Segment type counts - All ages**

(Fig. 1i, 3b, 3g, 4a, 5c, 6b, ED Fig. 1g, 5d)

| Segment type | *n* |
|---|---|
| Follicle - preantral | 238 |
| Follicle - antral mitotic | 191 |
| Follicle - antral luteinizing | 43 |
| Follicle - atretic | 181 |
| CL - Parm1 | 87 |
| CL - Akr1c18 | 147 |

**Oocyte counts by enclosing follicle type - All ages**

(Fig. 2a, 2e, ED Fig. 2b)

| **Enclosing follicle type** | ***n*** |
|---|---|
| Primordial | *57* |
| Follicle - preantral | 148 |
| Follicle - antral mitotic | 74 |
| Follicle - antral luteinizing | 75 |
| Follicle - atretic | 4 |

**Oocyte counts by age**

(Fig. 2f)

| **Age** | ***n*** |
|---|---|
| Young | 229 |
| Middle | 89 |
| Old | 40 |

**Non-primoridal oocyte counts by age**

(Fig. 2g, ED Fig. 2d)

| **Age** | ***n*** |
|---|---|
| Young | 198 |
| Middle | 72 |
| Old | 31 |

**Oocyte counts by enclosing follicle type and age**

(ED Fig. 2e, 2f)

| **Age** | **Enclosing follicle type** | ***n*** |
|---|---|---|
| Young | Follicle - preantral | 97 |
| Young | Follicle - antral mitotic | 38 |
| Young | Follicle - antral luteinizing | 3 |
| Young | Follicle - atretic | 50 |
| Old | Follicle - preantral | 9 |
| Old | Follicle - antral mitotic | 17 |
| Old | Follicle - antral luteinizing | 1 |
| Old | Follicle - atretic | 4 |

**Atretic follicle counts**

(Fig. 3d, ED Fig. 3g)

| **Follicle type** | ***n*** |
|---|---|
| Atretic | 188 |
| Non-atretic | 480 |

**Segment type counts by age and stage**

(Fig. 3i, 4i, 4j, 5e ED Fig. 3f, 4d, 4e, 5e, 5f)

| **Age** | **Stage** | **Segment type** | ***n*** |
|---|---|---|---|
| Young | Estrus | Follicle - preantral | 138 |
| Young | Estrus | Follicle - antral mitotic | 84 |

| | | | |
|---|---|---|---|
| Young | Estrus | Follicle - antral luteinizing | 6 |
| Young | Estrus | Follicle - atretic | 88 |
| Young | Estrus | CL - Parm1 | 56 |
| Young | Estrus | CL - Akr1c18 | 36 |
| Young | Metestrus | Follicle - preantral | 8 |
| Young | Metestrus | Follicle - antral mitotic | 23 |
| Young | Metestrus | Follicle - antral luteinizing | 7 |
| Young | Metestrus | Follicle - atretic | 28 |
| Young | Metestrus | CL - Parm1 | 0 |
| Young | Metestrus | CL - Akr1c18 | 15 |
| Middle | Estrus | Follicle - preantral | 12 |
| Middle | Estrus | Follicle - antral mitotic | 16 |
| Middle | Estrus | Follicle - antral luteinizing | 4 |
| Middle | Estrus | Follicle - atretic | 6 |
| Middle | Estrus | CL - Parm1 | 2 |
| Middle | Estrus | CL - Akr1c18 | 14 |
| Middle | Metestrus | Follicle - preantral | 55 |
| Middle | Metestrus | Follicle - antral mitotic | 33 |
| Middle | Metestrus | Follicle - antral luteinizing | 11 |
| Middle | Metestrus | Follicle - atretic | 43 |
| Middle | Metestrus | CL - Parm1 | 2 |
| Middle | Metestrus | CL - Akr1c18 | 27 |
| Old | Estrus | preantral | 18 |
| Old | Estrus | antral mitotic | 31 |
| Old | Estrus | antral luteinizing | 10 |
| Old | Estrus | atretic | 13 |
| Old | Estrus | CL - Parm1 | 16 |
| Old | Estrus | CL - Akr1c18 | 35 |
| Old | Metestrus | preantral | 9 |
| Old | Metestrus | antral mitotic | 10 |
| Old | Metestrus | antral luteinizing | 5 |
| Old | Metestrus | atretic | 10 |
| Old | Metestrus | CL - Parm1 | 11 |
| Old | Metestrus | CL - Akr1c18 | 22 |

**Spot counts - Atretic subclusters**

(Fig. 3j, 3l; ED Fig. 3i)

| **Cell type** | ***n*** |
|---|---|
| GC atretic 1 | 9062 |
| GC atretic 2 | 4185 |
| GC atretic 3 | 3928 |

**Atretic follicle type counts by age and stage**

(Fig. 3m)

| Age | Atresia status | Follicle type | *n* |
|---|---|---|---|
| Young | Non-atretic | preantral | 146 |
| Young | Non-atretic | antral mitotic | 107 |
| Young | Non-atretic | antral luteinizing | 13 |
| Young | Atretic | preantral | 18 |
| Young | Atretic | antral | 82 |
| Old | Non-atretic | preantral | 14 |
| Old | Non-atretic | antral mitotic | 25 |
| Old | Non-atretic | antral luteinizing | 9 |
| Old | Atretic | preantral | 10 |
| Old | Atretic | antral | 11 |

**Coarse follicle type by age**

(Fig. 4f, 7h, ED Fig. 4c)

| Age | Follicle type | *n* |
|---|---|---|
| Young | preantral | 144 |
| Young | antral | 120 |
| Young | atretic | 116 |
| Old | preantral | 27 |
| Old | antral | 56 |
| Old | atretic | 23 |

**Pucks counts by age and stage**

(Fig. 4g, 4j, 6b)

| Age | Stage | *n* |
|---|---|---|
| Young | Estrus | 26 |
| Young | Metestrus | 6 |
| Middle | Estrus | 5 |
| Middle | Metestrus | 11 |
| Old | Estrus | 13 |
| Old | Metestrus | 8 |

**Total follicle counts by age and stage**

(Fig. 4h, 4k, ED Fig. 4g)

| Age | Stage | *n* |
|---|---|---|
| Young | Estrus | 281 |
| Young | Metestrus | 66 |
| Old | Estrus | 58 |
| Old | Metestrus | 23 |

**Segment type counts by age**

(Fig.5f, 6d, ED Fig. 4a)

| Age | Segment type | *n* |
|---|---|---|
| Young | preantral | 146 |

| | | |
|---|---|---|
| Young | antral mitotic | 107 |
| Young | antral luteinizing | 13 |
| Young | atretic | 116 |
| Young | CL - Parm1 | 56 |
| Young | CL - Akr1c18 | 51 |
| Middle | preantral | 67 |
| Middle | antral mitotic | 49 |
| Middle | antral luteinizing | 15 |
| Middle | atretic | 49 |
| Middle | CL - Parm1 | 4 |
| Middle | CL - Akr1c18 | 41 |
| Old | preantral | 27 |
| Old | antral mitotic | 41 |
| Old | antral luteinizing | 15 |
| Old | atretic | 23 |
| Old | CL - Parm1 | 27 |
| Old | CL - Akr1c18 | 57 |

**Spot counts - Immune subcluster**

(Fig. 6a)

| **Cell type** | ***n*** |
|---|---|
| Immune - Macrophage 1 | 16585 |
| Immune - Macrophage 2 | 12148 |
| Immune - Macrophage 3 | 4655 |
| Immune - Macrophage 4 | 4571 |
| Immune - Dendritic cell | 2455 |
| Immune - Mast cell | 383 |
| Immune - Plasma cell | 629 |
| Immune - T cell | 453 |

**Spot counts - Stromal Immune/MNGC**

(Fig. 7e, ED Fig. 6c)

| **Age** | **MNGC Status/Localization** | ***n*** |
|---|---|---|
| Young | MNGC-low | 3,746 |
| Old | MNGC-low | 147 |
| Old | MNGC-high - Inside | 5,356 |
| Old | MNGC-high - Outside | 1,706 |

**Spot counts - Stromal**

(Fig. 7g, ED Fig. 6f)

| **Age** | ***n*** |
|---|---|
| Young | 33,666 |
| Middle | 17,188 |
| Old | 14,263 |
