## Extended Data Table 5 for "Aging disrupts spatiotemporal coordination in the cycling murine ovary"

| Gene | logfoldchange | pvalue_adj |
|---|---|---|
| Nlrp9b | 1.6326588 | 2.2992145235817063e-11 |
| Usp31 | 3.0865993 | 1.591868779578414e-10 |
| Tent4b | 1.8915353 | 2.4563910448081304e-10 |
| Osbpl11 | 1.5526497 | 2.6035339530215513e-10 |
| Rb1 | 2.1765637 | 5.577267726079616e-10 |
| Zfp979 | 1.753542 | 1.8052459700516551e-09 |
| Ide | 1.5089597 | 2.537600444139861e-08 |
| Ildr1 | 2.6328905 | 1.0208293498325488e-07 |
| Ctdp1 | 3.110516 | 1.7695578897935515e-07 |
| Gda | 1.1997943 | 1.0608020139593285e-07 |
| 1700034K08F | 3.72153 | 2.2563148056446756e-07 |
| Pde6b | 3.4584453 | 3.061757105354593e-07 |
| Atg16l1 | 1.3270646 | 1.8879895255075912e-07 |
| Txndc17 | 1.7335577 | 2.3010152748704557e-07 |
| Yipf4 | 1.8527143 | 2.934811447786352e-07 |
| Setd2 | 1.2059202 | 2.3010152748704557e-07 |
| Stag1 | 2.1652856 | 4.7572833675002807e-07 |
| Rbm33 | 1.270077 | 4.764850871567641e-07 |
| Phactr3 | 1.4924022 | 4.7572833675002807e-07 |
| Cyp39a1 | 2.3912468 | 6.610425352863848e-07 |
| Prpf39 | 3.0499375 | 1.0610152545260644e-06 |
| Ivns1abp | 1.3369948 | 5.952565919995774e-07 |
| Kbtbd3 | 2.5024903 | 1.2275897854609564e-06 |
| Odad1 | 25.977938 | 1.459182426807243e-06 |
| Sidt2 | 2.2361615 | 1.2275897854609564e-06 |
| Rpl21 | 4.2385983 | 1.960538229264304e-06 |
| 9830004L10F | 3.942308 | 2.177259558463411e-06 |
| Prss35 | 3.2273636 | 2.177259558463411e-06 |
| Ankrd50 | 2.0122974 | 2.058382404397213e-06 |
| Lzts1 | 1.6001973 | 1.9497297542396962e-06 |
| Bend5 | 1.4890532 | 2.177259558463411e-06 |
| Hsd17b10 | 1.7192217 | 2.4410859032639948e-06 |
| Stk24 | 1.2925166 | 2.4410859032639948e-06 |
| 4930566F21F | 2.6179156 | 3.285565287789486e-06 |
| D630045J12F | 1.7734804 | 2.8236711149549176e-06 |
| Arid4a | 1.1540011 | 2.5132290193686006e-06 |
| Ccp110 | 1.4926679 | 2.9786105173620525e-06 |
| Ccdc71l | 7.044669 | 5.6749849964875785e-06 |
| Riox2 | 1.4338341 | 3.92749472269737e-06 |
| Tmem64 | 3.7320032 | 6.1287518709500124e-06 |

| | | |
|---|---|---|
| Gm44087 | 26.289467 | 6.285304785873317e-06 |
| Fam20b | 2.1470902 | 5.6749849964875785e-06 |
| Sh3bp4 | 2.884049 | 6.286775510866932e-06 |
| Oxr1 | 2.448643 | 6.1710462136540065e-06 |
| Lysmd4 | 1.243524 | 5.6749849964875785e-06 |
| Rpl7a | 3.0717509 | 7.676913131372797e-06 |
| Zbtb33 | 1.8347303 | 7.289040271335674e-06 |
| Efnb2 | 1.1976204 | 6.3534329470448955e-06 |
| Kcnk13 | 1.607349 | 7.289040271335674e-06 |
| Sptlc2 | 1.4607533 | 7.838481469839551e-06 |
| Ppp3ca | 1.9427981 | 7.947216142709467e-06 |
| Incenp | 1.0031799 | 7.217307261034849e-06 |
| Usp1 | 1.0618618 | 7.289040271335674e-06 |
| Srsf9 | 1.2652193 | 7.289040271335674e-06 |
| Akr7a5 | 1.4038998 | 9.854188469572701e-06 |
| Calm3 | 1.4310203 | 1.0159250482654221e-05 |
| N4bp2 | 0.99919593 | 9.755037449789704e-06 |
| Tnpo1 | 1.0932454 | 1.0159250482654221e-05 |
| Twnk | 1.6682585 | 1.3078065689012658e-05 |
| Lamc1 | 1.9425603 | 1.3654953662621539e-05 |
| Ifitm3 | 1.4842172 | 1.1956270818121838e-05 |
| Cib1 | 1.6726261 | 1.3078065689012658e-05 |
| Erbin | 1.6597642 | 1.3234573029357222e-05 |
| Cdc42bpg | 4.315692 | 1.7576854684539277e-05 |
| Septin2 | 1.21873 | 1.3698560436032651e-05 |
| Dync2h1 | 2.1668782 | 1.6101625632956822e-05 |
| Tpx2 | 0.9653753 | 1.3078065689012658e-05 |
| Fbxo40 | 1.3432033 | 1.4122354338851713e-05 |
| Rps4x | 1.8326979 | 1.5770298070406485e-05 |
| Myh14 | 4.809172 | 1.835152490918034e-05 |
| Dse | 1.7544056 | 1.5225087582588408e-05 |
| Mageh1 | 1.3845596 | 1.5421356202845035e-05 |
| Nubp2 | 1.2937483 | 1.5462073385030558e-05 |
| Bach1 | 1.9065526 | 1.809296828006179e-05 |
| Kmt2b | 1.0448323 | 1.5588793560428565e-05 |
| Slf2 | 1.1701312 | 1.5588793560428565e-05 |
| Zkscan8 | 4.2020335 | 2.1418477146483393e-05 |
| Polr3k | 1.1706975 | 1.651260046256339e-05 |
| Il17f | 2.4699647 | 1.948605070178344e-05 |
| Tab2 | 1.265528 | 1.6987359080721448e-05 |
| Gstm1 | 1.3640403 | 1.6987359080721448e-05 |

| | | |
|---|---|---|
| Cep85l | 1.3476202 | 1.8416850540387257e-05 |
| Fam91a1 | 2.092557 | 2.2652910384930652e-05 |
| Dcbld1 | 1.6014906 | 2.0022725921177935e-05 |
| B020011L13F | 6.10766 | 2.636504589455389e-05 |
| Cacna1i | 3.1021914 | 2.6070226510260566e-05 |
| Mgat1 | 1.792379 | 2.105762062486324e-05 |
| Kdsr | 2.4475865 | 2.4670137673985445e-05 |
| Phc3 | 3.2609272 | 2.720014579201658e-05 |
| Lsm11 | 1.5458788 | 2.2253641531758487e-05 |
| Ltc4s | 1.4672467 | 2.189281920249707e-05 |
| Trim15 | 6.9901185 | 2.9703162715847916e-05 |
| Tmem151a | 4.1352363 | 2.9703162715847916e-05 |
| Arid1a | 1.0168183 | 2.189281920249707e-05 |
| Gpd2 | 3.3835464 | 2.9703162715847916e-05 |
| Rcc2 | 1.0484678 | 2.2724353210332206e-05 |
| Gm21411 | 2.8977218 | 3.002124766521159e-05 |
| Plpp3 | 0.95168984 | 2.632812857015929e-05 |
| Pcsk6 | 2.4704115 | 3.330218734679347e-05 |
