## Extended Data Table 6 for "Aging disrupts spatiotemporal coordination in the cycling murine ovary"

| Gene | logfoldchange | pvalue_adj |
|---|---|---|
| Prss35 | 2.390709 | 0 |
| Me1 | 2.2283835 | 0 |
| Ybx1 | 1.8812891 | 0 |
| Star | 1.8724681 | 0 |
| Mgarp | 1.7364048 | 0 |
| Sparc | 1.484137 | 0 |
| Col3a1 | 1.546786 | 0 |
| Cyp11a1 | 2.4814904 | 0 |
| Aldh1a1 | 2.222102 | 0 |
| Sbsn | 2.463564 | 0 |
| Pank1 | 1.3369651 | 0 |
| Hmgcs1 | 1.43618 | 0 |
| Dnajc15 | 1.0479804 | 0 |
| Lgals1 | 1.0386517 | 0 |
| Fdx1 | 1.0775607 | 0 |
| Hao2 | 1.2667428 | 0 |
| Tpt1 | 3.1642096 | 0 |
| Pcolce | 1.2165914 | 0 |
| Trf | 1.9787121 | 0 |
| Glul | 1.4216361 | 0 |
| Map1lc3a | 1.0281702 | 0 |
| Rps12 | 2.4788392 | 0 |
| Abcb1b | 1.2611561 | 0 |
| Vim | 1.0381731 | 0 |
| Hsd3b1 | 2.1685362 | 0 |
| Rps6 | 3.3795094 | 0 |
| Eef1a1 | 0.9740602 | 0 |
| Serinc3 | 0.85069454 | 0 |
| Ifi27 | 0.97188616 | 0 |
| Tspo | 1.3412713 | 0 |
| Rps17 | 2.7680697 | 0 |
| Dbi | 0.83386815 | 0 |
| Igfbp2 | 1.6082916 | 0 |
| Rpl11 | 1.7734764 | 0 |
| Rpl21 | 2.8298786 | 0 |
| Serpinb6a | 0.93299496 | 0 |
| Rps27a | 2.8848574 | 0 |
| Rpl17 | 2.5491307 | 0 |
| Fdxr | 0.8348511 | 0 |
| Rpl9 | 2.4329515 | 0 |

| | | |
|---|---|---|
| S100a10 | 0.8734504 | 0 |
| Wnt6 | 1.7467856 | 0 |
| Ggh | 1.1307552 | 0 |
| Gamt | 1.2447021 | 0 |
| Fau | 2.4934518 | 0 |
| Gpx1 | 0.8609339 | 0 |
| Rpl5 | 2.0695462 | 0 |
| Rpl31 | 1.506652 | 0 |
| Rps16 | 2.1311033 | 0 |
| Ndufa4 | 0.8869995 | 0 |
| Taldo1 | 0.9041029 | 0 |
| Rpl7a | 2.2924178 | 0 |
| Chchd2 | 0.68457526 | 0 |
| Prkar2b | 0.96294796 | 0 |
| Tceal9 | 0.6554572 | 0 |
| Hsp90ab1 | 0.6652516 | 0 |
| Rpl23a | 2.7113068 | 0 |
| Trp53inp2 | 1.2146889 | 0 |
| Dhcr24 | 1.0972068 | 0 |
| Tmem159 | 1.0623591 | 0 |
| Rpsa | 1.3045604 | 0 |
| Ppia | 1.738214 | 0 |
| Rpl12 | 1.6103235 | 0 |
| Rpl28 | 2.7024508 | 0 |
| Rps23 | 2.2959294 | 0 |
| Serpinb1a | 1.121543 | 0 |
| Il31ra | 2.0454302 | 0 |
| Tle5 | 0.7598912 | 0 |
| Ccnd2 | 1.0386174 | 0 |
| Rps7 | 2.0602489 | 0 |
| Cited2 | 0.849071 | 0 |
| Rpl36a | 2.8411255 | 0 |
| Atp5b | 0.6723521 | 0 |
| Rpl19 | 1.7951523 | 0 |
| Rpl41 | 0.83756816 | 3.390998305515415e-306 |
| Rps2 | 0.68933874 | 3.4068281424279013e-305 |
| Hspd1 | 0.6931458 | 5.383801653395963e-304 |
| Cox6a1 | 0.6119932 | 2.3147784494186996e-295 |
| Mdh1 | 0.73443305 | 7.333916927197615e-288 |
| Cox6b1 | 0.661399 | 6.735040816337942e-283 |
| Tsc22d1 | 0.57945025 | 8.440764367625054e-282 |

| | | |
|---|---|---|
| Maoa | 1.1791325 | 3.808927125039666e-278 |
| Pebp1 | 0.75491595 | 1.0446346024198514e-274 |
| Dmpk | 1.438762 | 2.0429571518299873e-265 |
| mt-Nd1 | 0.75355715 | 1.622631282967427e-261 |
| Atpif1 | 0.57119554 | 1.4074292775083287e-258 |
| Prdx3 | 0.749295 | 1.2035599299779295e-257 |
| Rps25 | 1.507558 | 2.3676799476402636e-255 |
| Uqcrb | 0.6925844 | 1.8247176530761928e-255 |
| Gpx3 | 0.82167655 | 3.7529532141391314e-253 |
| Rps4x | 1.1076816 | 4.2880641653693657e-250 |
| mt-Nd2 | 0.55103636 | 2.8114538928750132e-248 |
| Smarca1 | 0.633673 | 2.602269280018853e-246 |
| Rpl27 | 1.9174075 | 2.626276605162699e-243 |
| Ubb | 0.539993 | 2.333897943640973e-242 |
| Atp5a1 | 0.5713355 | 5.10406191138449e-239 |
| Timp1 | 1.243419 | 3.773765304480297e-238 |
| Rpl10 | 2.2744002 | 6.3862865770387656e-232 |
| Rps27 | 2.187134 | 1.7927503063287005e-228 |
